# Entropy sorting of single cell RNA sequencing data reveals the inner cell mass in the human pre-implantation embryo

**DOI:** 10.1101/2022.04.08.487653

**Authors:** Arthur Radley, Elena Corujo-Simon, Jennifer Nichols, Austin Smith, Sara-Jane Dunn

## Abstract

A major challenge in single cell gene expression analysis is to discern meaningful cellular heterogeneity from technical or biological noise. To address this challenge, we present Entropy Sorting, a mathematical framework that distinguishes genes indicative of cell identity. ES achieves this in an unsupervised manner by quantifying if observed correlations between features are more likely to have occurred due to random chance versus a dependent relationship, without the need for any user defined significance threshold. On synthetic data we demonstrate the removal of noisy signals to reveal a higher resolution of gene expression patterns than commonly used feature selection methods. We then apply ES to human pre-implantation embryo scRNA-seq data. Previous studies failed to unambiguously identify early inner cell mass (ICM), suggesting that the human embryo may diverge from the mouse paradigm. In contrast, ES resolves the ICM and reveals sequential lineage bifurcations as in the classical model. Entropy sorting thus provides a powerful approach for maximising information extraction from high dimensional datasets such as scRNA-seq data.

## 1 INTRODUCTION

Single cell RNA sequencing (scRNA-seq) (Tang et al. 2009) is a powerful technique for studying cell identity and heterogeneity by capturing transcriptome-wide RNA expression at single cell resolution. As such, scRNA-seq yields an unbiased dataset, rather than being restricted to a pre-defined subset of genes of interest. However, the cost of this information-rich data is a practical limitation known as the curse of dimensionality (CoD) (Bellman 1967). This phenomena arises when analysing datasets with increasingly large dimensions - the number of features or variables. In the context of scRNA-seq, we typically refer to each gene as a feature, and each cell as sample. As the number of features increases, our ability to discern patterns between samples and/or features decreases (Altman and Krzywinski 2018). Thus, by sequencing tens of thousands of genes, we reduce our ability to identify differential gene expression patterns. The challenge is exacerbated by technical artefacts introduced during data collection, such as batch effects and false negative drop outs (Kiselev, Andrews, and Hemberg 2019), which further weaken the correlations between cells and genes.

The antidote to this challenge is the blessing of dimensionality (Zimek, Schubert, and Kriegel 2012): if the features within a dataset are highly structured, such that their values correlate strongly, the presence of additional correlated features will increase our ability to separate distinct samples. This implies that the CoD may be viewed as the presence of a large number of features whose values are random in relation to groups of similar samples. In scRNA-seq, such features correspond to genes that do not inform cell state, such as house-keeping genes. It has been estimated that of the tens of thousands of distinct transcripts captured in a typical scRNA-seq assay, only 3000-5000 of them relate to cell type specific expression patterns (Ramskö Ld et al. 2009).

To overcome the high dimensionality of scRNA-seq data, several methodologies have been developed (Kiselev, Andrews, and Hemberg 2019; Wu and Zhang 2020). The most commonly used are feature extraction and highly variable gene (HVG) selection. Feature extraction methods such as principal component analysis (PCA), and Uniform Manifold Approximation and Projection (UMAP) (McInnes et al. 2018) attempt to compress a high dimensional dataset into a smaller set of highly informative features. HVG selection seeks to identify a subset of genes more predictive of distinct cell types than randomly expressed genes. While it is a widely used pre-processing strategy, HVG selection can struggle to account for important but lowly expressed genes, or genes present in only a small fraction of cells (Källberg, Vidman, and Rydén 2021). Furthermore, evaluation of various HVG methods found that different techniques show poor overlap in HVG suggested from the same datasets, and that highly expressed genes were often incorrectly flagged as HVGs (Yip, Sham, and Wang 2018). This poor consistency may arise because gene selection is carried out in a univariate manner based on a weak mechanistic assumption that genes with high expression variance correspond to different cell types.

In this work, we introduce a mathematical framework termed Entropy Sorting (ES). ES allows us to simultaneously measure the correlations between features while quantifying the likelihood that these correlations have been weakened due to the introduction of technical error, such as drop outs. We encode ES in an algorithm called FFAVES: Functional Feature Amplification Via Entropy Sorting. We use FFAVES to amplify the signal of groups of co-regulating genes in an unsupervised, multivariate manner. By amplifying the signal of genes with correlated expression, while filtering out genes that are randomly expressed, we can identify a subset of genes more predictive of different cell types. The output of FFAVES can then be used in our second algorithm, Entropy Sort Feature Weighting (ESFW), to create a ranked list of genes that are most likely to pertain to distinct sub-populations of cells in a scRNA-seq dataset. Unlike HVG selection, ESFW performs gene selection in a multivariate manner that specifically seeks to identify genes with consistent expression patterns, indicative of a distinct cellular identity.

## 2 RESULTS

### Entropy Sorting

#### Pairwise Feature Correlations

The foundation of ES is to create a correlation metric analogous to metrics such as Pearson’s Correlation or Mutual Information to define a common structure between the discrete states of two features. To do so, we re-imagine conditional entropy (*CE*) as a sorting problem between features whose samples can display two states, e.g. a gene being functionally active or inactive. As such, discretisation of continuous data is a requirement for ES (SI 1). Given two features, *CE* ∈ [0, 1] quantifies the information needed to predict the state of one feature for a given sample when conditioned on the other feature. For example, *CE* = 0 indicates that the observed state of the conditioned feature is entirely determined upon observing the state of the second feature.

To develop *CE* into a sorting problem, we consider a toy example to guide the theoretical exposition (Fig. 1A) in which 30 samples (cells) display discrete states for two features (genes). We hypothesise that partitioning all samples by the observed states of one feature will perfectly sort the states of the other. We designate the partitioning feature the Reference Feature (RF). We then seek to quantify to what degree the RF sorts the states of the second feature, the Query Feature (QF). We calculate *CE* for the RF/QF pair via the Entropy Sort Equation (ESE, Fig. 1B). A detailed derivation of the ESE is provided in SI 2. Partitioning samples according to the RF forms two groups (*G*_*i*_, *i* = 1, 2, Fig. 1A). We can calculate the entropy of each group as a function of the number of QF minority states in *G*_1_: 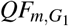 (hereafter denoted as *x*).

**Figure 1:**
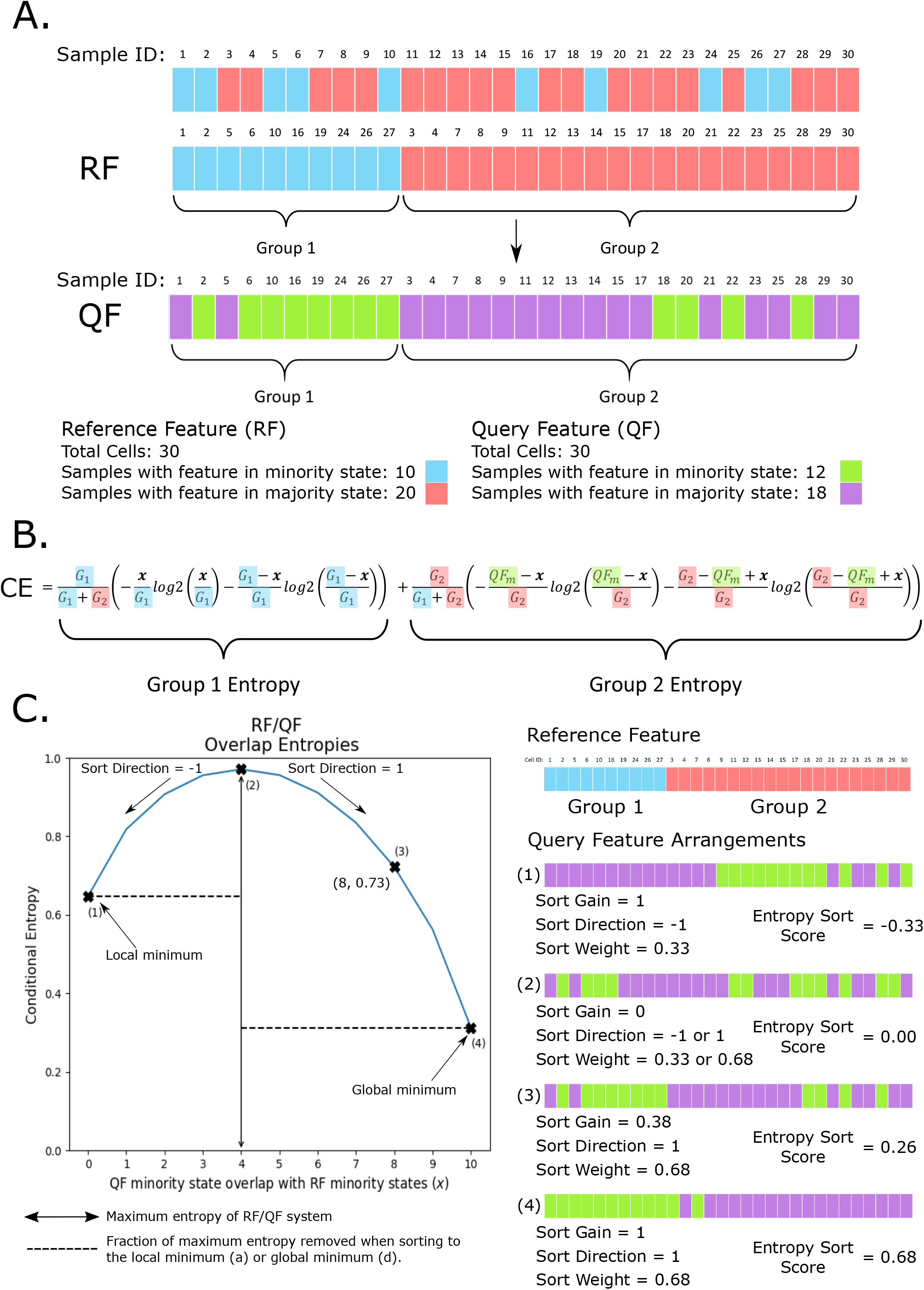
Quantifying the dependent relationship between two features. **A**. A toy example. The states of each RF sample are sorted into two groups. The QF is then inspected while maintaining the RF sample ordering. **B**. The ESE for calculating *CE. G*_1_ (group 1) and *G*_2_ (group 2) are the number of minority or majority states of the RF respectively. *QF*_*m*_ is the total number of QF minority states. For brevity, we use *x* to denote 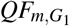, the number of QF minority states that overlap with the RF minority states, which is the only independent variable. Each constant is highlighted with their corresponding colours in panel A. **C**. Given any observed pair of features, we may form an ESE parabola by fixing the constants of the ESE and calculating the CE for different values of *x*. Points **(1)** and **(4)** correspond to the local and global minimum. **(2)** is the maximum *CE*, where the RF and QF are independent. **(3)** is the *CE* corresponding to the observed arrangement in A. Each of **(1-4)** is illustrated by an example arrangement.

The ESE defines a smooth parabolic function for calculating *CE*(*x*). We plot the ESE parabola for our toy example in Fig. 1C, and highlight four points of interest on the curve. Points (1) and (4) are the boundaries of the ESE, since *x* ≥ 0 (1) and *x* ≤ *G*_1_ (4). Point (2) corresponds to the maximum *CE*, the point at which the RF/QF pair are independent. Point (3) is *CE* of the observed samples in Fig. 1A. The ESE parabola represents a common structure for any RF/QF pair that we use to define our correlation metric, which is important for two reasons. It demonstrates that any RF/QF pair, regardless of sample number or minority/majority state cardinality, has a quantifiable common structure. Further, it defines a mathematical framework that can be used to calculate the relationship between two features under the assumption that the features are dependent upon one another. Later we consider the assumption of feature independence, to quantify which of the two hypotheses is more likely.

The ES correlation metric can be broken down into three parts: Sort Direction (SD), Sort Weight (SW), and Sort Gain (SG). SD describes whether the value of *x* corresponds to an enrichment of QF minority states in *G*_1_ or *G*_2_, such that

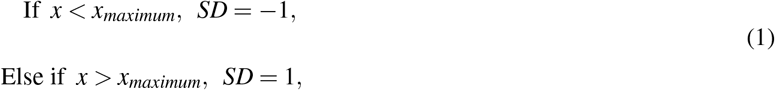

where *x*_*maximum*_ is the value of *x* at the maximum of the parabola. If SD = 1, the system is sorting towards the global minimum, since the minimum when *x > x*_*maximum*_ always has a lower *CE* than when SD = -1. SW is maximum amount of entropy that would be removed from the system if it existed at either minima. Hence it is dependent on SD:

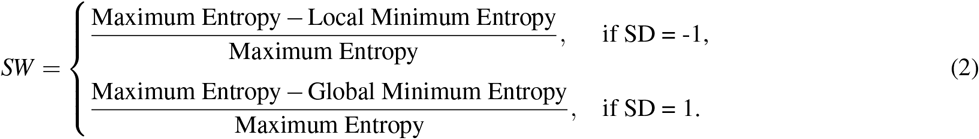

Lastly, SG is similar to a well established metric from Information Theory: Information Gain (IG) (Quinlan 1986). IG refers to the decrease in entropy from the scenario in which QF and RF are independent to when QF is dependent on RF. SG is the same decrease in entropy, but as a fraction of the total entropy that would be removed at the relevant minimum. Formally,

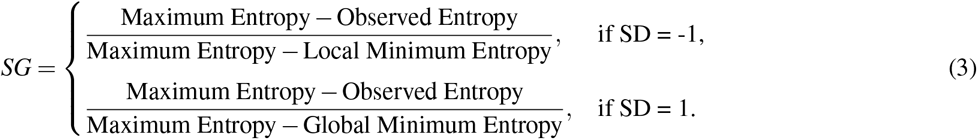

The Entropy Sort Score (ESS) is the product of these values for any RF/QF pair:

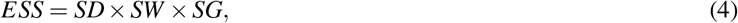

where *SD* = ±1, 0 ≤ *SG* ≤ 1 and 0 ≤ *SW* ≤ 1. Hence for any RF/QF pair, *ESS* ∈ [−1, 1], similar to other correlation metrics (Fig. S1).

### Divergence

The ESE parabola describes how *CE* changes for a RF/QF system with fixed minority/majority state proportions, but varying RF/QF dependencies. To quantify to what degree observed RF/QF samples appear to be in the wrong state, we introduce divergence: a measure of how far away the observed states of a RF/QF pair are from an optimally dependent system.

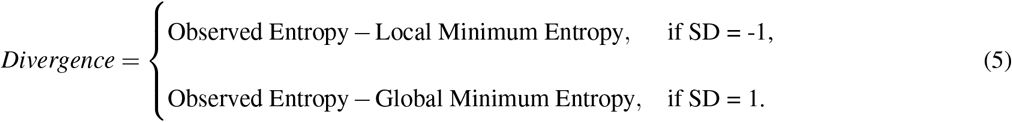

To demonstrate the concept of divergence, we consider three toy examples (Fig 2A). Example (i) represents the ground truth, where a strong but imperfect dependent relationship exists between the observed states of the RF/QF pair. Here we observe the maximum number of minority state RF/QF samples overlap, indicated by the samples that have blue and green expression states (*x* = 10). Accordingly, *CE* is equal to the global minimum of the ESE parabola. In (ii) we introduce an error such that one of the minority state QF samples is incorrectly observed as a majority state (marked “X”). This changes the parameters of the ESE, causing the parabola to shift (blue to orange). Since the error occurs in the non-overlapping region of the RF/QF minority states (the green minority state where the dropout occurred does not overlap with a blue minority state in the ground truth), the observed system still exists on the global minimum and no divergence has been observed. In example (iii) we introduce an erroneous data point in the overlapping minority state samples. This does not alter the parameters of the ESE, so the parabola is unchanged (orange), but the observed *CE* is greater than the global minimum. This movement away from the global/local minimum (*SD* = 1*/* − 1) is the divergence, calculated by Eqn. 5.

**Figure 2:**
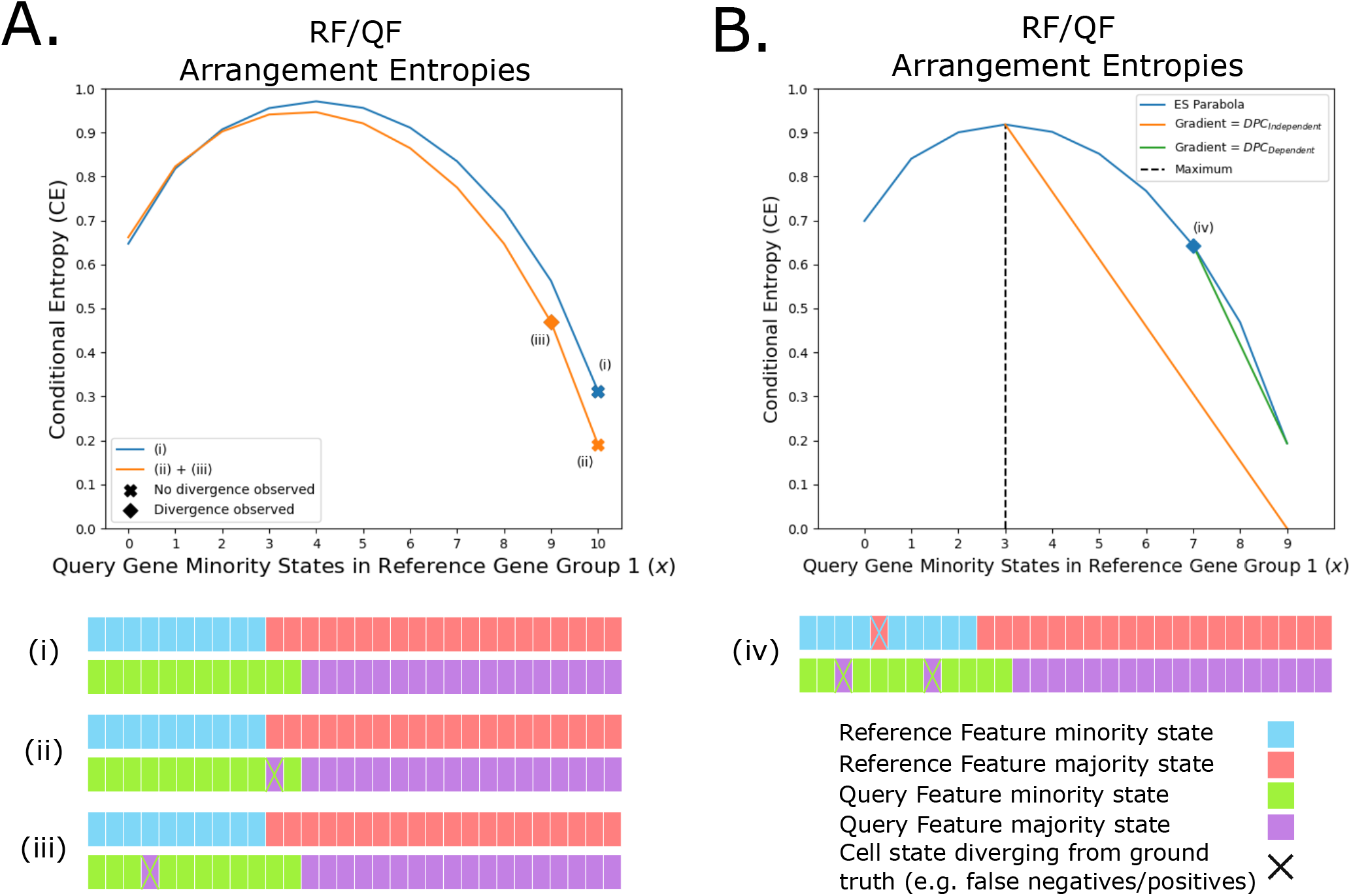
ES divergence and error potential. **A**. ESE parabolas highlighting three toy examples to demonstrate divergence: (i) ground truth for a partially dependent system; (ii) a FN drop out added that does not produce observable divergence on the ESE parabola; (iii) the addition of a FN drop out that generates observable divergence. **B**. DPC (Eqn 6) introduced to the RF/QF pair due to erroneous data points (example (iv)) under both the assumption of either RF/QF dependence (green line), or independence (orange line).

The simple examples in Fig 2A introduce scenarios where error can be observed through divergence. We have determined that 8 distinct categories cover all scenarios in which error in a RF/QF system may be observed through divergence (Fig S2). Distinguishing between these scenarios is important for the implementation of ES, as discussed later.

#### ES Hypothesis Testing

We use the concept of divergence to perform hypothesis testing. We test both the hypothesis that the features are dependent, such that observed divergence is due to technical noise/error, and the null hypothesis that the features are independent.

Under the assumption of dependence, the ESE quantifies the distance an observed RF/QF pair is from optimal dependence. We consider each divergent state – a state that moves the observed system away from the local/global minimum. We assign a proportion of the total divergence to each cell showing a divergent state, as the divergence per cell (DPC),

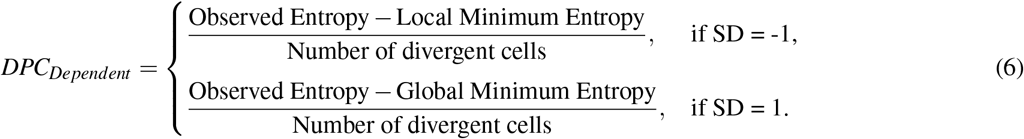

When *SD* = 1, divergent cells are those where a RF minority state overlaps with a QF majority state (Fig 2A (iii) and B (iv)). When *SD* = −1, divergent cells occur when a RF minority state overlaps with a QF minority state (Fig SI 3C, D). We visualise the DPC for example (iv) in Fig 2B as the gradient between the the global minimum and the observed entropy (green line).

Under the null hypothesis, we assume RF/QF are independent and the observed minority state overlap (*x*) has occurred by chance. Therefore, after a sufficient number of re-samplings of observed states from both RF and QF, we would expect on average that *x* and *CE* would equal their values at the maximum of the ESE parabola. Hence, under the null hypothesis,

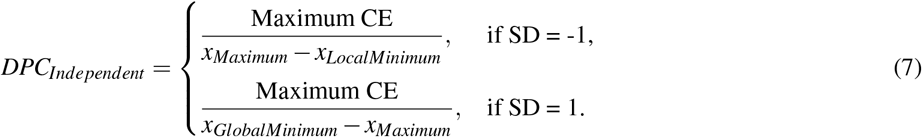

In Fig 2B, *DPC*_*Independent*_ is the gradient of the orange line.

We combine *DPC*_*Dependent*_ and *DPC*_*Independent*_ to define our final metric, the Error Potential (EP). EP allows us to compare whether observed divergence is more likely due to the hypothesis that RF and QF are dependent and error has been introduced to the system, or instead that the features are independent:

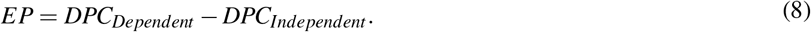

*EP >* 0 indicates that hypothesis of dependence holds, while *EP <* 0 indicates that the null hypothesis holds. In our software, we use EP to minimise the presence of erroneous data points, and in turn, amplify the signal of features that have dependent relationships. For further detail, please see SI 3. Further, in SI 4 we explain how to identify which feature is the RF and QF for any pair.

### FFAVES and ESFW

The metrics defined by ES are encoded in two algorithms. The first, FFAVES, uses ES to identify data points in a discrete matrix that are statistically likely to be displaying the wrong state. By correcting these data points, we aim to amplify the signal of feature correlations. The second algorithm, ES Feature Weighting (ESFW), assigns a importance weight to each feature in the data. Higher weights indicate that a feature is more likely to belong to a set of dependent features, while lower weights pertain to features that are randomly expressed throughout the data.

FFAVES and ESFW encode the mathematical framework of ES to perform multivariate expression state correction and feature importance weighting, respectively. EP is the cornerstone of both software, enabling unsupervised analysis while identifying combinatorial gene expression patterns. EP implicitly identifies whether the relationship between any two features is more likely to have occurred by chance or due to some functional relationship, without any need to define a threshold for this decision. This simultaneously allows the software to ignore uninformative pairs of genes in a manner that directly addresses the CoD. To the authors’ knowledge, there are no alternative unsupervised multivariate feature selection techniques available that are specialised for bioinformatics (Y, I, and P 2007).

Fig 3 provides a workflow for FFAVES and ESFW, while detailed descriptions are found in SI 5 and SI 6. We emphasise that the aim of FFAVES and ESFW is to identify genes that are consistently co-expressed within a scRNA-seq dataset. This does not necessarily constitute all genes that have unique expression patterns within distinct cell populations. Some genes may be missing due to technical limitations or poor discretisation. Rather, by filtering to ES selected genes we amplify the predominant expression structure in the data. Subsequently, users should consider how other genes may relate to the refined resolution of cell populations. This is illustrated below, in our analysis of human pre-implantation embryo data.

**Figure 3:**
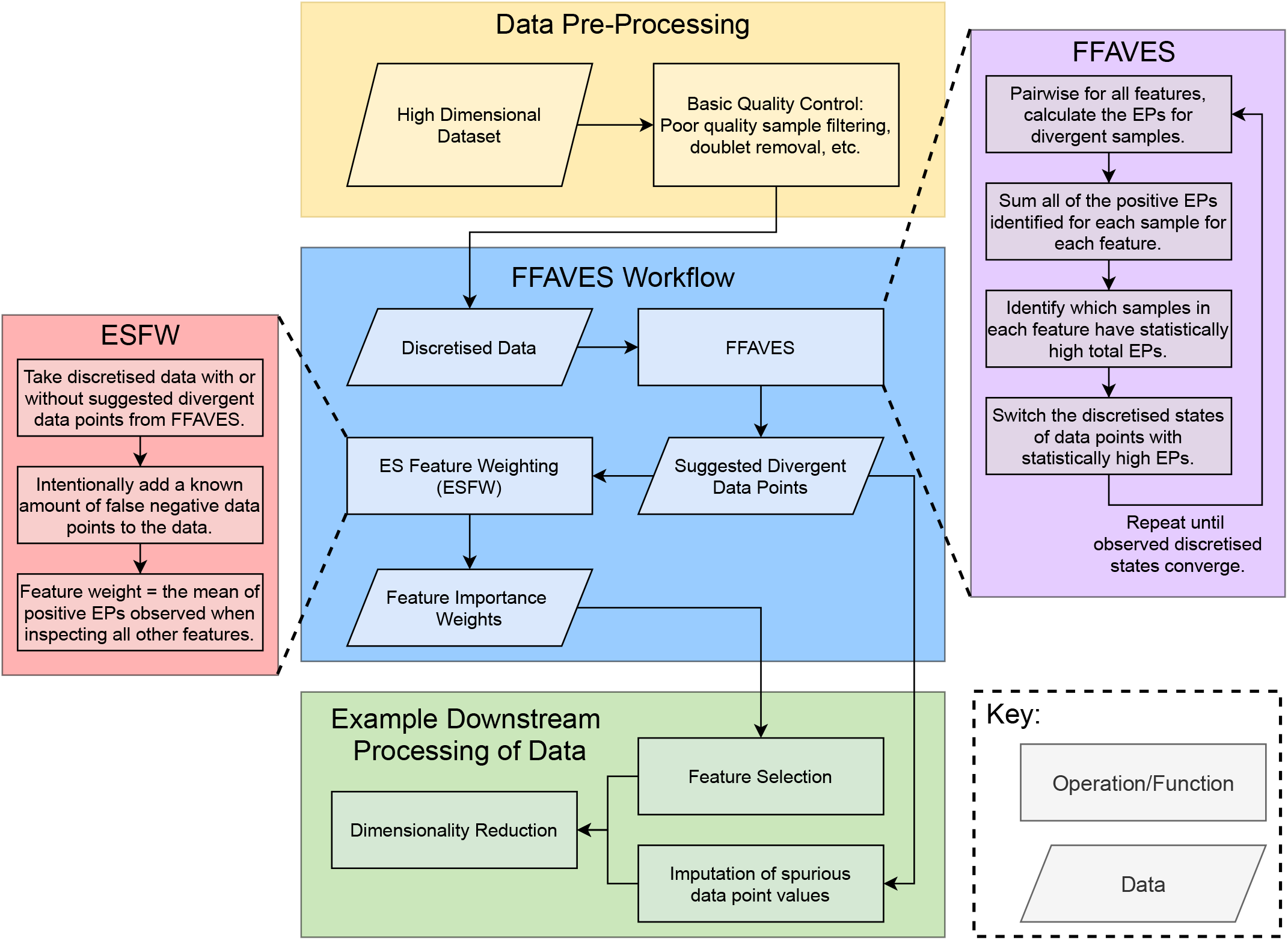
FFAVES and ESFW workflow. Yellow, blue and green boxes provide the proposed workflow to apply FFAVES and ESFW to high dimensional data for unsupervised feature selection. The purple and red boxes outline each algorithm.

### Synthetic Data

We curated a simple synthetic dataset with known ground truth to quantify the performance of FFAVES and ESFW (Fig. 4A, with further detail in SI 8). Briefly, we define 5 synthetic cell types according to sets of 200 “highly structured” genes that are tightly and uniquely co-expressed (Fig. 4E, bottom panel). Subsequently, we introduce random drop outs to these cells to mimic technical error (Fig. 4E, top panel). We include an additional 500 randomly expressed genes that are designated as “uninformative” genes, as well as “leaky gene expression”, which refers to false positive (FP) expression of highly structured genes outside of a gene’s specific cell type. We introduce 50 multi-modal genes that have a “medium” expression in one cell type and “high” in a second. Finally, we created a set of multiple cells (20% of all cells) by randomly combining expression profiles from the 5 synthetic cell types.

**Figure 4:**
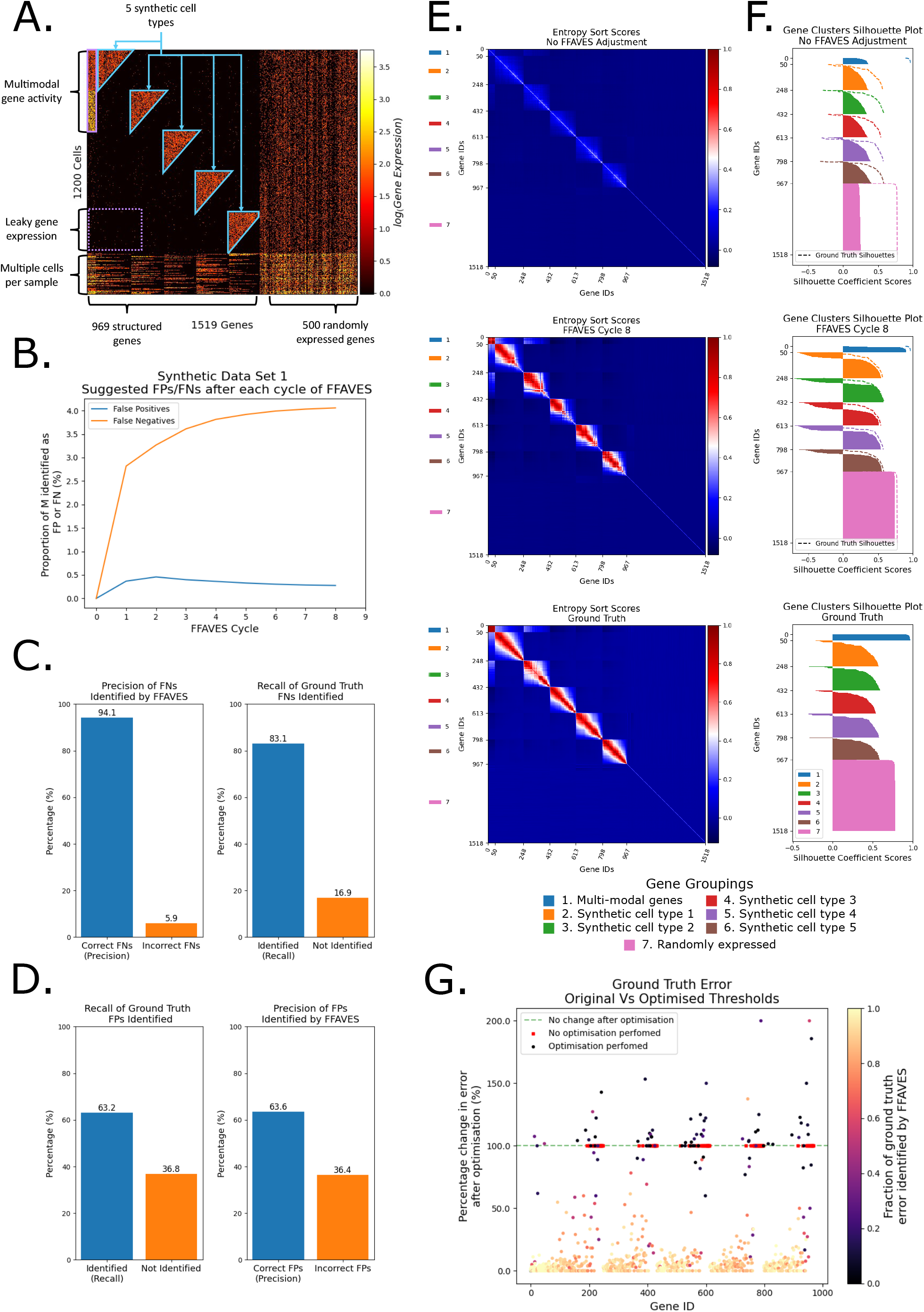
FFAVES accurately identifies false negatives and false positives. **A**. The synthetic scRNA-seq dataset. **B**. Convergence of FN/FP data points identified after each cycle of FFAVES. **C, D**. Precision and recall scores of FNs and FPs identified by FFAVES, respectively. See Fig. S6 and Fig. S7 for visualisation of where FNs/FPs occur in the data. **E**. Heatmaps of pairwise feature ESSs. Top - before identification of FNs and FPs by FFAVES. Middle - after application of FFAVES. Bottom - Ground truth, i.e. synthetic data prior to introduction of FN dropouts. **F**. Silhouette scores of the 7 main gene groups calculated from the respective ESSs in E. Dashed lines outline the ground truth silhouette scores. **G**. Reduction in FN errors that were intentionally introduced by sub-optimal feature discretisation.

We use this relatively simple representation of scRNA-seq data to dissect the performance and limitations of FFAVES and ESFW, as it facilitates a level of detailed analysis that could not be obtained from a dataset generated by stochastic simulation. We further apply ES to synthetic data generated by the Dyngen scRNA-seq simulation software (Cannoodt et al. 2021), as shown in Figs. S3, S4 and S5.

#### FFAVES accurately identifies FN and FP data points

We apply FFAVES to our first synthetic dataset to illustrate how the algorithm amplifies co-regulatory patterns of the 969 structured genes in an unsupervised, multivariate manner. First, to discretise the data, we sampled a discretisation threshold from a 𝒩 (4, 0.2) distribution for each gene (Fig. S9B). Since the mean expression for each gene is 5, these thresholds increase the probability that a cell in which the gene has non zero expression (Fig. 4A) will display that gene as inactive in the discrete matrix. This introduces FNs caused by sub-optimal discretisation strategies, which we do intentionally to demonstrate that FFAVES can automatically account for such problems.

After 8 cycles of FFAVES, the system converges on a set of suggested FN/FP data points (Fig. 4B). We can then compare the discrete expression matrix output by FFAVES with the ground truth matrix (Fig. SI 13A). FFAVES performs well, with precision and recall scores of 94.1% and 83.1% respectively for the identification of FNs, and 63.2% and 63.6% for FPs (Fig. 4C and D, Fig. S6). We verified that these precision and recall scores are robust across repeated stochastically generated synthetic datasets (Fig. S8).

The majority of FNs that FFAVES fails to identify occur in genes active in very few cells, even in the ground truth data with no intentional FNs (Fig. S6D). This represents a limit of sensitivity where genes expressed in roughly 20 or fewer samples are difficult for FFAVES to repair (SI 7). Although there is a drop in performance for FP identification, FFAVES is designed to be intentionally conservative when suggesting FP data points to maximise the accuracy of FN identification (SI 5). Furthermore, the authors are unaware of an existing methodology that can discriminate between FN and FP data points. We also find that FFAVES does not suggest that any of the 500 randomly expressed genes contained FNs or FPs (Fig S6, Fig 4). This is because ES Hypothesis testing can discriminate between overlapping minority states between independent and dependent features.

In Fig. 4E, we show the ESS (Eqn. 4) for all gene pairs to illustrate the recovery of gene co-expression relationships. We compare the synthetic data with drop outs (top) with after the application of FFAVES (middle) and the ground truth (bottom). Fig. 4F corroborates Fig. 4E, showing that the silhouette scores for the 7 main gene expression groups are corrected to values close to those of the ground truth. The small groups of genes with high negative scores (Fig. 4F, middle) are the same genes as previously highlighted, which are expressed in a small number of cells such that FFAVES struggles to recover ground truth co-expression patterns (Fig. S6D). These genes have expression profiles more similar to randomly expressed genes than the repaired highly correlated genes. Hence the negative scores of genes in clusters 2-6 indicate that they should be part of randomly expressed genes cluster (pink), rather than their ground truth cluster.

Finally, Fig. 4G demonstrates the ability of FFAVES to account automatically for sub-optimal discretisation thresholds. Controlled identification of such FNs is equivalent to correcting the discretisation threshold (Fig. S9). Since we know which FNs resulted from the sub-optimal discretisation process, we can quantify the proportion that were corrected by FFAVES. For our synthetic dataset, FFAVES correctly identified 83.1% of these FNs. Once again, a large proportion of the 16.9% FNs not identified correspond to genes for which the number of cells with active expression was below the sensitivity of FFAVES (Fig. 4G dark or red markers, Fig. S9).

#### FFAVES facilitates accurate false negative imputation

To compare against other tools designed to repair FN data points, we used the FNs suggested by FFAVES to perform imputation. To focus on quantifying correctly identified FNs, we chose a simple method for estimating the values of suggested FNs, the *impute*.*IterativeImputer* function from the *sklearn* Python package. This yields an imputed gene expression matrix for comparison with other scRNA-seq imputation software: MAGIC (Dijk et al. 2018), ALRA (Linderman et al. 2022) and SAVER (Huang et al. 2018). These tools were chosen as top-ranking examples of software that cover the three main classes of imputation methods (Hou et al. 2020) – smoothing (MAGIC), low rank matrix-based approximation (ALRA), and probabilistic modelling (SAVER).

Fig. 5A uses UMAPs to visualise how well each imputation method repairs the synthetic data with drop outs and batch effects. Qualitatively, FFAVES performs best, yielding an embedding most similar to the ground truth. The most notable difference in the MAGIC and ALRA embeddings is the amplification of batch effects, observed as tight groups of the same cell types separated by batch (yellow and black clusters). When the ground truth is known, identification of spurious imputations such as batch effect amplification are easily identified. However, without a ground truth, identifying spurious imputation becomes more difficult. This can lead to clusters of cells that are computational artefacts, rather than true biological signals (Hou et al. 2020; Andrews and Hemberg 2019). SAVER performs better than MAGIC and ALRA in mitigating noise due to batch effects. However, SAVER recovers a lower resolution of cell type heterogeneity compared to FFAVES, MAGIC and ALRA, indicated by looser connectivity between local cell populations.

**Figure 5:**
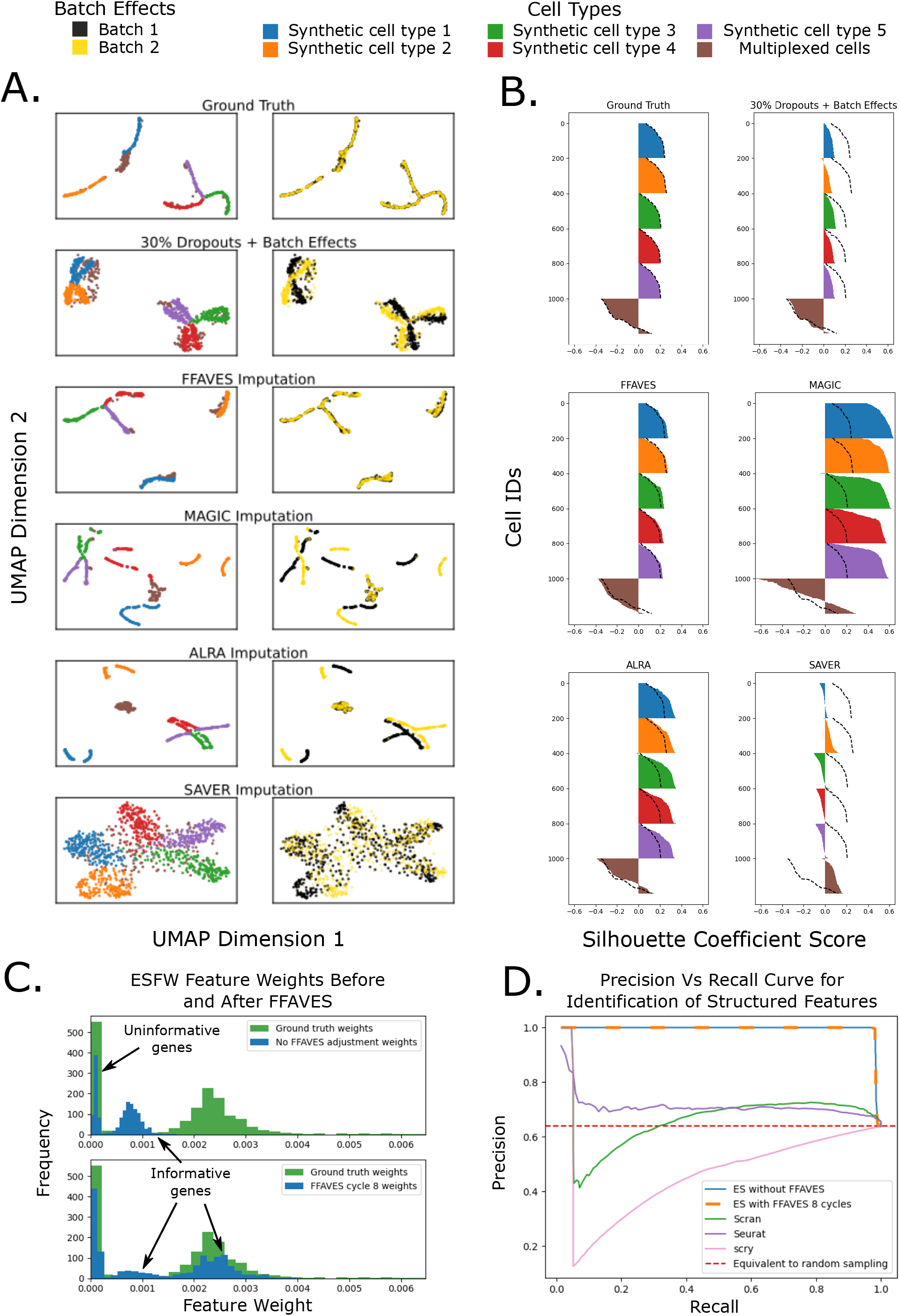
Performance of FFAVES and ESFW against comparable software. **A**. UMAPs of the synthetic dataset before and after imputation. The top two plots show the synthetic data before and after FN drop outs were introduced, with no imputation. **B**. Silhouette scores for each of the 6 main clusters of cells in the synthetic dataset. Black dashed lines mark the silhouette scores of the ground truth data prior to the introduction of FNs. **C**. Feature importance weights for all genes in the synthetic data according to ESFW. Top: Feature weights estimated from the synthetic data with FNs introduced to the ground truth. Bottom: Feature weights estimates after FFAVES has identified statistically significant divergent data points. **D**. Precision/Recall curves for distinguishing structured and randomly expressed genes. Each line is generated from the ranked gene lists of the respective feature selection software.

In Fig. 5B we use silhouette scores for 6 cell type clusters to quantitatively assess imputation performance. Imputation via FFAVES recovers cell clusters that closely resemble the ground truth. The silhouette scores after MAGIC imputation suggest data overfitting – samples appear considerably more similar to each other than in the ground truth. Such overfitting is an example of how metrics such as the silhouette score can be misleading without a ground truth for context. Higher silhouette scores can be incorrectly assumed to be synonymous with better imputation. The ALRA silhouette scores are similar to FFAVES and the ground truth, albeit with some slight overfitting. Finally, the SAVER silhouette scores are worse than the dropout + batch effect synthetic data. In particular, some cells have negative silhouette scores, indicating that they no longer cluster with their original cell type labels.

#### ESFW accurately discriminates informative features

The ESFW algorithm is designed to weight features using an unsupervised process that identifies combinatorial gene expression patterns. Higher weights indicate those that are more likely to be informative for sample identities/clusters. For a detailed description of ESFW see SI 6. We used ESFW to calculate the feature weights for our drop out + batch effect synthetic data before/after FFAVES, as well as for the ground truth data. Even without applying FFAVES, a bimodal distribution of weights emerges (Fig. 5C, top). Those genes with feature weights close to or equal to 0 are the uninformative genes. The second group with feature weights around 0.001 comprise over 97% of the 969 highly structured informative genes. However, as expected, these highly structured genes in the drop out + batch effect synthetic data have feature weights lower than that of the ground truth data. After applying FFAVES, we find that a large proportion of the informative genes have feature weights that closely resemble those of the ground truth (Fig. 5C, bottom), further demonstrating that FFAVES accurately re-captures feature dependencies. The small fraction of informative genes that retain feature weights of around 0.001 comprise the same genes previously suggested to have ground truth minority state cardinalities too low for FFAVES to identify and repair.

Since ESFW provides a score for each feature with regards to sample structure, we can form a ranked list of gene importance. We can use ranked gene lists from ESFW and other feature selection software to compare their ability to distinguish between the highly structured cell type specific genes and the uninformative genes in our synthetic data. Because feature selection is a common and important step in workflows for analysing scRNA-seq data, several implementations have been developed. We selected three unsupervised and popular tools (Yip, Sham, and Wang 2018) for comparison: Scran (Lun, McCarthy, and Marioni 2016), Seurat (Hao et al. 2021), and scry (Townes et al. 2019).

Fig. 5D presents the precision/recall curves from applying each tool to our synthetic data. For all plots except the results obtained after applying FFAVES to the dropout synthetic data (orange dashed line), feature selection was performed on the synthetic data with drop outs + batch effects. The ESFW precision/recall curves show high discrimination between structured and randomly expressed features, up to a recall of 0.97. Conversely, Seurat, Scran and scry show a considerable drop in precision at recall values less than 0.1, indicating that these methods struggle to differentiate between the informative and uninformative genes. Precision scores below the red dashed line are those where a higher fraction of uninformative genes are inspected than would on average be observed under random sampling of genes.

### FFAVES and ESFW reveal a high resolution scRNA-seq embedding of the human pre-implantation embryo

To test the utility of ES for examination of real biological systems, we considered human pre-implantation embryo scRNA-seq data containing 1751 cells and 34054 genes, compiled by Meistermann et al. 2021. Using FFAVES and ESFW, we identified a set of 3700 highly informative genes. Restricting the expression matrix to these genes we generated UMAPs that identify distinct cell type populations along the developmental time course (Fig. 6A, Fig. S10). Thus, the embryonic day (E) labels progress chronologically from the bottom right to the top left of the UMAP. Importantly, we generated this high resolution embedding without any augmentation of the original expression matrix – no data transformations, batch correction, smoothing or imputation. This gives confidence that any cell similarities or gene expression signatures identified are likely to be biologically significant, rather than introduced during computational pre-processing.

**Figure 6:**
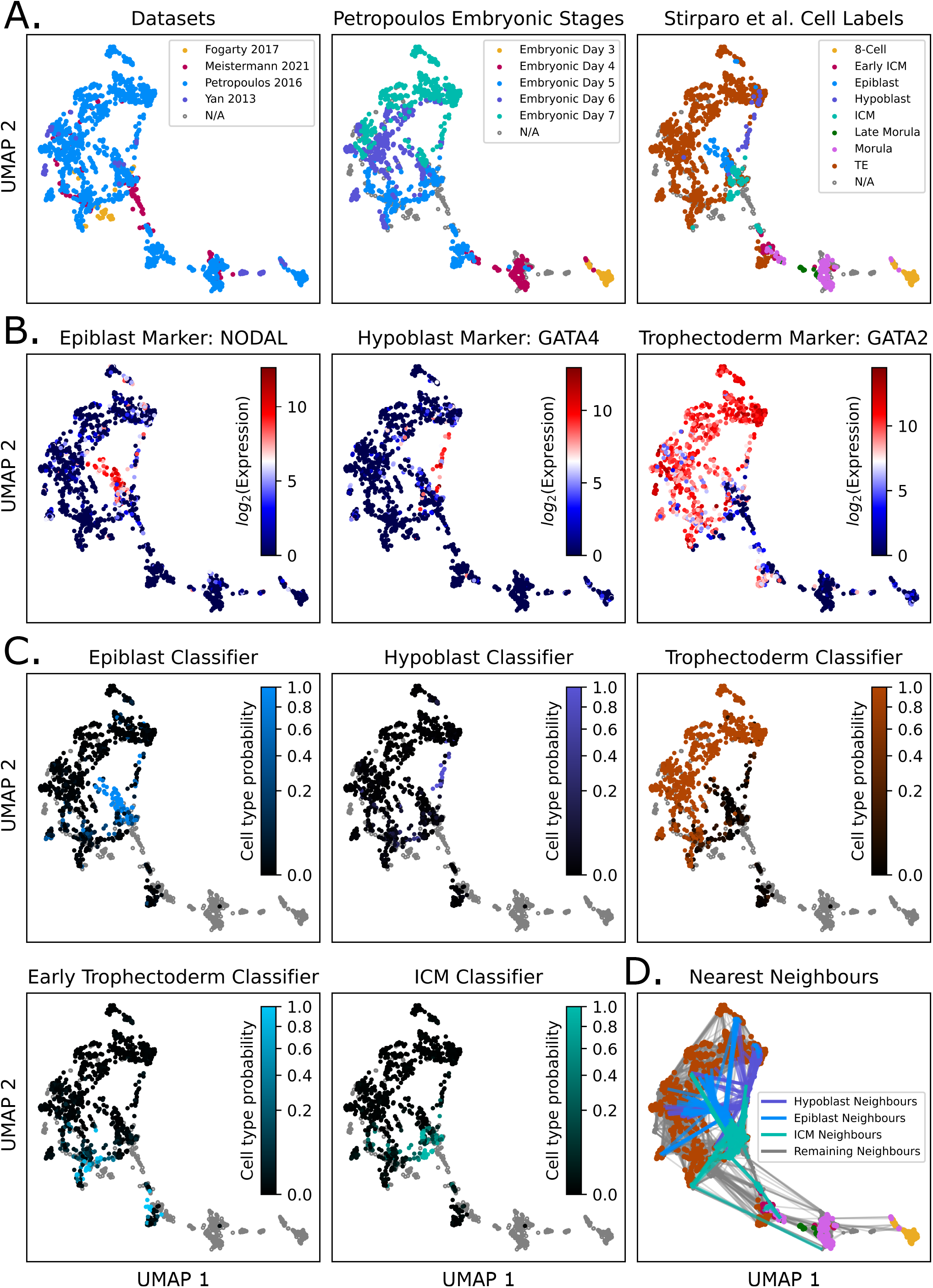
Independent validation of the FFAVES + ESFW human pre-implantation embryo embedding. **A**. The FFAVES + ESFW UMAP embedding overlaid with different label information: left to right, the datasets that samples originate from, the time point labels from the Petropoulos dataset, and the cell type labels for the Petropoulos dataset that were independently assigned by Stirparo et al. 2018. **B**. Example epiblast, hypoblast and trophectoderm marker expression. See Fig. S13 for more examples. **C**. Predicted cell type probabilities of individual cells from a classifier trained on human pre-implantation embryo scRNA-seq data from the independent Yanagida et al. 2021 dataset. Grey samples are those that were not processed by the classifier to avoid confounding variables such as batch effects. See Fig. S14 for the same analysis with Macaca classifiers. **D**. Nearest neighbour embedding where each cell is connected by lines to their 10 most similar samples according to gene expression. See Fig. S15 for individual cell type nearest neighbour embeddings.

An initial comparison of our UMAP embedding with the labels identified by the Meistermann et al. 2021 analysis indicates generally good agreement with their supervised analysis (Fig. S11). However, their analysis contains discrepancies among proposed Epiblast (Epi) and Hypoblast (Hyp) populations, and in contrast to Meistermann our study identifies a distinct early ICM population. Identification of the ICM in human embryos has been a subject of debate that has provoked alternative models of early lineage segregation (Weltner and Lanner 2021).

#### Independent validation of our UMAP embedding

To examine and validate the UMAP embedding, we compare it against previous analyses of human and primate pre-implantation embryos. Stirparo et al. 2018 analysed the Petropoulos et al. 2016 dataset (which accounts for 85% of the data in Meistermann, 2021), and proposed cell type annotations based on known gene expression signatures and unsupervised clustering. Overlaying Stirparo’s annotations onto our embedding, we find distinct groups of cells, indicating that the UMAP identifies cell clusters with specific gene expression signatures (Fig. 6A). The increased clustering performance when using the 3700 ES selected genes compared to the 4484 HVG genes selected by Meistermann et al. 2021 can at least partially be explained by the ES genes further distinguishing cell type clusters (quantified by Silhouette scores) for the ground truth cell types proposed by Stirparo et al. 2018 (Fig. S12). Thus our unsupervised feature selection approach is in better agreement with the supervised analysis of Stirparo et al. 2018 than HVG selection. Overlaying the UMAP with gene expression profiles of canonical Epi, Hyp and Trophectoderm (TE) markers (Stirparo et al. 2018; Amrani et al. 2019), shows consistency with the proposed cell labels (Fig. 6B, Fig. S13).

To demonstrate that the structure in our UMAP is conserved in independent datasets, we created two sets of cell type classifiers from: (i) human pre-implantation embryo cells Yanagida et al. 2021; (ii) cynomolgus monkeys (Macaca fascicularis) pre-implantation embryo cells Nakamura et al. 2016. We used the scANVI machine learning package (Gayoso et al. 2022) to train cell type classifiers on the cell labels allocated to the reference cells from Yanagida et al. or Nakamura et al. We then applied the classifiers to our UMAP embedding to predict cell types based on gene expression signatures. These analyses showed good agreement with the clusters identified in our embedding, with each cell type classifier scoring cells most highly in the expected regions of the UMAP (Fig. 6C, Fig. S14).

Together, the cell type labels from Stirparo et al. 2018, the cell type specific gene expression profiles, and the cell type classifiers created from independent human or Macaca pre-implantation embryo datasets provide four independent validations of our UMAP embedding. We conclude that FFAVES and ESFW have provided a higher resolution of cell type/gene expression dynamics than achieved by previous analyses of these data.

#### Defining the human pre-implantation embryo inner cell mass

There are two prevailing hypotheses regarding the establishment of the Epi, Hyp and TE lineages during human pre-implantation development. Petropoulos et al. 2016 concluded from scRNA-seq analysis that the three lineages may emerge simultaneously. However, mouse experimental embryology studies have established a two step model, with the first cell fate decision segregating TE from ICM at the late morula stage (Chazaud and Yamanaka 2016) after which the ICM differentiates into Epi and Hyp in the blastocyst. More recently, Meistermann et al. 2021 analysed human and mouse pre-implantation embryo scRNA-seq data with an aim to resolve which model is operative. Although Meistermann et al. 2021 found supporting evidence for the two step model in human development, they were unable to confidently identify an ICM population. In the absence of a clear ICM population, Meistermann et al. 2021 surmised that Hyp cells may emerge from the Epi.

In the FFAVES/ESFW UMAP embedding divergence into either TE or ICM populations is apparent at E5 (Fig. 6A, C). Proceeding from E5 to E6/7, the ICM cells differentiate into Epi and Hyp. As mentioned before, the proposed E5 ICM population has been previously suggested by Stirparo et al. 2018, but could not be resolved through dimensionality reduction techniques. The existence of the ICM is further supported by our classifier analysis utilising independently generated ICM gene expression signatures (Fig. 6C, Fig. S14). Furthermore, by nearest neighbour analysis (inspecting the 10 most similar cells based on gene expression for the suggested ICM, Epi and Hyp cells), we find that the Epi and Hyp cells are each connected to the ICM population, but have very little connectivity to each other (Fig. 6D, Fig. S15). The lack of connectivity between the Epi and Hyp cells supports the hypothesis that both differentiate from the ICM, rather than Hyp emerging from Epi.

Identification of a distinct ICM population enables us to suggest gene markers for future studies. We sought to identify genes whose expression was localised to the human ICM cells in our UMAP embedding. In our GitHub repository (see experimental procedures) we describe how these markers were identified and list more potential ICM markers. For validation we examined their expression in an independent tSNE embedding from Yanagida et al. 2021 (Fig. 7A, Fig. S16). We present two broad types of ICM markers. Those such as *FGF1* and *PRSS3* display expression specifically in the ICM labelled cells in both embeddings. The second set exhibit upregulated expression at E4 in addition to E5 ICM, and are markedly downregulated in the Epi, Hyp and TE populations. Examples of this group are *BHMT* and *SPIC*. We note that previous studies have proposed *SPIC* and *PRSS3* as human ICM markers (Singh et al. 2019).

**Figure 7:**
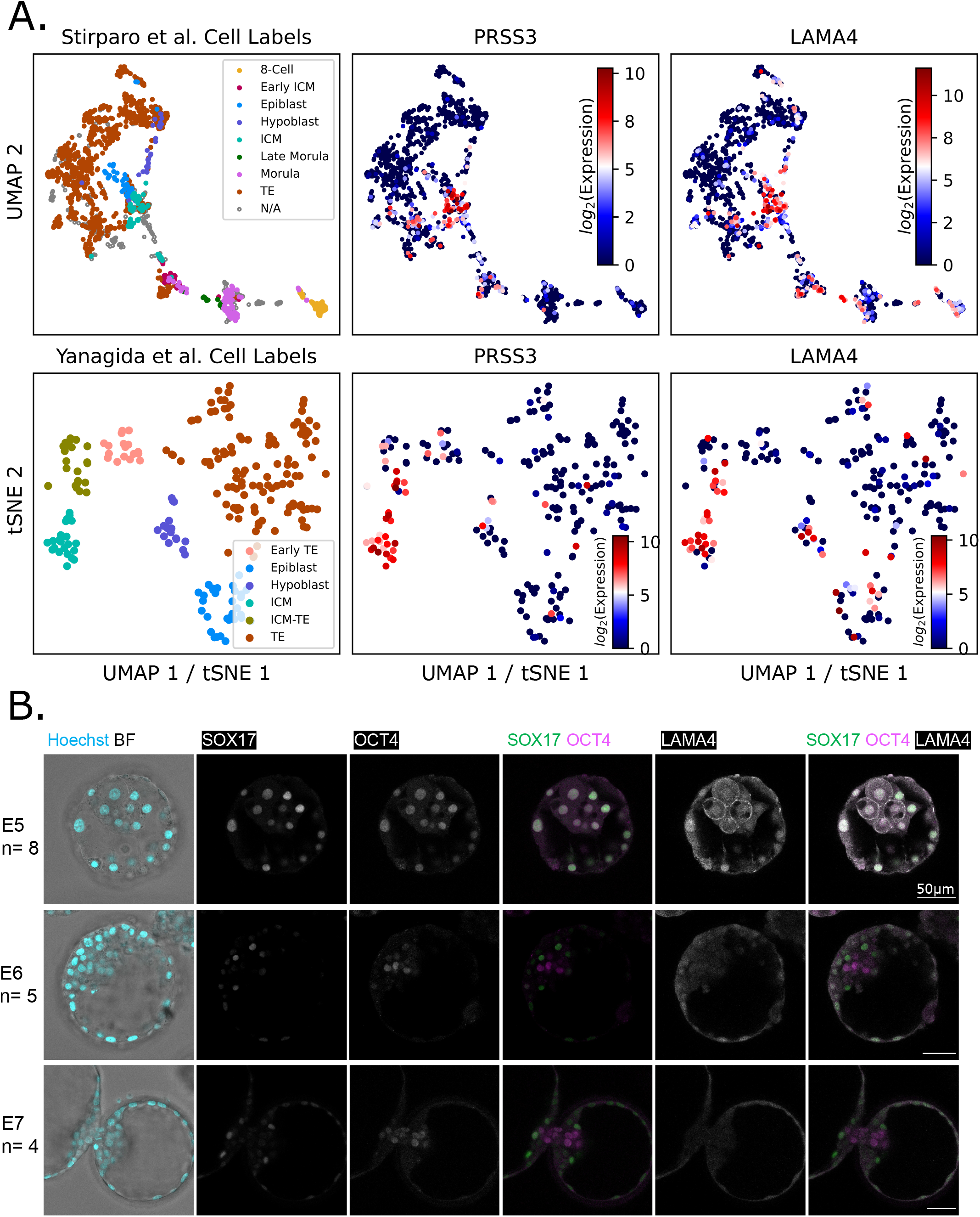
Identification of potential ICM markers. **A**. ICM markers were selected based on localised expression in the ICM population of the FFAVES + ESFW UMAP embedding (top row) corroborated in the tSNE embedding generated by Yanagida et al. (bottom row). See Fig. S16 and online methods for additional proposed ICM markers. **B**. Confocal images of human embryos immunostained for LAMA4 cell surface protein together with SOX17 and OCT4 nuclear transcription factors. Nuclei are visualized with Hoechst staining. The zona pellucida has been removed at E5 and E6 but not at E7 due to the embryo beginning to hatch. Staining patterns were consistent for all embryos examined: E5, n=8; E6, n=5; E7, n=4. Scale bar equals 50 µm.

#### Immunostaining validates LAMA4 as an ICM marker that is extinguished in Epiblast

To test whether markers identified from our UMAP embedding localise to the ICM, we performed immunostaining of LAMA4 on embryonic days 5 to 7 (E5-E7) human embryos. We selected LAMA4 as a cell surface protein with relatively high expression levels (>*log*_2_10) within the ICM cells (Fig. 7A) and with an available antibody reagent. At E5 (mid-blastocyst), LAMA4 is clearly localised to the cell surface of the ICM cells (Fig. 7B). Co-expression of OCT4 and SOX17 confirms that hypoblast and epiblast lineages are not yet specified (Niakan and Eggan 2013). By E6/7 (late blastocyst), LAMA4 expression is downregulated while OCT4+ and SOX17+ cells are segregated to epiblast and hypoblast fates respectively. These results provide validation that our embedding can identify the ICM population and specific markers prior to epiblast and hypoblast specification.

## 3 DISCUSSION

Over the last decade, the advancement of NGS techniques has markedly increased the types and quantity of data that can be obtained on genome control of cell behaviour (Anaparthy et al. 2019). While this increase in molecular information is exciting, it also presents new challenges around how best to analyse large, high dimensional datasets to generate biological insight (Angerer et al. 2017). In this work we present Entropy Sorting, a mathematical framework that quantifies the correlations between features (genes) in a high dimensional dataset as a sorting problem. The theory of ES is encoded in two algorithms: FFAVES and ESFW. Together, these provide unsupervised pre-processing to increase the resolution of information extracted from scRNA-seq data, and high dimensional data in general.

To demonstrate the effectiveness of Entropy Sorting, we applied our software to both synthetic and experimental scRNA-seq datasets. On synthetic data with known ground truth, we demonstrate that FFAVES and ESFW perform markedly better than popular HVG identification software at discriminating highly-correlated and randomly expressed genes. When compared to other popular imputation software (Dijk et al. 2018; Huang et al. 2018; Linderman et al. 2022), we show that FFAVES can identify FNs and FPs with high accuracy, and facilitates imputation such that ground truth cell-cell similarities are recovered. Furthermore, ESFW was shown to outperform current popular methods in performing feature selection to distinguish cell type specific genes from randomly expressed genes.

Applied to scRNA-seq data from human pre-implantation embryos (Meistermann et al. 2021), FFAVES and ESFW identified a subset of 3700 genes that were highly predictive of cell state. Filtering to these highly structured genes yielded UMAP embeddings with a higher resolution of gene expression dynamics during pre-implantation development than previously observed. Crucially, this was achieved by unsupervised filtering, without changing any values in the original gene expression matrix. Notably, FFAVES/ESFW revealed a distinct ICM population that precedes both the Epiblast and Hypoblast lineages. These analyses provide evidence for the two-step model of pre-implantation lineage segregation, which is well established in mouse development but has been disputed in human embryos due to failure of previous analyses to discriminate a distinct ICM population (Petropoulos et al. 2016; Meistermann et al. 2021; Stirparo et al. 2018). Immunostaining for LAMA4 shows ICM-specific expression at E5 with down-regulation in Epiblast and Hypoblast at E6/E7. This result substantiates the reliability of our embedding and demonstrates the potential to identify new lineage-specific markers for analysis of early human development.

FFAVES and ESFW should be viewed as pre-processing steps that help to maximise the signal of highly structured features. We chose not to apply any batch correction, imputation or feature extraction methods (other than UMAPs for visualisation) to the human embryo data. In doing so we show that ES is able to address the CoD in a manner that elucidates hidden structure in scRNA-seq data simply by removing spurious/uninformative features, rather than needing to augment or smooth the data. However, it may be possible to gain an even higher resolution of gene expression dynamics by applying other scRNA-seq analysis tools (Wu and Zhang 2020), post application of FFAVES/ESFW.

ES has the potential to be useful in other domains. Within the scope of NGS techniques, ES should be straightforward to apply to methods such as single cell ATAC-seq and BS-seq. Furthermore, the requirement to provide ES with discrete data may be advantageous for multi-omics single cell analyses (Anaparthy et al. 2019), which provide simultaneous read outs for multiple NGS techniques. Each technique can produce very different types of numerical outputs, such that combining them into a single dataset/readout is non-trivial. ES has the potential to overcome this challenge by discretising the data and framing the problem as the identification of functional relationships between the presence/absence of mRNA, chromatin accessibility/inaccessibility or sequence methylation/non-methylation. Therefore, ES offers the possibility of combining different types of data for coherent analysis. Finally, beyond NGS techniques, ES should be a powerful tool for reducing the complexity of a wide variety of high dimensional datasets, such as medical diagnostic or marketing data, in an efficient and unsupervised manner.

## 4 EXPERIMENTAL PROCEDURES

Computational workflows and data used for the generation of results in this article are available through the following sources.

### Data and code availability

The human pre-implantation embryo data may be found at https://data.mendeley.com/datasets/689pm8s7jc/draft?a=cc12423c-c19c-49b9-8cd9-d883064c048f. This repository also contains detailed workflows to reproduce our results.

The data used to create our UMAP embedding is a combination of raw counts scRNA-seq data from Yan et al. 2013, Petropoulos et al. 2016, Fogarty et al. 2017, and Meistermann et al. 2021, which was complied into a single gene expression matrix kindly provided by Meistermann et al. 2021. For information regarding data processing please refer to their manuscript.

The Yanagida et al. 2021 human embryo data analysed in this paper is available via GEO accession number GSE171820. The tSNE used in this paper for visualising the data was kindly provided by the authors.

The Nakamura et al. 2016 Macaca embryo data analysed in this paper is available via GEO accession number GSE74767.

Instructions to install FFAVES and ESFW can be found at, https://github.com/aradley/FFAVES. This repository also contains the synthetic data and workflows to reproduce the synthetic data results in this article.

### Human embryos

Supernumerary frozen human embryos were donated with informed consent by couples undergoing in vitro fertility treatment. Use of human embryos in this research is approved by the Multi-Centre Research Ethics Committee, approval O4/MRE03/44, and licensed by the Human Embryology Fertilization Authority of the United Kingdom, research license R0178.

Detailed descriptions of embryo preparation and embryo staining experimental procedures can be found in SI 9.

## Acknowledgements

This research was supported by the Biotechnology and Biological Sciences Research Council (BBSRC, grant number BB/P021573/1). A.R. was funded by a BBSRC PhD studentship (1943266) with co-funding from the Microsoft Research PhD scholarship program. A.S. is a Medical Research Council Professor (G1100526/1).

## Author contributions statement

Conceptualisation: A.R., S-J.D., A.S.; Data curation: A.R., E.C-S., J.N.; Formal Analysis: A.R.; Funding acquisition: S-J.D., A.S.; Investigation: A.R., E.C-S., J.N.; Methodology: A.R.; Project administration: S-J.D., A.S.; Resources: S-J.D., A.S., J.N.; Software: A.R.; Supervision: S-J.D., A.S.; Validation: A.R., E.C-S., J.N.; Visualisation: A.R., E.C-S.; Writing – original draft: A.R.; Writing – review & editing: S-J.D., A.S.

## Conflicts of interests

We declare no competing interests. Sara-Jane Dunn was an employee at Microsoft Research during this study and is currently employed at DeepMind. Microsoft Research provided co-funding for Arthur Radley’s research council studentship and access to computational resources. Neither Microsoft Research nor DeepMind have directed any aspect of the study nor exerted any commercial rights over the results.

## Notes

### Competing Interest Statement

The authors have declared no competing interest.

### Summary of Updates

We have updated the results in two major ways. 1) We include additional synthetic data sets that were generated using an independent method. These results demonstrate the generalisability of entropy sorting. 2) We include human embryo stainings to validate one of the markers we propose as a human pre-implantation ICM marker.

