## Supplemental figures 1-9 for "Entropy sorting of single cell RNA sequencing data reveals the inner cell mass in the human pre-implantation embryo"

### SUPPLEMENTARY MATERIAL

#### Supplemental Figures

| Antibody | Supplier | Catalogue number | Dilution |
|---|---|---|---|
| OCT4 (C-10) | Santa Cruz | #sc-5279 RRID:AB_628051 | 1:200 |
| SOX17 | R&D | AF1924 RRID:AB_355060 | 1:200 |
| LAMA4 | Invitrogen | # PA5-38938 RRID: AB_2555530 | 1:100 |
| Donkey anti-goat Alexa Flour 488 | Invitrogen | # A32814, RRID: AB_2762838 | 1:500 |
| Donkey anti-rabbit Alexa Flour 555 | Invitrogen | # A-31572, RRID: AB_162543 | 1:500 |
| Donkey anti-mouse Alexa Fluor 647 | Invitrogen | # A-31571, RRID: AB_162542 | 1:500 |
| Hoechst 33342 | Invitrogen | H3570 | 1:1000 |

**Table S1.** **Human embryo immunostaining antibody information.**

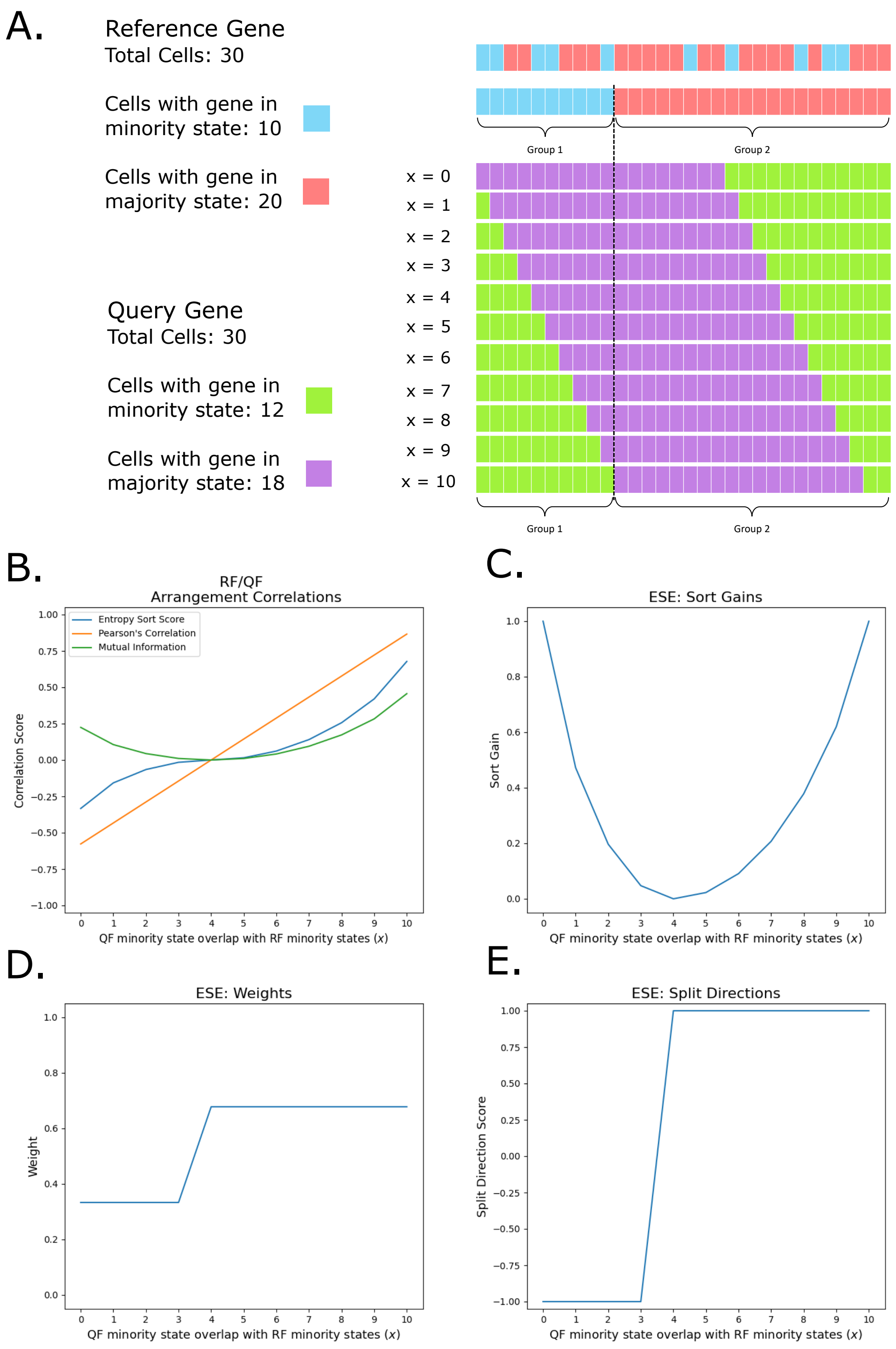


**Figure S1. Correlation Comparisons. A.** A simple example where the constants of the ESE (Fig. 1B) are fixed such that $G_1 = 10$, $G_2 = 20$ and $QF_m = 12$. We then inspect 10 different arrangements of the QF such that every possible value of $x$ (the overlap between the RF and QF minority states) is observed for the given system. **B.** ESS, Pearson's Correlation and Mutual Information scores for each RF/QF pair in A. **C-E.** The sort gain ($SG$), weights ($SW$) and split directions ($SD$) for each RF/QF pair in A..

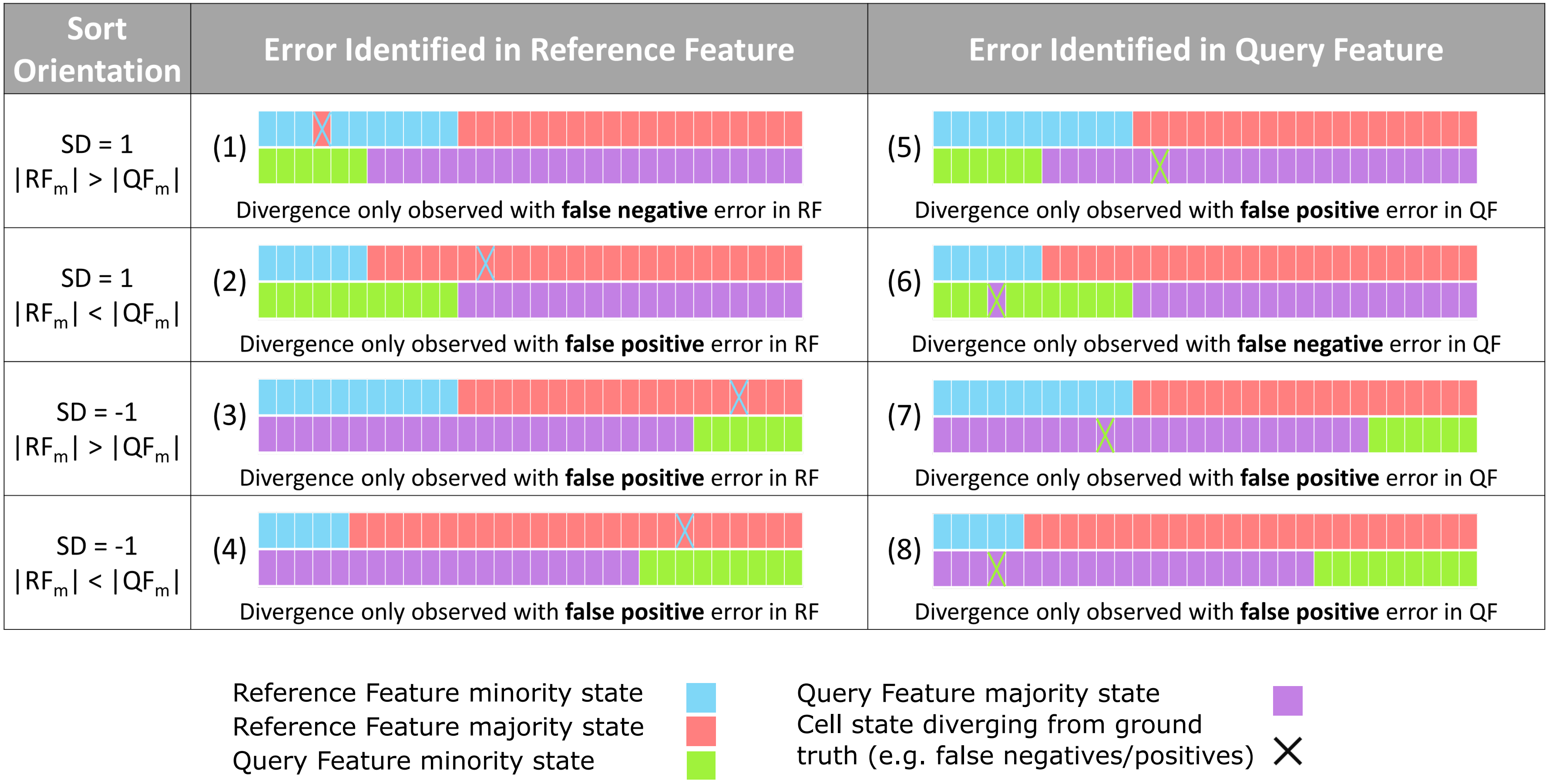


**Figure S2.** **Observable Divergence Error Scenarios.** Here we present the eight scenarios in which the introduction of error would lead to observable divergence on an ESE parabola. These scenarios are initially separable by whether the system has a SD of 1 or -1, and whether the number of QF minority states ($|QF_m|$) is greater than or less than the number of RF minority states ($|RF_m|$), as shown in the sort orientation column. These scenarios are subsequently defined by whether the error occurs in the RF or QF. For each scenario we show a basic example where the error would lead to observable divergence.

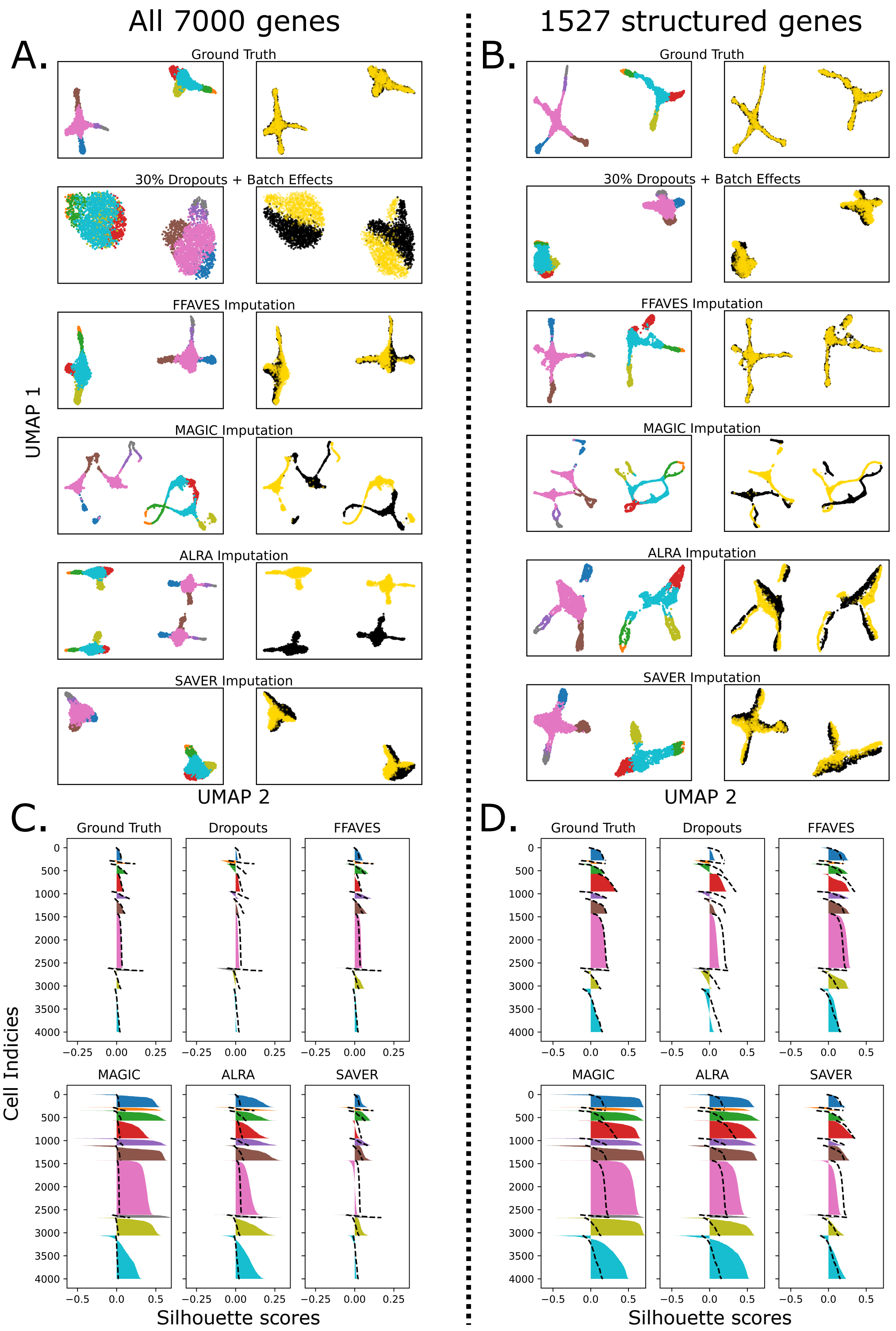


**Figure S3. Imputation comparisons on Dyngen simulated data containing 4000 cells and 7000 genes. Related to Fig 5.** We simulated a single cell RNA sequencing dataset with 4000 cells and 7000 genes using the Dyngen simulation software (Cannoodt et al. 2021). Of the 7000 genes, 1527 were highly structured genes part of the gene regulatory networks used to simulate cell types. 2878 of the 7000 genes were simulated as part of the house keeping gene regulatory networks and the final 2595 were genes ubiquitously randomly expressed throughout the cells. For the workflow used to create this dataset, see our online data repository. UMAPs of the synthetic data before and after imputation show that FFAVES facilitated imputation performs favourably compared to MAGIC, ALRA and SAVER when considering all 7000 genes **A.**, and just the 1527 structured genes **B.** Left hand columns of the UMAPs are coloured by 10 clusters of cells identified through K-means clustering. Right hand columns show the two batches created through batch specific simulated dropouts. Silhouette plots for 10 distinct clusters of cells, quantify how well the cells cluster together after imputation, compared to the ground truth dataset for all 7000 genes **C.** and just the 1527 structured genes **D.** Dashed lines in the silhouette plots trace the ground truth silhouette scores.

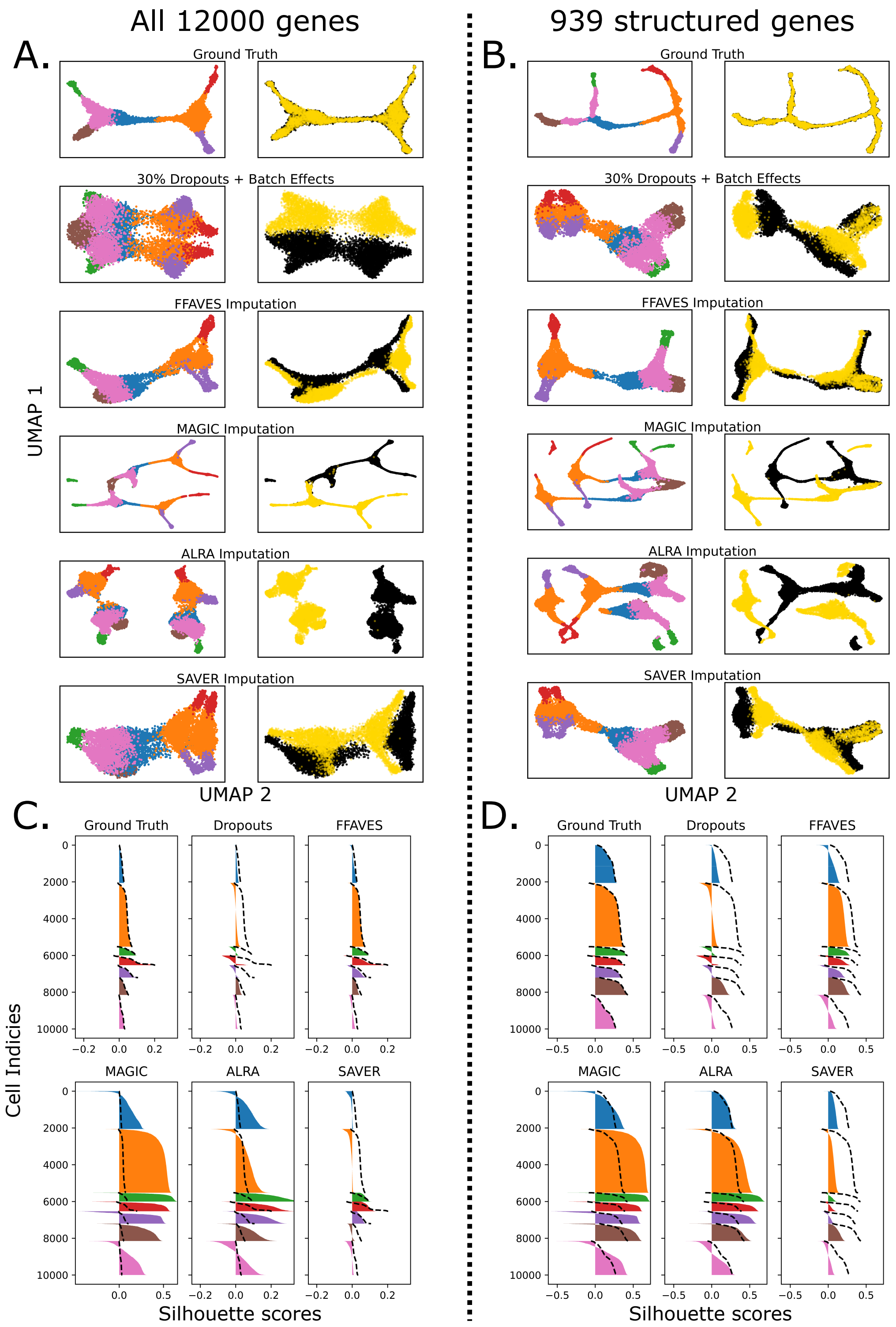


**Figure S4.** **Imputation comparisons on Dyngen simulated data containing 10000 cells and 12000 genes. Related to Fig 5.** We simulated a single cell RNA sequencing dataset with 10000 cells and 12000 genes using the Dyngen simulation software (Cannoodt et al. 2021). Of the 12000 genes, 996 were highly structured genes part of the gene regulatory networks used to simulate cell types. 4975 of the 12000 genes were simulated as part of the house keeping gene regulatory networks and the final 6029 were genes ubiquitously randomly expressed throughout the cells. For the workflow used to create this dataset, see our online data repository. UMAPs of the synthetic data before and after imputation show that FFAVES facilitated imputation performs favourably compared to MAGIC, ALRA and SAVER when considering all 12000 genes **A.**, and just the 996 structured genes **B.** Left hand columns of the UMAPs are coloured by 7 clusters of cells identified through K-means clustering. Right hand columns show the two batches created through batch specific simulated dropouts. Silhouette plots for 7 distinct clusters of cells, quantify how well the cells cluster together after imputation, compared to the ground truth dataset for all 12000 genes **C.** and just the 996 structured genes **D.** Dashed lines in the silhouette plots trace the ground truth silhouette scores.

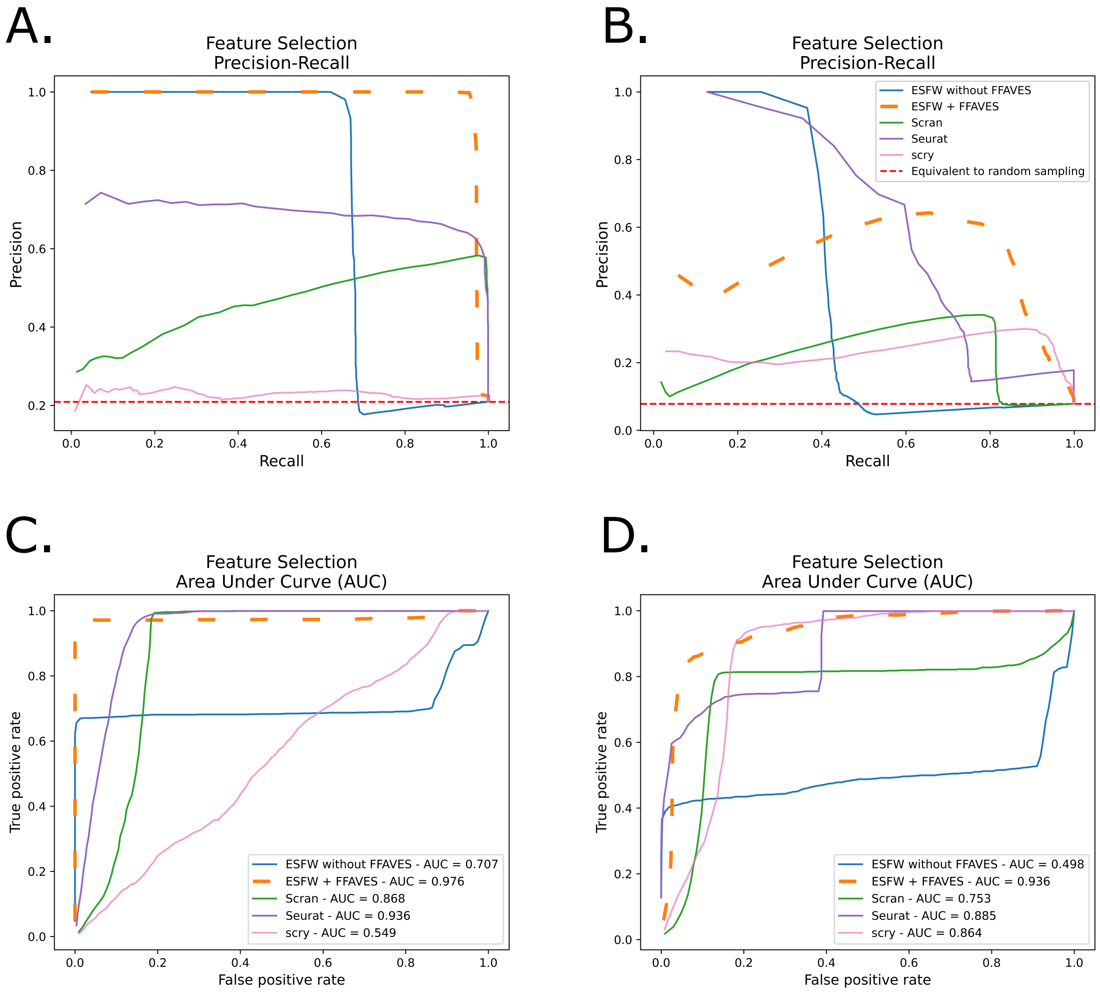


**Figure S5. Comparison of feature selection methods on Dyngen simulated datasets. Related to Fig 5.** Feature selection using FFAVES + ESFW was compared against Scran, Seurat and scry HVG feature selection software, when applied to the Dyngen synthetic scRNA-seq datasets presented in figures S3 and S4. The precision/recall curves are presented for these datasets in **A.** and **B.** respectively. We note that for our second Dyngen simulated dataset, ESFW without FFAVES does not outperform Seurat in terms of feature selection. ESFW + FFAVES initially appears to perform poorly. This is due to a small number of housekeeping genes having their signal falsely amplified by FFAVES. Once FFAVES + ESFW reaches a recall value of around 0.8 is it enriching highly structured genes considerably more than all other methods. However, to further emphasise that ESFW + FFAVES outperforms the other feature selection methods, we also provide the AUC curves to quantify feature selection performance in **C.** for the Dyngen simulated data in Fig. S3 and **D.** for the the data in Fig. S4. In AUC curves, a higher AUC indicates better performance. Generally AUC curves are not used when there is a significant class in balance (Davis and Goadrich n.d.). Despite there being a high degree of class imbalance in the Dyngen simulated data, since there are far more uninformative genes than structured genes, the AUC curves are useful for further validating the ESFW + FFAVES performance.

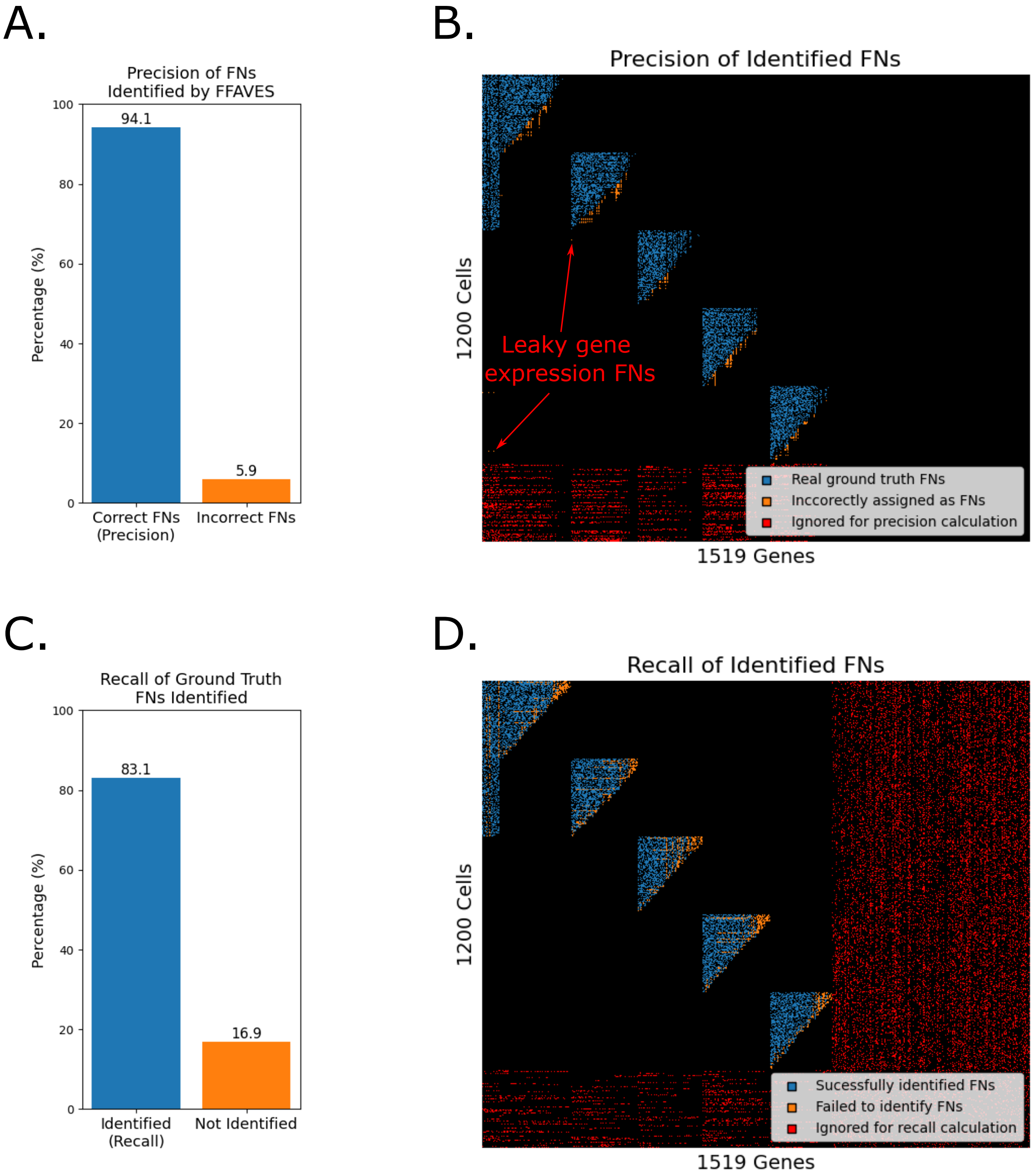


**Figure S6.** **Precision and Recall for identifying FN data points in the synthetic data. Related to Fig. 4C. A.** Percentage of FNs identified by FFAVES that are correctly identified in the synthetic data according to the known ground truth. **B.** Visualisation of where the correctly (blue) and incorrectly (orange) FNs identified by FFAVES reside in the expression matrix. **C.** Percentage of all ground truth FNs that were correctly identified by FFAVES according to the known ground truth. **D.** Visualisation of all successfully (blue) and unsuccessfully (orange) identified FNs. Red data points in b and d are FNs that were added to the ground truth but are not considered FNs because adding random FN error to an already random distribution (the randomly expressed genes), is equivalent to a gene with random gene expression but with lower mean expression.

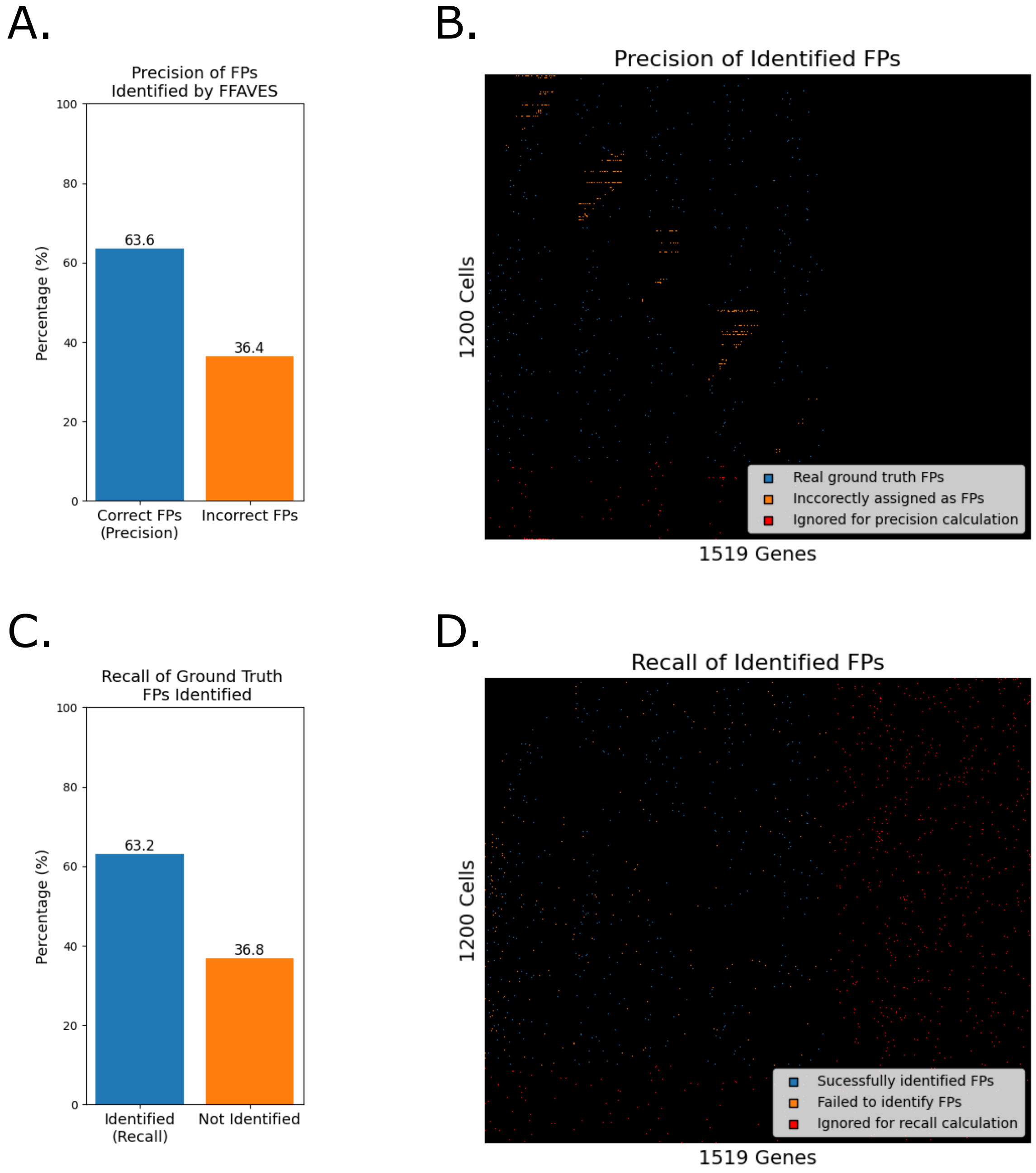


**Figure S7. Precision and Recall for identifying FP data points in the synthetic data. Related to Fig. 4C. A.** Percentage of FPs identified by FFAVES that are correctly identified in the synthetic data according to the known ground truth. **B.** Visualisation of where correctly (blue) and incorrectly (orange) FPs identified by FFAVES reside in the expression matrix. **C.** Percentage of all ground truth FPs that were correctly identified by FFAVES according to the known ground truth. **D.** Visualisation of all successfully (blue) and unsuccessfully (orange) identified FPs. Red data points in b and d are FPs that were added to the ground truth but are not considered FPs because adding random FP error to an already random distribution (the randomly expressed genes), is equivalent to a gene with random gene expression but with higher mean expression.

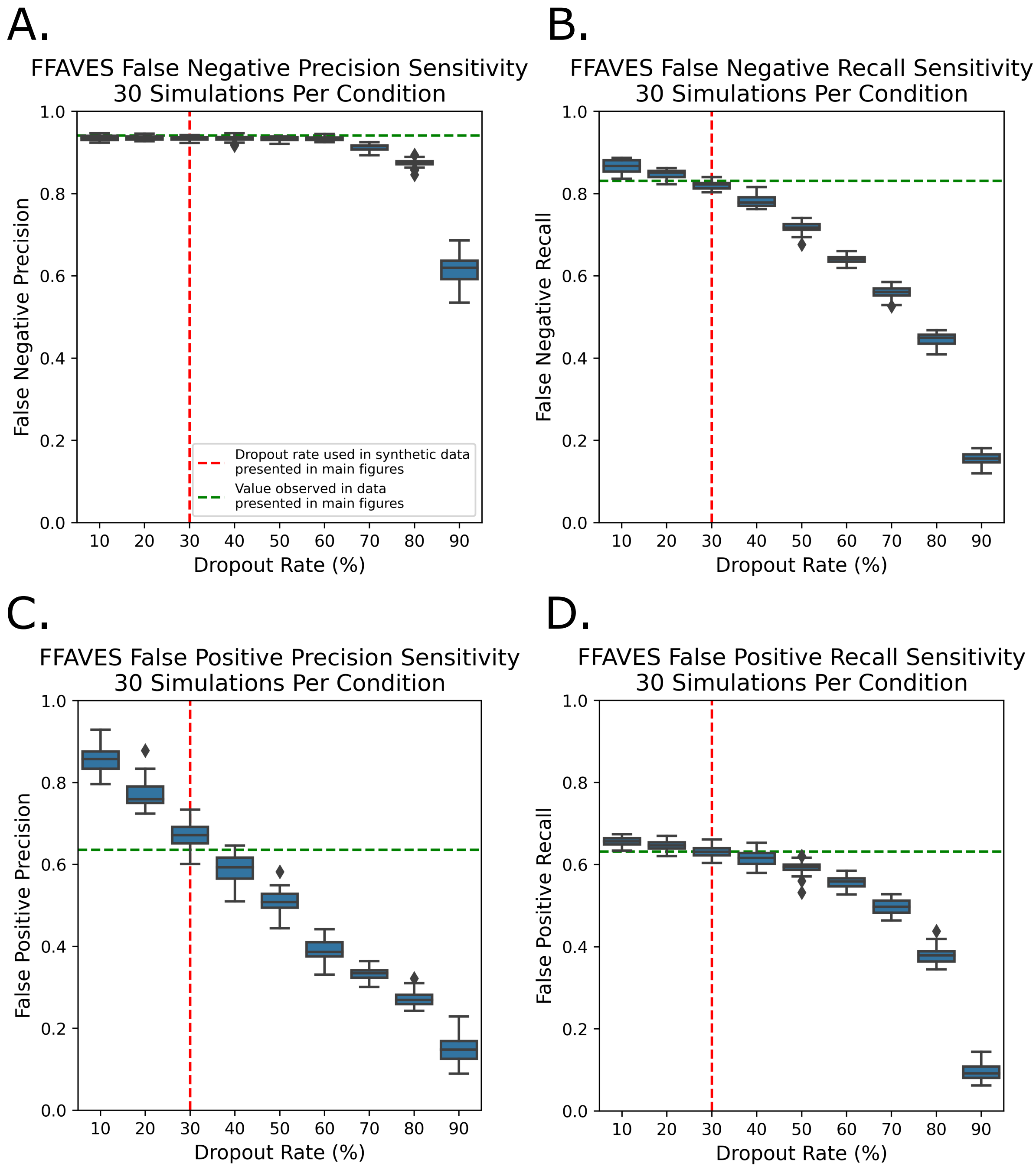


**Figure S8. False negative and false positive precision and recall scores are robust to stochastically generated datasets across varying dropout rates. Related to Fig. 4C, D.** To verify that FFAVES is robust to stochastically generated synthetic data, we created 30 new datasets in the same manner as was done for the synthetic data presented in the main text (SI 8) and calculate the precision/recall scores for the FPs/FNs identified by FFAVES. This process was repeated while varying the intentionally added dropout rates from 0.1-0.9. The results for each dropout rate are summarised in box plots. Diamond points are data points outside of the interquartile range range. Red dashed lines indicate the dropout rate that was used to generate the synthetic data presented in the main text (30%). Green dashed lines show the scores that were calculated in the data presented in the main text for, **A.** FN precision, **B.** FN recall, **C.** FP precision, and **D.** FP recall.

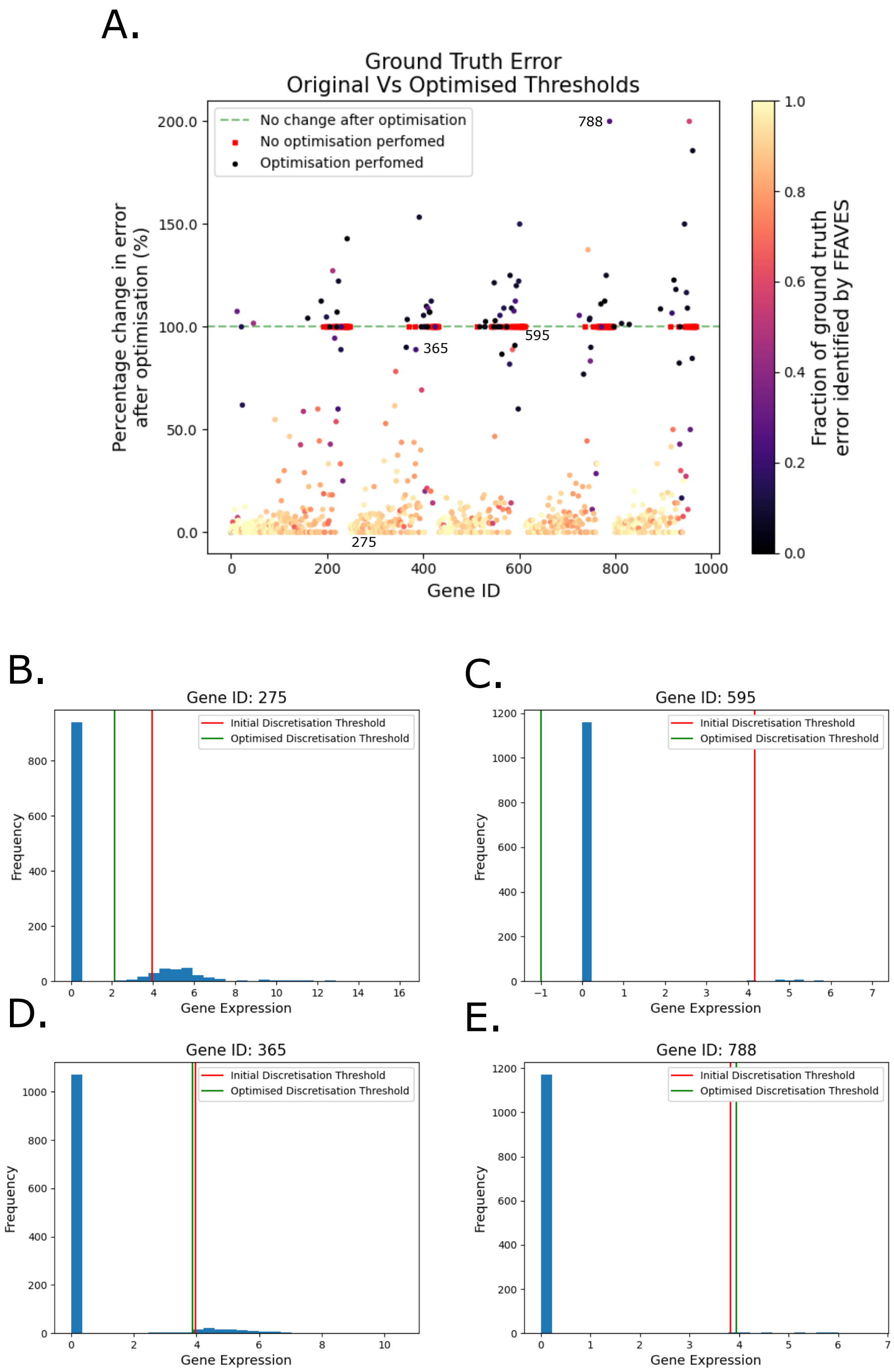


**Figure S9.** **FFAVES accurately performs discretisation threshold optimisation. Related to Fig. 4G. A.** Percentage change in the number of erroneous data points (FNs + FPs) when discretising $SD1_{GT}$ with the sub-optimal discretisation thresholds used to create $M$ from $SD1_N$ vs. the optimised thresholds identified after FFAVES has corrected for FN/FP data points. Each point in the plot corresponds to one of the 969 highly structured genes of SD1. The IDs of the genes shown in panels B-E are annotated on the the plot in A.. **B-E.** Gene expression histograms showing the initial sub-optimal discretisation threshold used to create $M$ from $SD1_N$ (red), and the optimised discretisation threshold identified after application of FFAVES (green). **B.** Threshold optimisation successfully identified the threshold that separates active/inactive gene states according to our ground truth, where any value greater than 0 is designated an active expression state. **C.** No FN/FP data points were identified so no optimisation could occur. **D.** Optimisation led to only a small reduction in erroneous discretisation. **E.** Optimisation led to an increase in erroneous discretisation.
