## Supplemental figures 10-16 for "Entropy sorting of single cell RNA sequencing data reveals the inner cell mass in the human pre-implantation embryo"

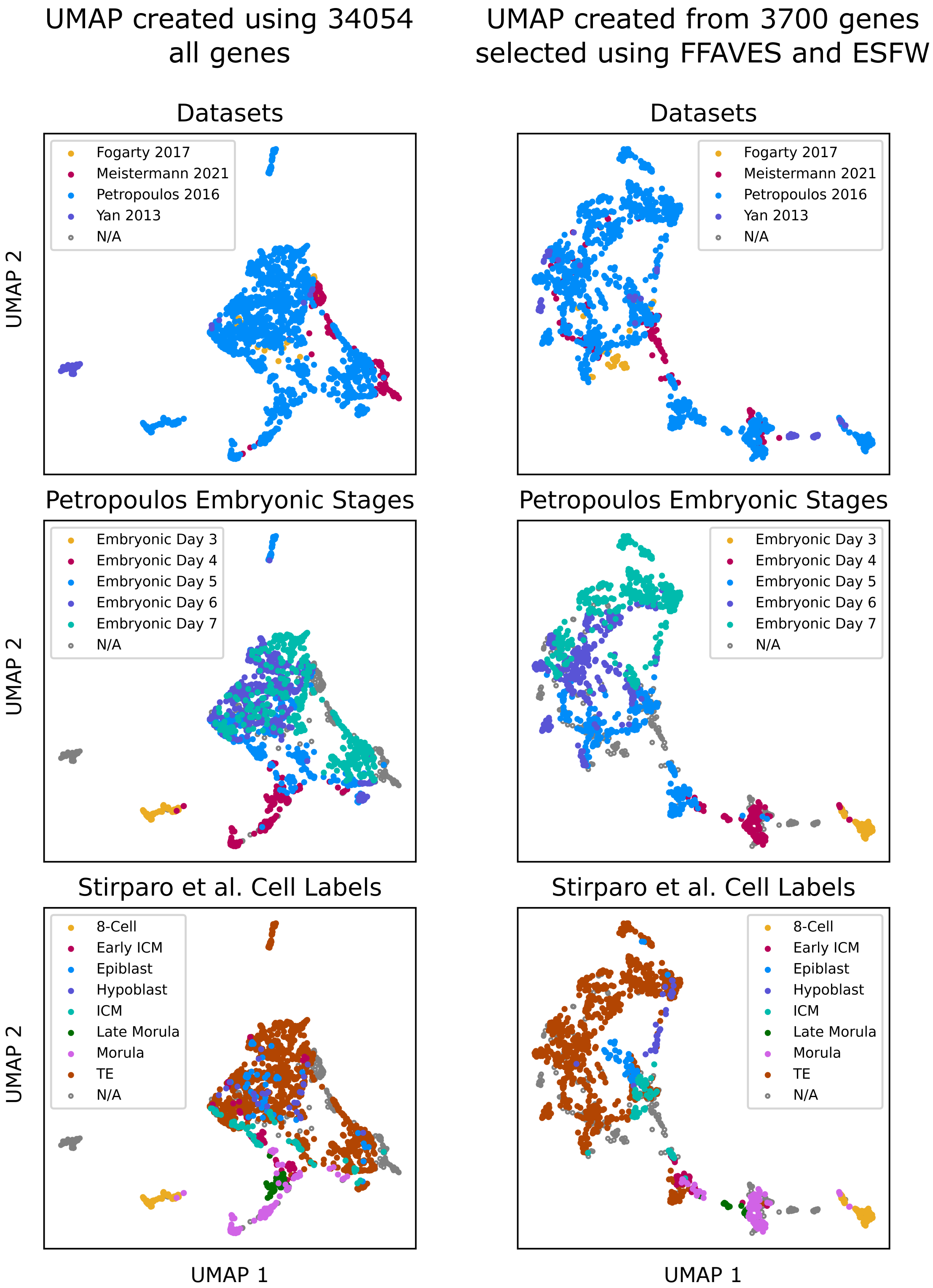


**Figure S10. Sub-setting to the 3700 genes identified by FFAVES + ESFW reveals a high resolution description of human pre-implantation embryo development. Related to Fig. 6A.** Comparison of UMAPs created using all 34054 genes in the original dataset vs. the 3700 highly structured genes identified by FFAVES + ESFW. Both embeddings were created using the counts matrices without any data transformations, imputation or smoothing. The lack of data augmentation demonstrates that a large potion of batch effects and noise can be reduced simply by removing randomly expressed/uninformative genes.

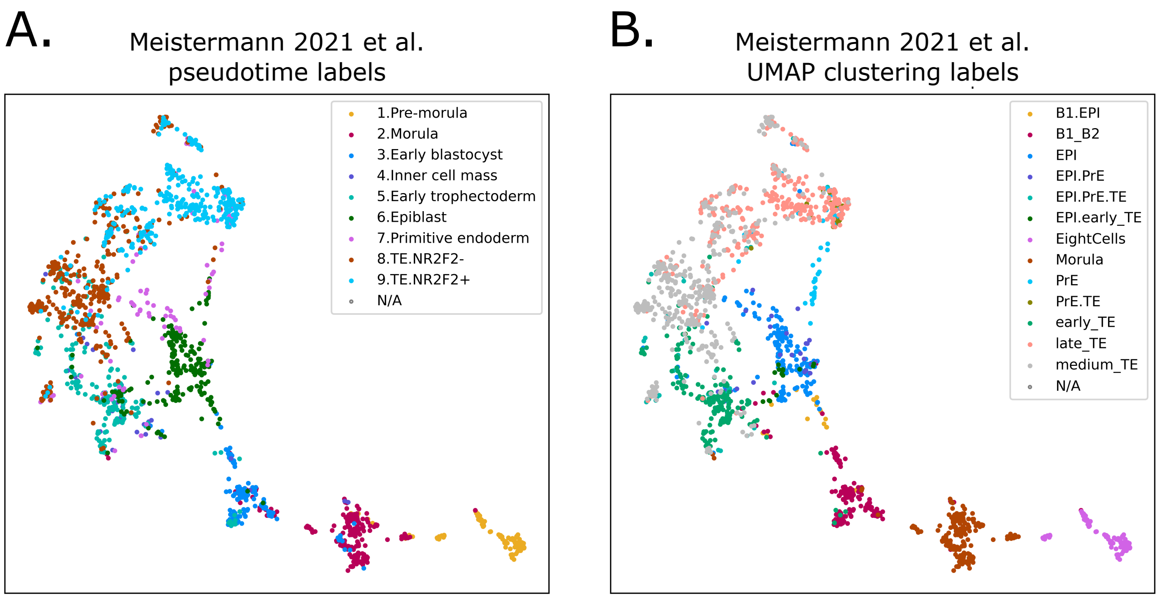


**Figure S11. The Meistermann et al. 2021 of human pre-implantation embryo scRNA-seq data are unable to resolve an ICM population. Related to figure 6.** In their analysis, Meistermann et al. 2021 note that they were unable to isolate a distinct ICM population. By overlaying their assigned labels from their pseudotime analysis (**A.**) and their UMAP clustering analysis (**B.**), we find that our unsupervised analysis of the scRNA-seq data largely agree with their supervised analysis. However our unsupervised UMAP suggests that in their pseudotime analysis, the epiblast has been incorrectly labeled as the hypoblast (primitive endoderm) and their suggest epiblast population is our suggested inner cell mass cluster. Likewise in their UMAP clustering, their analysis appears to have been unable to separate our proposed inner cell mass cells from the epiblast cells.

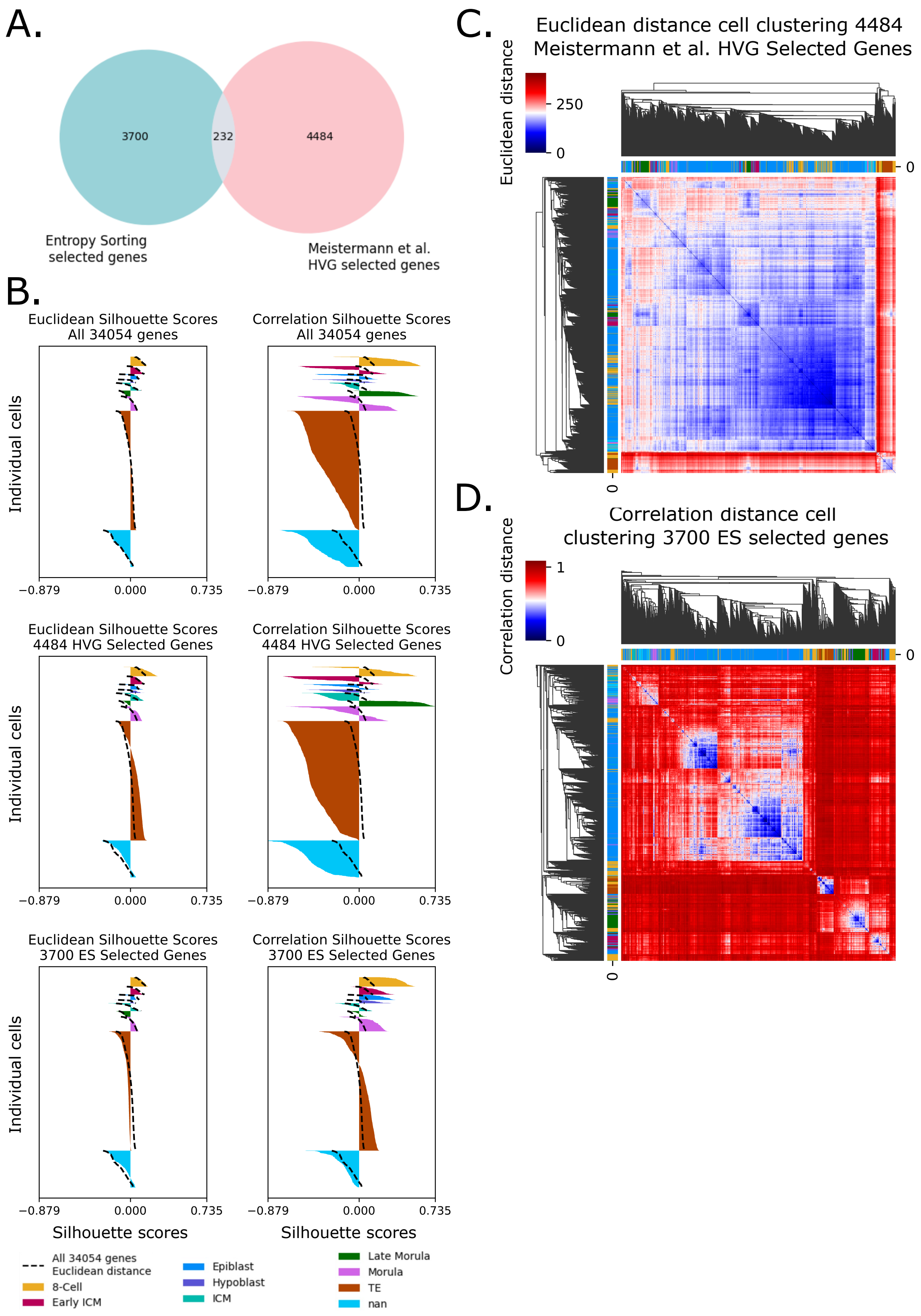


**Figure S12.** **FFAVES + ESFW selected genes produce a higher resolution of specific cell type clusters than HVG selection in the human pre-implantation embryo data. A.** Overlap of genes selected as informative for cell identity in the Meistermann et al. 2021 human pre-implantation embryo dataset through Entropy Sorting verses through highly variable gene (HVG) selection by Meistermann et al. 2021. **B.** Silhouette plots comparing the clustering performance of cells into the cell identities determined by Stirparo et al. 2018 through supervised analysis. The silhouette scores obtained when using the 3700 ES selected genes and a correlation distance metric are consistently higher than all other cases, indicating that the cells have been more confidently assigned to the cell type labels given by Stirparo et al. 2018. **C.** Heatmap of cell similarities when using the HVG selected genes and euclidean distance. **D.** Heatmap of cell similarities when using the ES selected genes and correlation distance. C. and D. are a visual representations of how cell type identities can become more distinct when using the ES selected genes. The full workflow for choosing the ES highly structured genes and discussion around using euclidean or correlation distance metrics can be found in our online methods.

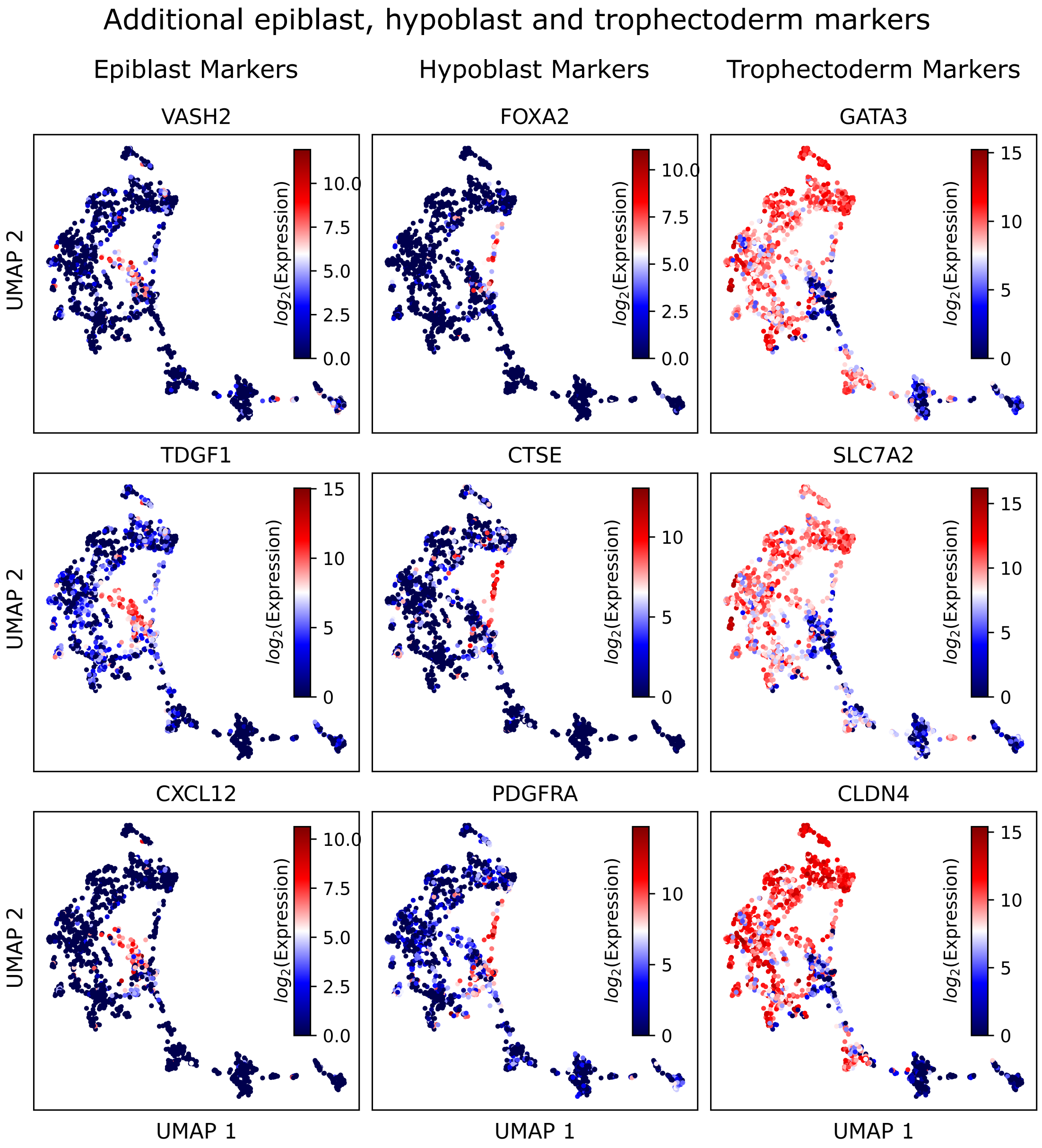


**Figure S13.** **Additional epiblast, hypoblast and trophectoderm cell type marker expression profiles. Related to Fig. 6B.**

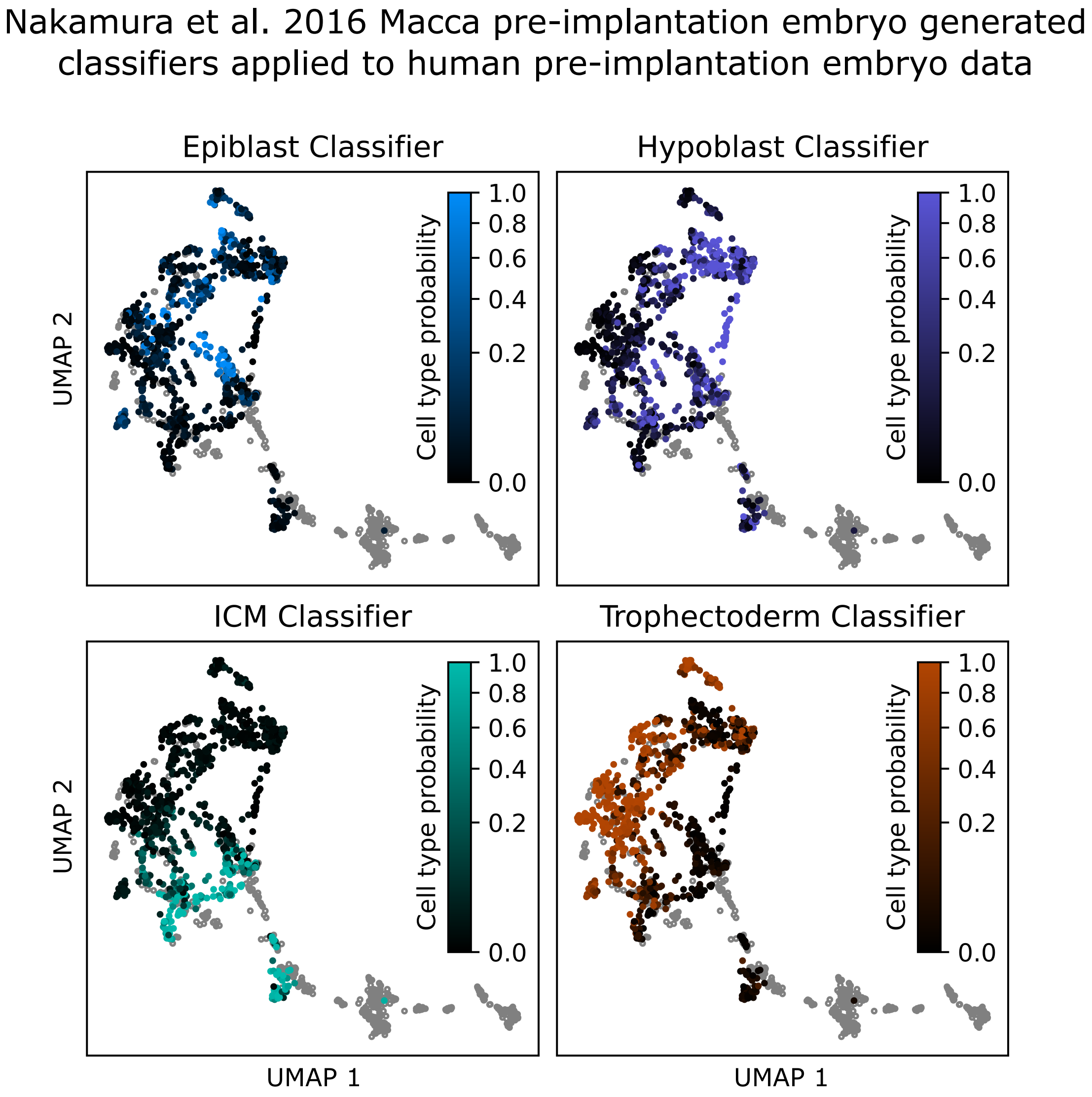


**Figure S14.** **Predicted cell type probabilities of individual cells when a classifier trained on Macaca primate pre-implantation embryo scRNA-seq data from the independent Nakamura et al. dataset. Related to Fig. 6C.** Grey samples are those that were not processed by the classifier to avoid confounding variables such as batch effects.

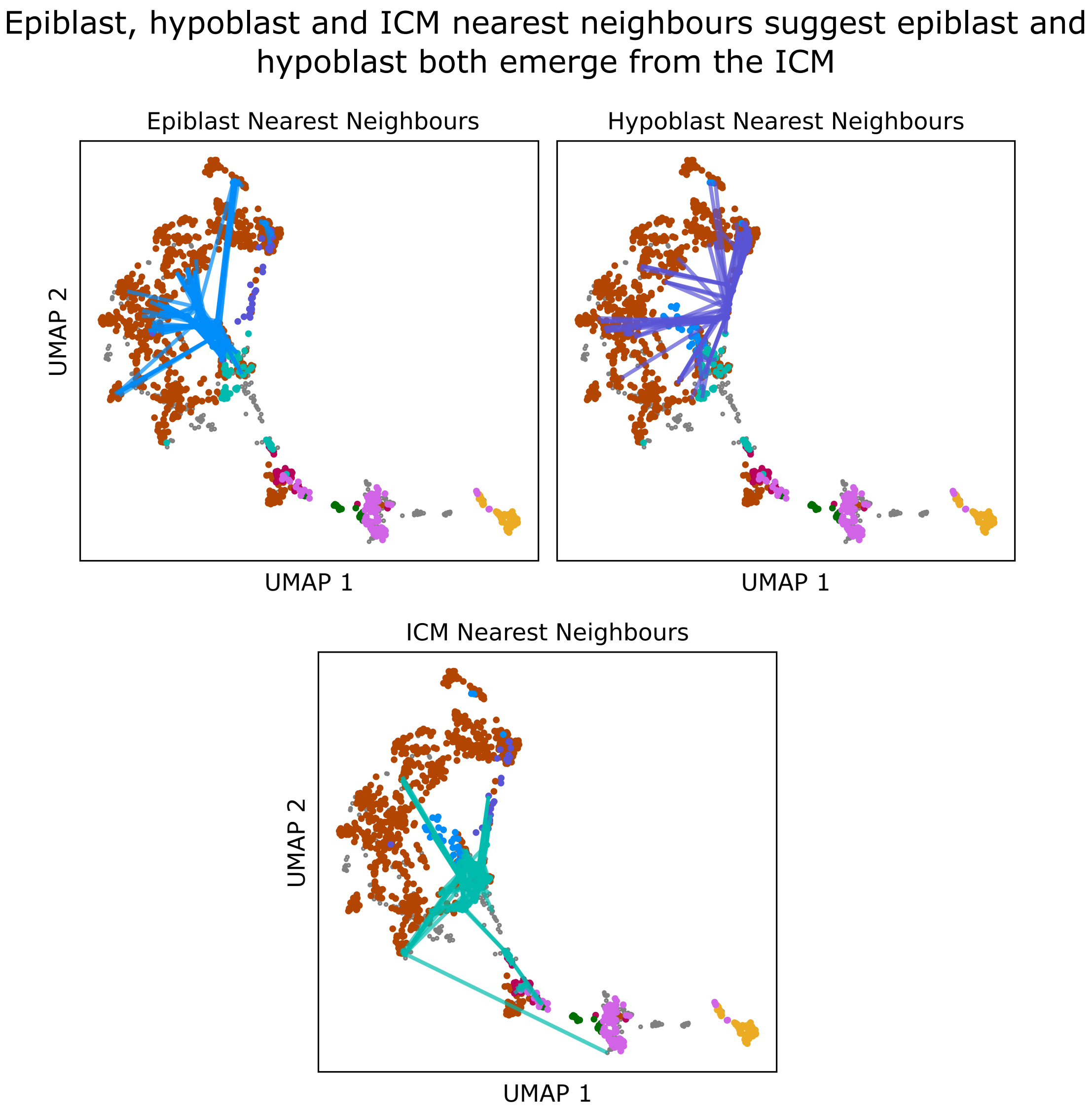


**Figure S15. Separate nearest neighbour embeddings for epiblast, hypoblast and ICM cells indicate that the epiblast and hypoblast gene expression signatures are more similar to the ICM than to each other. Related to Fig. 6D.** Each of the epiblast, hypoblast and ICM cells identified by Stirparo et al. is connected by lines to their 10 most similar samples according to gene expression.

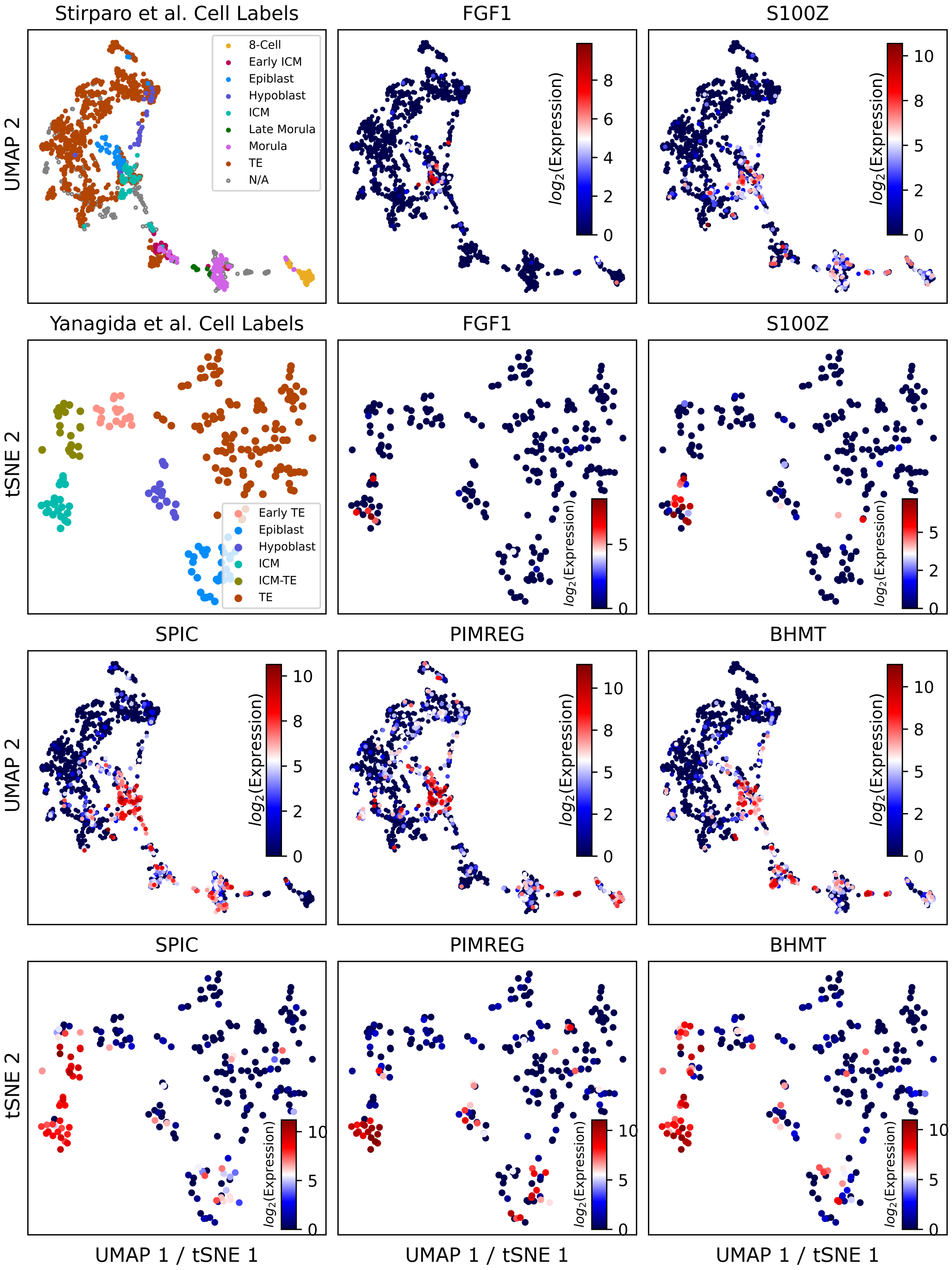


**Figure S16. Additional potential ICM marker expression profiles. Related to Fig. 7.** Potential ICM markers were selected based on their localised expression in the ICM population of our FFAVES + ESFW embedding that is corroborated in the UMAP embedding generated by Yanagida et al. on their own independent human pre-implantation embryo scRNA-sequencing data.
