## Supplemental information for "Entropy sorting of single cell RNA sequencing data reveals the inner cell mass in the human pre-implantation embryo"

### Supplementary Information

#### SI 1 Discretisation of Continuous Data

A requirement for applying ES is that the data be discrete. We arbitrarily choose to represent these distinct states as 0's or 1's. How to discretise continuous gene expression data can be non-trivial. Consider the hypothetical gene expression profiles in Fig. SI 1. In the bimodal scenario (Fig. SI 1A), there are two distinct populations of gene expression. A large proportion of the cells display an expression value of 0, indicating that the gene is inactive. A second population display a normal distribution of expression with mean = 5 and standard deviation = 1. Hence, it could be considered reasonable that any data point with a value greater than zero corresponds to an active state and should be discretised to 1 (e.g. a discretisation threshold of 1, as indicated by the green dashed line, achieves this discretisation). However, from a biological standpoint, we may be sceptical as to whether low non-zero expression values represent a functionally active expression state. If they do not, we should discretise them to 0 to signify an inactive state (for example, using a discretisation threshold of 3.5, indicated by the dashed red line). The problem is further confounded if you consider multimodal data, for example if a gene is considered to have distinct functionality at more than two distinct expression states. In this case, as illustrated in Fig. SI 1B, we may wish to distinguish the different functionalities of the expression states defined by threshold 2 and threshold 3 (red and yellow dashed lines, respectively).

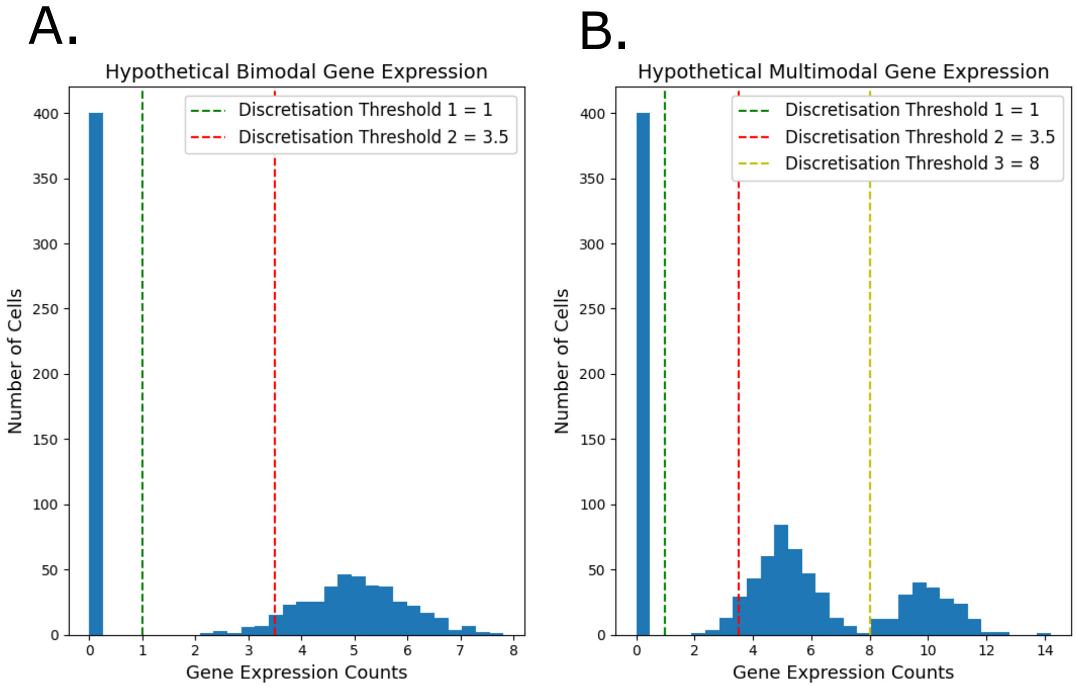


**Figure SI 1. Choosing discretisation thresholds for continuous data. A.** Example of a bimodal distribution for a given feature from continuous data with two possible suggested thresholds for discretisation. **B.** Example of a multimodal distribution for a given feature from continuous data with three possible suggested thresholds for discretisation.

Although it is a requirement that ES be presented with discrete data, this abstraction of the data can be justified in many scenarios. The strength of ES is the ability to identify discrete functional relationships between features in high dimensional data, rather than correlative patterns on the continuous scale. This is often a tractable and useful task. In the context of gene expression, we often talk about a gene being active or inactive, or being functionally related to other genes in the data, without specifically being interested in the raw expression values. Often we are simply interested in whether a gene's expression is distinctly different in one context versus another. To further emphasise that FFAVES and ESFW are tools to aid interpretability of data, having identified functional relationships between features in a data set using FFAVES/ESFW, we recommend that if the data was originally in a more complex format (i.e. a continuous scale), to subsequently return to this modality and use the information garnered through FFAVES/ESFW to enhance analysis.

How best to discretise the data is context dependent and left to the user. However we can simplify the procedure by demonstrating that as long as the discretisation is achieved reasonably well, and not necessarily perfectly, FFAVES has the capacity to automatically adjust and optimise the discretisation threshold of each feature based on the prevailing structure in the data (Fig. 4g, Fig. S9).

For now, consider the bimodal example presented in Fig SI 1A. Either of Threshold 1 or 2 constitute reasonable levels to discretise the data, depending on assumptions. Let us now assume that we chose Threshold 2, but the ground truth of the data actually agrees with Threshold 1. In this scenario, we would assign every data point with a value greater or equal to 3.5 the expression state 1 and every observation less than 3.5 an expression state of 0. Since the ground truth is that all values greater than 0 constitute a functionally active expression state, it follows that all values less than 3.5 but greater than 0 are FNs. Hence, when we perform ES on this feature, the FN data points will be quantified as divergent. If there is enough evidence for this divergence to be statistically significant, the expression states of the FN points will be switched from 0's to 1's. This is equivalent to lowering the discretisation threshold. If this process were to perfectly adjust the threshold, FFAVES would automatically adjust the threshold from Threshold 2 down to Threshold 1.

Finally, to address the issue of multimodality, consider the example of a feature displaying a multimodal expression distribution from Fig. SI 1B. There are two perspectives from which FFAVES can be found to deal with multimodal functionality. First, while there may be scenarios where FFAVES will struggle to directly elucidate the subsets of features that are related to separate functional modalities for a feature of interest, the resulting substructure can still be identified. For example, we could reasonably hypothesise that there exists a set of genes ($S_1$) that are functionally activated when the feature has an expression value between 1 and 8 (red line to yellow line). We could then also postulate that there is a second set of genes ($S_2$) that only become functionally active when the expression of the given gene is greater than 8 (above yellow line). If we were to discretise the given gene at a threshold of 1, then it is unlikely ES would identify the genes that are members of $S_1$ or $S_2$. This is because their functionally active states would span distinctly different regions of the data. However, this issue is mitigated by the fact that if the genes within $S_1$ are functionally related to each other in a distinct region of the data, ES will identify these genes as forming a distinct functional module. Likewise for $S_2$. Hence, although FFAVES would fail to identify that the genes within $S_1$ or $S_2$ are functionally related to distinct modalities of the multimodal feature in Fig. SI 1B, the information regarding the genes within $S_1$ and $S_2$ existing in a distinct region of the data is still captured. This process is exemplified in the synthetic data within the main body of this paper through the 50 multimodal genes (Fig. 4a).

An alternative method for discretisation exists, which may be desirable if a user is confident that features in the data exhibit functional multimodal activity, and they would like to identify these cases. In this scenario, the user may wish to create discrete 'pseudo-features' before inputting them into FFAVES. In Fig. SI 1B this could be achieved by turning the multimodal feature into 3 new discrete features. The first pseudo-feature ($PF_1$) could have all values greater than 1 discretised to 1 and all other values equal to 0. The second pseudo-feature ($PF_2$) could have all values between 1 and 8 discretised to 1 and all other values equal to 0. Finally, the third pseudo-feature ($PF_3$) could have all values greater than 8 equal to 1 and all other values equal to 0. Now ES would find features that become functionally active whenever the feature of interest is expressed with a value greater than 0 as being functionally related to $PF_1$. Likewise the genes within $S_1$ and $S_2$ would be identified as being functionally related to $PF_2$ and $PF_3$ respectively. Once again, if such a threshold technique was carried out reasonably well, FFAVES should be able to automatically optimise the thresholds used to create each pseudo-feature. However, it should be noted that seeking higher resolution in this manner for many features could dramatically increase the number of observed features in a given data set and thereby increase the computational run time.

#### SI 2 Deriving the Entropy Sort Equation

In the following we derive the Entropy Sort Equation (ESE). To do so we use the toy example outlined in Fig. SI 2, in which the expression profile of 30 cells is measured, and consider the activity of two genes. We seek to quantify to what degree the expression of one gene correlates with the expression of the other. We treat this as a sorting problem, and first discretise the expression of each gene (Fig. SI 2A). In our example, if we inspect either Gene 1 or Gene 2, we can calculate their Shannon Entropy (Shannon 1948), $H$, as follows:

$$H = \sum_{i=1}^{n} -p_i \log_2(p_i), \tag{1}$$

where $p_i$ is the probability of selecting a particular expression state $i$ in the samples and $n$ is the number of unique states (here, $n = 2$ since there are two discrete states). For Gene 1, 10 cells display Gene 1 as active and 20 display Gene 1 as inactive. Hence, the entropy for Gene 1 is given by

$$H_{Gene\ 1} = -\frac{10}{30}\log_2(\frac{10}{30}) - \frac{20}{30}\log_2(\frac{20}{30}) = 0.918. \tag{2}$$

Likewise, the entropy for Gene 2 is calculated to be

$$H_{Gene\ 2} = -\frac{8}{30}\log_2(\frac{8}{30}) + -\frac{22}{30}\log_2(\frac{22}{30}) = 0.837. \tag{3}$$

The Shannon Entropy of a given variable describes the average amount of “information" or “uncertainty" inherent in the variable’s possible outcomes. Up to this point Gene 1 and Gene 2 are considered as two independent variables. To extend Shannon Entropy to quantify the relationship between the two genes, we set up the following hypothesis. We assume that splitting the data into two groups based on the expression of Gene 1 is a perfect partition of disorder in the system. That is, if we group all cells for which Gene 1 is active and then separately group all the cells for which Gene 1 is inactive, the expression states of every other gene in the data would also be perfectly sorted into these two groups. We then test how true this is for the other genes in the data, i.e. Gene 2 (Fig. SI 2B).

We designate group 1 ($G_1$) to contain those cells in which Gene 1 is active, and group 2 ($G_2$) to contain those cells in which it is inactive. This minimises the entropy of the system to zero, since both groups are homogeneous. We now maintain the Gene 1 groupings while inspecting the expression states of Gene 2. As shown in Fig. SI 2B, these groups fail to perfectly sort the expression states of Gene 2. To quantify this, we calculate the entropies of $G_1$ and $G_2$ with regards to Gene 2. For $G_1$, there are 10 cells in total, 5 of which display Gene 2 as active and 5 display Gene 2 as inactive. Therefore,

$$H_{G_1} = -\frac{5}{10}\log_2(\frac{5}{10}) + -\frac{5}{10}\log_2(\frac{5}{10}) = 1.00. \tag{4}$$

Likewise for $G_2$, since 3 cells show Gene 2 as active and 17 as inactive, the entropy is given by

$$H_{G_2} = -\frac{3}{20}\log_2(\frac{3}{20}) + -\frac{17}{20}\log_2(\frac{17}{20}) = 0.610. \tag{5}$$

It follows that the total entropy of the system is 1.610. To account for the different cardinalities of $G_1$ and $G_2$ we add a weight term for each calculation. In doing so we acknowledge that the entropy of $G_1$ and $G_2$ are portions of the entropy of the entire data set, and ensure that entropy of the system will always be a value between 0 and 1. Hence,

$$\text{Gene 2 System Entropy} = \frac{|G_1|}{C_T}H_{G1} + \frac{|G_2|}{C_T}H_{G2} = \frac{10}{30} * 1 + \frac{20}{30} * 0.610 = 0.740. \tag{6}$$

$C_T$ designates the total number of cells in the system ($C_T = 30$) and $|G_1|$ and $|G_2|$ are the cardinalities of $G_1$ and $G_2$ respectively.

Intuitively, Eqn 6 corresponds to the conditional entropy of Gene 2 given Gene 1. In other words, given the expression states of Gene 1, it defines the uncertainty around the expression states of Gene 2 in any particular cell. Typically conditional entropy is considered as purely probabilistic, resulting in a single value between 0 and 1 that quantifies the dependency between two or more random variables. We seek a similar quantification of the relationship between Gene 1 and 2, but as a sorting problem. To achieve this, we note that there are optimal arrangements where the entropy of the system defined by the expression states of two genes can be minimised (Fig. SI 2C). To minimise the conditional entropy of an observed system between two genes we must first constrain the system around one of the genes, Gene 1. The expression states of Gene 1 define two groups of fixed size. We then inspect Gene 2, and permute the locations of the expression states of Gene 2 into an optimal arrangement that minimises the entropy of the system (Fig. SI 2C). As with the expression states of Gene 1, the number of active/inactive states

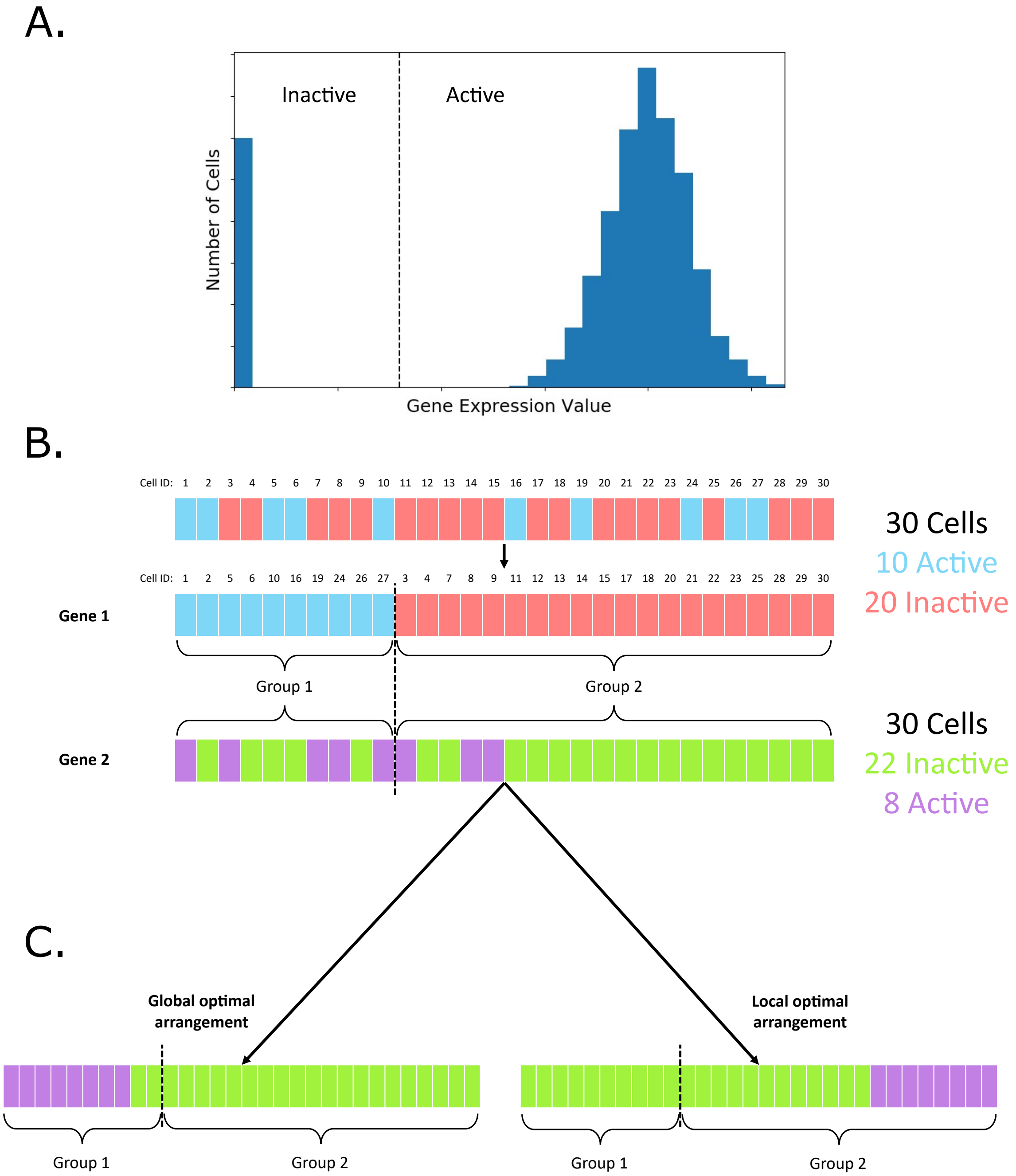


**Figure SI 2. Pairwise gene correlation as a sorting problem. A.** Illustrative histogram of bimodal gene expression across a population of cells. Continuous expression data must be discretised into two states. This can be performed by identifying a threshold (vertical dashed line) above which values are considered as active, and below which they are considered inactive. **B.** Example discrete expression state profiles for 30 cells. We initially inspect Gene 1 and rearrange the cells such that all the active states are sorted together into group 1, and all inactive states are in group 2. We then inspect a second gene, Gene 2, and observe the expression states of gene 2 for each cell. **C.** There are two theoretical optimal sorts where the heterogeneity of group 1 or group 2 (as defined by Gene 1) are minimised when inspecting the states of Gene 2.

of Gene 2 is fixed to the value initially observed in the data, and it is simply their abundance within $G_1$ and $G_2$ that may be changed.

To highlight that entropy sorting generalises beyond scRNA-seq data, moving forward we will refer to genes as features. We designate the feature that defines groups 1 and 2 as the Reference Feature (RF), and any feature that is subsequently inspected as the Query Feature (QF). We will now formulate a smooth function that describes a range of conditional entropies for any RF/QF pair.

First, we outline some terms to generalise the ESE to any system. In Fig. SI 2 we conveniently set up the system such that the number of cells with Gene 1 or Gene 2 active was smaller than the number of cells with the same gene inactive. To formulate the ESE as a sorting problem we can envision the sorting task as enriching the overlap of less commonly observed states, which in our example would be the active gene expression states. Alternatively, we could view the sorting problem as enriching the overlap of the more commonly observed states, but in either scenario the conditional entropy is the same. The practical difference is simply whether you choose to count how many of the less commonly observed states of the RF/QF pair overlap, or how many of the more commonly observed states overlap.

To this end, we employ notation to indicate whether the sample is displaying the less common or more common state for a given feature. We designate the more common state as the Majority (M) state, and the less common state the Minority (m) state. Hence, for any RF/QF pair, the minority state of the RF refers to those samples that form Group 1 ($G_1$) when partitioning the data and the majority state of the RF refers to samples that form Group 2 ($G_2$). Having partitioned the data via the RF, we can formulate terms describing how the QF is arranged within $G_1$ and $G_2$. Here, $QF_{m,G_1}$ and $QF_{M,G_1}$ correspond to the number of minority and majority QF states that occupy $G_1$, respectively. Similarly, $QF_{m,G_2}$ and $QF_{M,G_2}$ represent the number of minority and majority QF states in $G_2$.

We return to Eqn 4 and Eqn 5 to reform the calculations for the entropies of $G_1$ and $G_2$ such that

$$H_{G_1} = \frac{|G_1|}{|G_1|+|G_2|}\left(-\frac{QF_{m,G_1}}{|G_1|}\log_2\left(\frac{QF_{m,G_1}}{|G_1|}\right)-\frac{QF_{M,G_1}}{|G_1|}\log_2\left(\frac{QF_{M,G_1}}{|G_1|}\right)\right), \tag{7}$$

$$H_{G_2} = \frac{|G_2|}{|G_1|+|G_2|}\left(-\frac{QF_{m,G_2}}{|G_2|}\log_2\left(\frac{QF_{m,G_2}}{|G_2|}\right)-\frac{QF_{M,G_2}}{|G_2|}\log_2\left(\frac{QF_{M,G_2}}{|G_2|}\right)\right). \tag{8}$$

Equations 7 and 8 allow us to calculate the entropy of $G_1$ and $G_2$ based on the observed arrangement of any QF with respect to a given RF. The conditional entropy is then the sum of $H_{G_1}$ and $H_{G_2}$. The highlighted terms emphasise the dependent variables. We can re-write three of these variables such that they are each functions of ($QF_{m,G_1}$):

$$QF_{M,G_1} = |G_1| - QF_{m,G_1}, \tag{9}$$

$$QF_{m,G_2} = QF_m - QF_{m,G_1}, \tag{10}$$

$$\begin{aligned} QF_{M,G_2} &= |G_2| - QF_{m,G_2} \\ &= |G_2| - (QF_m - QF_{m,G_1}) \\ &= |G_2| - QF_m + QF_{m,G_1}. \end{aligned} \tag{11}$$

Here we have included a constant term, $QF_m$, which describes the total number of QF samples that display the QF minority state (regardless of $G_1$ or $G_2$ overlap). Substituting into (7) and (8),

$$H_{G_1} = \frac{|G_1|}{|G_1|+|G_2|}\left(-\frac{QF_{m,G_1}}{|G_1|}\log_2\left(\frac{QF_{m,G_1}}{|G_1|}\right)-\frac{|G_1|-QF_{m,G_1}}{|G_1|}\log_2\left(\frac{|G_1|-QF_{m,G_1}}{|G_1|}\right)\right), \tag{12}$$

$$H_{G_2} = \frac{|G_2|}{|G_1|+|G_2|}\left(-\frac{QF_m-QF_{m,G_1}}{|G_2|}\log_2\left(\frac{QF_m-QF_{m,G_1}}{|G_2|}\right)-\frac{|G_2|-QF_m+QF_{m,G_1}}{|G_2|}\log_2\left(\frac{|G_2|-QF_m+QF_{m,G_1}}{|G_2|}\right)\right). \tag{13}$$

Finally, by summing (12) and (13), we derive the Entropy Sort Equation, which calculates the conditional entropy for any RF/QF pair:

$$CE = H_{G_1} + H_{G_2}. \tag{14}$$

By defining the ESE to have only one variable, we have formed a continuous, bounded function. The curve produced by this equation represents a spectrum of conditional entropies that are defined by all possible arrangements of the QF with respect to the RF. In the main text we demonstrated that the resulting parabolic curve has a common structure for any RF/QF pair, with distinct properties that can give us new insights into the relationships between features.

One of the properties of the ESE that makes its use computationally tractable is that it is differentiable, such that;

$$\frac{dCE}{dx} = \frac{log_2(\frac{QF_m - x}{G_2}) - log_2(\frac{x}{G_1}) - log_2(\frac{G_2 - QF_m + x}{G_2}) + log_2(1 - \frac{x}{G_1})}{G_1 + G_2}. \tag{15}$$

Setting the left hand side of Eqn.(15) equal to zero and rearranging the equation we find,

$$x = \frac{|G_1| * QF_m}{|G_1| + |G_2|}. \tag{16}$$

By calculating the second derivative, we prove that this point is a maximum.

$$\frac{d^2CE}{dx^2} = -\frac{1}{G_1 + G_2}\left(\frac{1}{x} + \frac{1}{G_1 - x} + \frac{1}{QF_m - x} + \frac{1}{G_2 + QF_m + x}\right). \tag{17}$$

Since $x \geq 0$ (we cannot have less than 0 minority states overlapping), and $x \leq G_1$, $x \leq G_2$, $x \leq QF_m$, it follows that $\frac{d^2CE}{dx^2} < 0$, and the turning point is a maximum. Therefore, using Eqn. (16) we can calculate the maximum $CE$, from the value of $x$ where the RF and QF are independent.

#### SI 3 Visualising ES hypothesis testing and error potential

In Fig 2B we introduced $DPC_{Dependent}$ (green line) and $DPC_{Independent}$ (orange line). The parabolas in Fig SI 3 allow us to explain why $DPC_{Dependent}$ is calculated such that the green line in Fig 2B intersects with the global minimum, rather than the point (9,0) as the orange $DPC_{Independent}$ line does.

Fig SI 3A and B correspond to RF/QF arrangements in which SD = -1, whereas Fig SI 3C and D show scenarios where SD = 1. On each plot the black brackets enclose a region of potential hidden error/imperfect dependence penalty. Such a penalty refers to the fact that for each of the examples, even if the observed CE was found to exist at the local minimum or global minimum, there would be a portion of $CE$ that can never be removed from the system. As such, the assumption of dependence between the RF and QF can never be completely realised. This demonstrates that the ESE correctly identifies the sub-optimal dependency by assigning a local minimum with $CE > 0$. The phrase 'region of potential hidden error' describes the region of the data in which the addition of erroneous data points cannot lead to observable divergence.

To demonstrate the importance of the region of hidden error, in Fig SI 3A, two erroneous sample states were introduced to the ground truth (marked with an X), each of which overlapped with minority states ($x = 2$) and hence produced observable divergence on the ESE parabola (green line). Conversely in Fig SI 3B, 6 erroneous sample states are introduced, but 4 of these errors occurred in the region of potential hidden error. As such, the observable divergent points is 2 rather than 6 ($x = 2$). The region of potential hidden error means that the $DPC_{Dependent}$ calculation (Eqn. 6) automatically penalises $DPC_{Dependent}$ when there is a region of ambiguity regarding the dependent relationship between the RF/QF. Accounting for this region of ambiguity is one of the properties of ES that allows hypothesis testing and EP (Eqn. 8) to quantify the likelihood of dependency between features been disrupted by erroneous data points vs. the two features being independent.

To visualise ES hypothesis testing further, on each plot in Fig SI 3 we transposed the orange line representing $DPC_{Independent}$ along the y-axis to intersect with the local/global minimum (red dashed line). From this red dashed line we can find the point at which $DPC_{Dependent}$ (gradient of the green line) is equal to $DPC_{Independent}$ (gradient of the orange line), which we highlight as a red dot where EP = 0 (Fig SI 3A, C). This is the tipping point for the RF/QF system, where divergent samples are equally likely to be present due to error or feature independence. Hence we can clearly see that there exists a finite window in Fig SI 3A and C where positive EP values may be calculated ($0 < x < 6$ in Fig 2A and $7 < x < 12$ in Fig SI 3C).

Finally, observe that in Fig SI 3B and D, the red dashed line only intercepts the ESE parabola at the local/global minimum. This indicates that because the RF/QF states are so heterogeneous, we can never obtain evidence that divergent data points are present due to the introduction of error, since $DPC_{Independent}$ is always larger than $DPC_{Dependent}$.

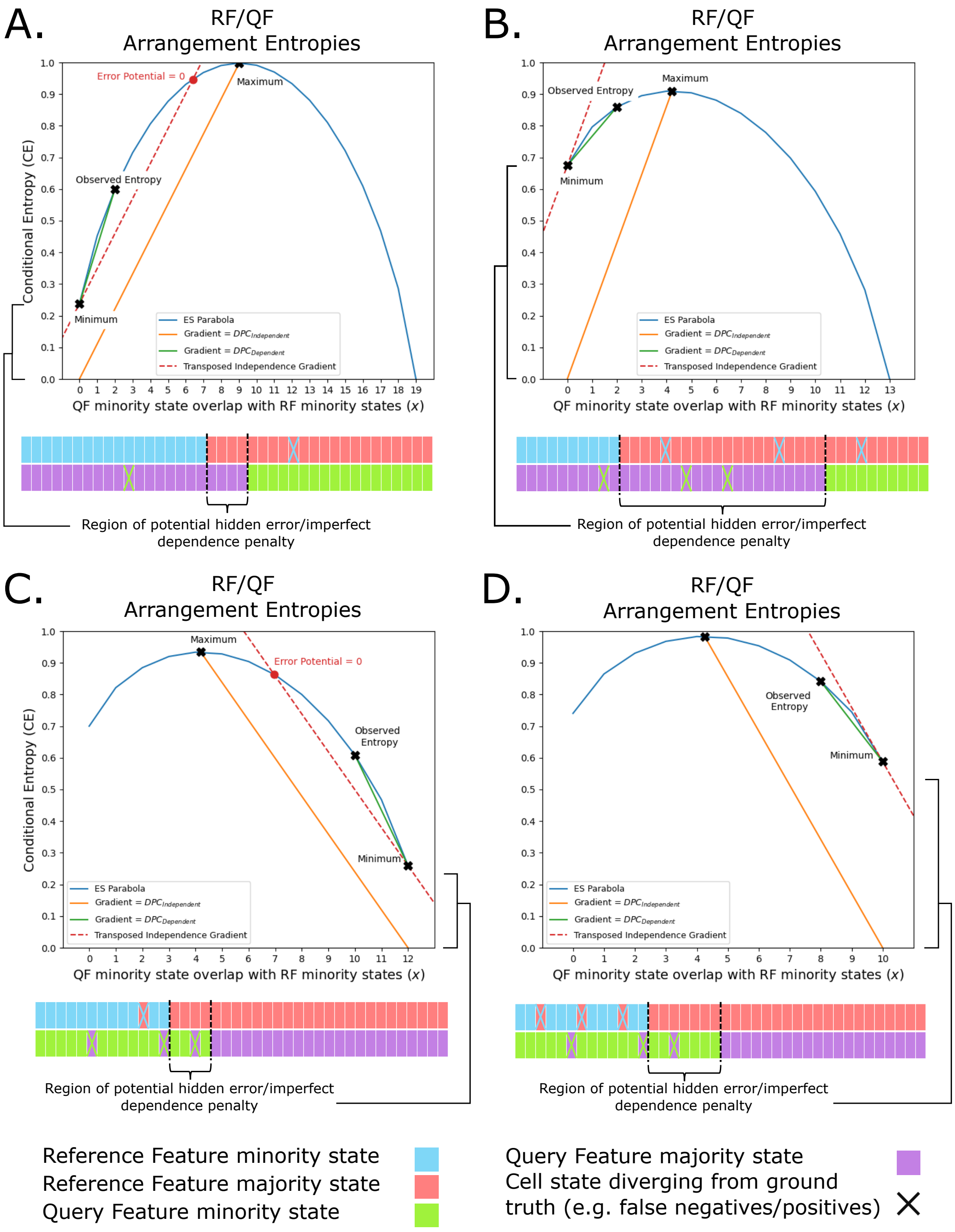


**Figure SI 3. Visualising ES hypothesis testing and error potential.** A and B present scenarios where SD = -1, and C and D where SD = 1. Black square brackets highlight the RF/QF imperfect dependence penalty that causes the ESE parabola minimum to have $CE > 0$. In B and D, the imperfect dependence penalty has become large enough such that the error induced uncertainty per cell (green line gradient) cannot become greater than or equal to the independent uncertainty per cell (orange line). As such, the red dashed line only intersects with the ESE parabola at the minimum.

#### SI 4 Identifying the Reference Feature

In this section, we provide the rationale for determining which feature from a pair should be considered the RF or QF. In agreement with the established knowledge regarding conditional entropy, the ESE is non-symmetrical. In the notation of conditional entropy this corresponds formally to

$$H(RF_m|QF_m) \neq H(QF_m|RF_m). \quad (18)$$

We can view the asymmetry graphically if we take two example features and plot the ES parabolas generated when each feature is assigned as the RF (Fig. SI 4A, B). As expected, if the minority states of each feature have different cardinality, the ESE parabolas do not overlay with each other. However, they do have key common properties, such as existing within the same domain and having identical shapes. In fact, the two curves are simply linear transformations of one another along the y-axis. The curves exist within the same domain since the number of possible overlapping states between both features is encapsulated by the feature with the smaller minority state group. The magnitude of the transformation along the y-axis between the two curves can be identified by inspecting the maximum of each curve. Since the maximum is equivalent to each feature being independent from one another, the distance between the two curves is the difference in entropy:

$$\text{y-axis transformation} = |H(Feature\ 1) - H(Feature\ 2)|. \quad (19)$$

For any ES parabola, the QF is that which describes the possible rearrangements of the system and hence the conditional entropy along the ESE parabola. From this we can infer the important property that for any pair of features, the curve with the maximum entropy will be described when the feature with the larger minority state group is the QF.

Having identified the relationship between the two ES parabolas generated from a pair of features, we now consider the consequence of allowing either feature to act as the RF. Quantifying the correlation between two features will produce different values based on which feature is the RF, leading to potential asymmetry in the results. For example, if we were to take a set of features and create a matrix of feature correlations via Pearson's Correlation, the resulting matrix would be symmetric. This has practical consequences such as allowing the matrix to be used as a distance matrix for downstream analysis. Conversely, if we were to fix the RF for each row of the correlation matrix when calculating the ESS (Eqn. (4)), the upper and low triangles of the resulting matrix would not be equal. This would arise because the SW (Sort Weight) terms of the ESS would be different for a pair of features depending on which was the RF. This is because the weight term is dependent on the maximum of the ESE parabola, whereas SD and SG are not.

From the perspective of ES hypothesis testing, ambiguity around which feature is the RF further complicates the analysis. A critical example is when calculating $EP$ for a pair of features. Since the ESE parabolas have identical shapes regardless of which feature is the RF, $DPC_{Dependent}$ (Eqn. 6) is equal for both ESE parabolas and so the assignment of the RF has no effect. However, $DPC_{Independent}$ (Eqn. 7) changes depending on which feature is assigned as the QF, since the ESE parabola maximum moves along the y-axis. Hence, if we do not identify a rationale to determine which feature is the RF in any pair, we cannot quantify errors across an entire data set without ambiguity.

To address the problem of RF assignment we present two distinct arguments. The first considers the Principle of Maximum Entropy (Jaynes 1957a; Jaynes 1957b), while the second is our own rationale that allows us to reject the ES parabola with the lower maximum entropy.

##### *Maximum entropy principle*

The Principle of Maximum Entropy (MaxEnt) (Jaynes 1957a; Jaynes 1957b), states that given data and some constraints on a system, the probability distribution with the maximal entropy that satisfies those constraints should be chosen to represent the underlying data. We can relate MaxEnt to the problem of identifying the best distribution that describes an ES system. When setting up an ESE parabola for a pair of features, we have two distributions to choose from. Each distribution is constrained to the same domain and their maxima occur at the same value of $x$. MaxEnt states that we should always pick the distribution with the higher maximum entropy, which we have already identified as where the feature with the larger cardinality of minority states is designated the QF. Typically, identifying the distribution with maximum entropy would have to be approximated using mathematical approaches such as Lagrange Multipliers, as in more general systems there could be an infinite number of distributions that would satisfy the given constraints. However, we are fortunate that for our purposes there are only two possible distributions.

We can further justify the application of MaxEnt through Lesne's work on the Entropy Concentration Theorem (Lesne 2014). The Entropy Concentration Theorem rigorously quantifies the observation that when the number of samples in a data set is sufficiently large, the number of microscopic states (i.e. RF/QF pairs) underlying the MaxEnt distribution is exponentially larger than under any other distribution. The implication of this is that for a given data set with defined constraints, it is exponentially less likely that the observed values were sampled from any distribution other than the MaxEnt one.

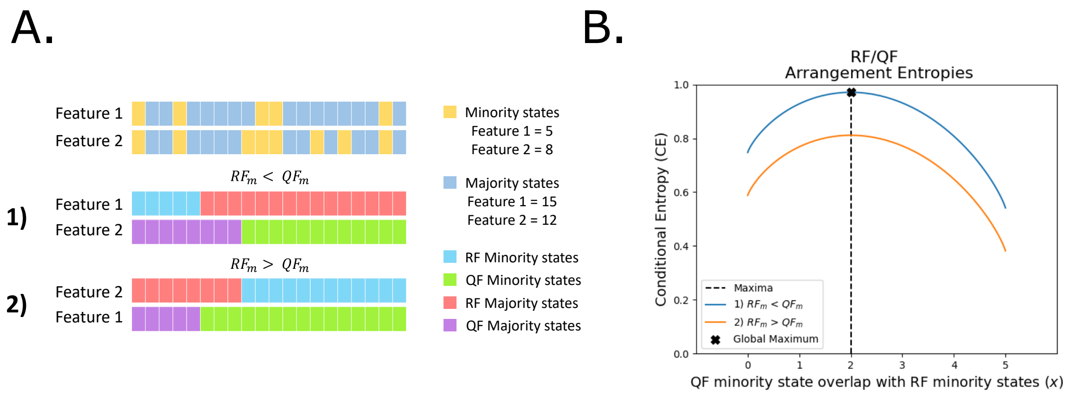


C.

#### RF/QF Microscopic State Summary

| | Microstate Cardinality | Minority State Overlap Frequencies | | | | | | |
|---|---|---|---|---|---|---|---|---|
| | | $x = 0$ | $x = 1$ | $x = 2$ | $x = 3$ | $x = 4$ | $x = 5$ | Sum |
| **1)** $RF_m < QF_m$ | $\binom{20}{8} = 125970$ | 6435 | 32175 | 50050 | 30030 | 6825 | 455 | 125970 |
| **2)** $RF_m > QF_m$ | $\binom{20}{5} = 15504$ | 792 | 3960 | 6160 | 3696 | 840 | 56 | 15504 |
| | | | | | | | | Sum |
| Overlap Probabilities: | | 0.051 | 0.255 | 0.397 | 0.238 | 0.054 | 0.004 | 1.0 |

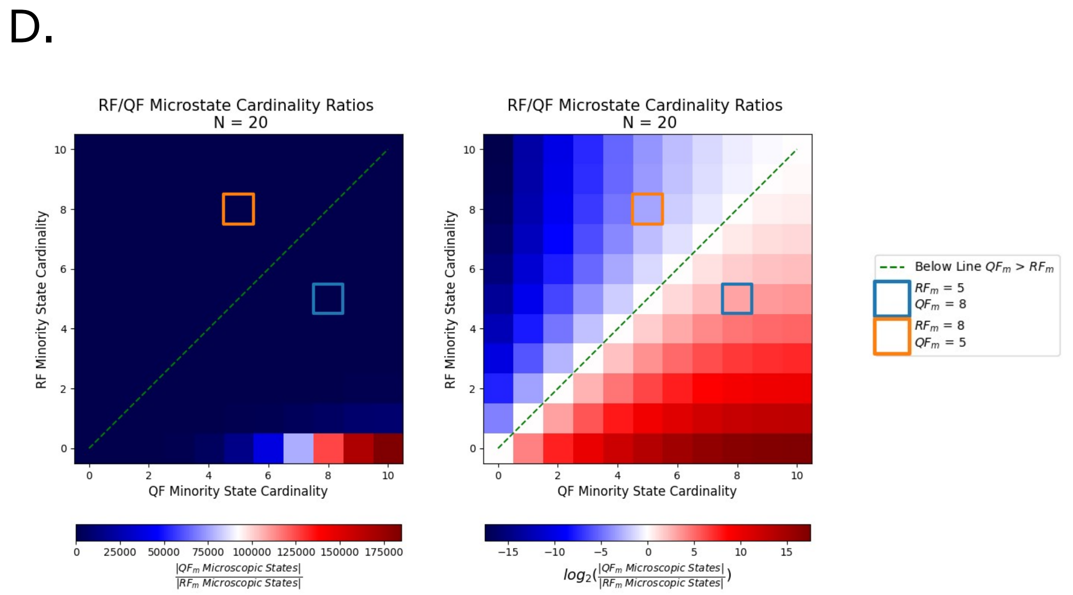


**Figure SI 4.** **The maximum entropy principle informs which feature should be the RF. A.** A simple example in which we could consider the feature with fewer minority states as the RF (1) or the feature with more minority states as the RF (2). **B.** The ESE parabolas formed from (1) or (2) from A. **C.** Summary of the frequency distributions for the observed RF/QF minority state overlaps ($x$) that could occur from all possible arrangements of the QF for (1) and (2). **D.** Heatmaps showing the ratio of all possible QF minority states vs. all possible RF minority states changes exponentially as the difference in cardinality of the RF/QF minority states increases linearly.

We can demonstrate this with regards to ES via a simple example (Fig. SI 4A). Let the number of samples ($N = 20$) be small enough such that we can apply brute force to identify every microscopic state. We then observe two features, $F1$ and $F2$. $F1$ contains 5 minority states ($F1_m = 5$) and $F2$ contains 8 minority states ($F2_m = 8$). As expected, if we plot the two possible ES parabolas, the parabola with the global maxima occurs when $F2$ is the QF (Fig. SI 4B). ES requires us to fix the arrangement of samples based on the RF and then consider all permutations of the QF, observing the frequency of overlaps between the $RF_m$ and $QF_m$ states. The number of microscopic states (M) for any RF/QF pair is easily obtained as the number of possible arrangements of the QF minority states. Hence,

$$M = \binom{N}{QF_m}. \tag{20}$$

We can reformulate Eqn 20 in terms of $M_x$, the number of microscopic states corresponding to each possible observable overlap of minority states, $x$, within the domain of the ESE such that,

$$M = \sum_{x=0}^{RF_m} M_x = \sum_{x=0}^{RF_m} \binom{RF_m}{x} \binom{N - RF_m}{QF_m - x}. \tag{21}$$

We summarise the results of Eqn. 21 for our example in Fig. SI 4C. First observe that the total number of possible microscopic states when the $RF_m < QF_m$ is much larger than when the $RF_m > QF_m$ (125970 vs. 15504). This is in agreement with Lesne's Entropy Concentration Theorem (Lesne 2014), stating that the probability of observing a particular value of minority state overlap through sampling the MaxEnt distribution is exponentially more likely than from the other distribution. Hence, we should select the MaxEnt distribution as the best description of the system.

To further confirm that the two possible distributions are constrained in the same manner, if we count the frequencies of each value of $x$ (Fig. SI 4C) amongst all possible microscopic states, we find them to be in equal proportions for both distributions. As such, the probability of each observation of $x$ is equal, regardless of which distribution you choose, demonstrating equivalency. Finally in Fig. SI 4D, we illustrate how quickly the exponential nature of the Entropy Concentration Theorem is realised by presenting the ratio of RF/QF microscopic state cardinalities when we vary the value of $RF_m$ and $QF_m$. Notice that even at a small value of $N$ ($N = 20$), the number of possible microscopic states when $QF_m > RF_m$ is orders of magnitude higher than when $QF_m < RF_m$. This exponential increase is even more dramatically amplified as $N$ increases, making it a strong rationale for selecting the MaxEnt ES parabola over the lower entropy parabola.

##### *Consistent Vs. Inconsistent sampling distributions*

Here we present the rationale for excluding scenarios where $RF_m > QF_m$. We consider a RF/QF system and demonstrate that, when $RF_m < QF_m$ (the MaxEnt distribution), sampling from the system while viewing the two features as independent is equivalent to viewing the two features through the ESE under the assumption of independence (the maximum of the ESE parabola). Conversely, when $RF_m > QF_m$, we show that the frequency distribution when treating the two features as independent is different from the frequency distribution generated when viewing the same system through the ESE but constrained to the assumption of independence. As such we find that all scenarios where $RF_m > QF_m$ must be sampling from different distributions under the assumption of independence compared to viewing the system through the ESE, and hence they are incompatible and cannot be used for ESE hypothesis testing. This rationale is similar to Jaynes's Principle of Maximum Entropy (Jaynes 1957a; Jaynes 1957b) and Lesne's work on the Entropy Concentration Theorem (Lesne 2014), in that we are constraining the system to an assumption – feature independence – and identifying that only the MaxEnt distribution produces equivalent minority state overlap ($x$) frequency distributions between independent features and the ESE under this same constraint.

We consider an example in which $N = 50$, $F1_m = 10$, and $F2_m = 15$ (Fig. SI 5A). In Fig. SI 5B we plot the ESE parabola (blue curve) for the scenario where $RF_m < QF_m$, such that $F1$ is the RF and $F2$ the QF. The domain of the ESE is constrained between 0 and the value of the feature whose minority state has the smaller cardinality, which here is $F1_m = 10$. The ESE parabola describes the $CE$ if we were to partition the data by the states of $F1$ and then inspect different arrangements of the $F2$ (QF) minority states. For any observed arrangement we could assume that the observed overlap between the minority states, $x$, has occurred due to two distinct sampling regimes. The first would be that the observed RF and QF data sets have been sampled from two independent distributions and the results are then compared against one another to identify how many minority states overlap. Over many re-samplings of $N = 50$ samples, we would be able to approximate the expected value of minority state overlap ($E(x)$), between the two independent features. The second way that we could view the sampling of the minority state overlaps would be to postulate that there may be a dependency between the two features, and that we should use the ESE to model the system.

As mentioned at the start of this section, regardless of whether we view the sampling as two independent features or two features with possible dependence, we can constrain both models to an assumption of independence. We will now identify

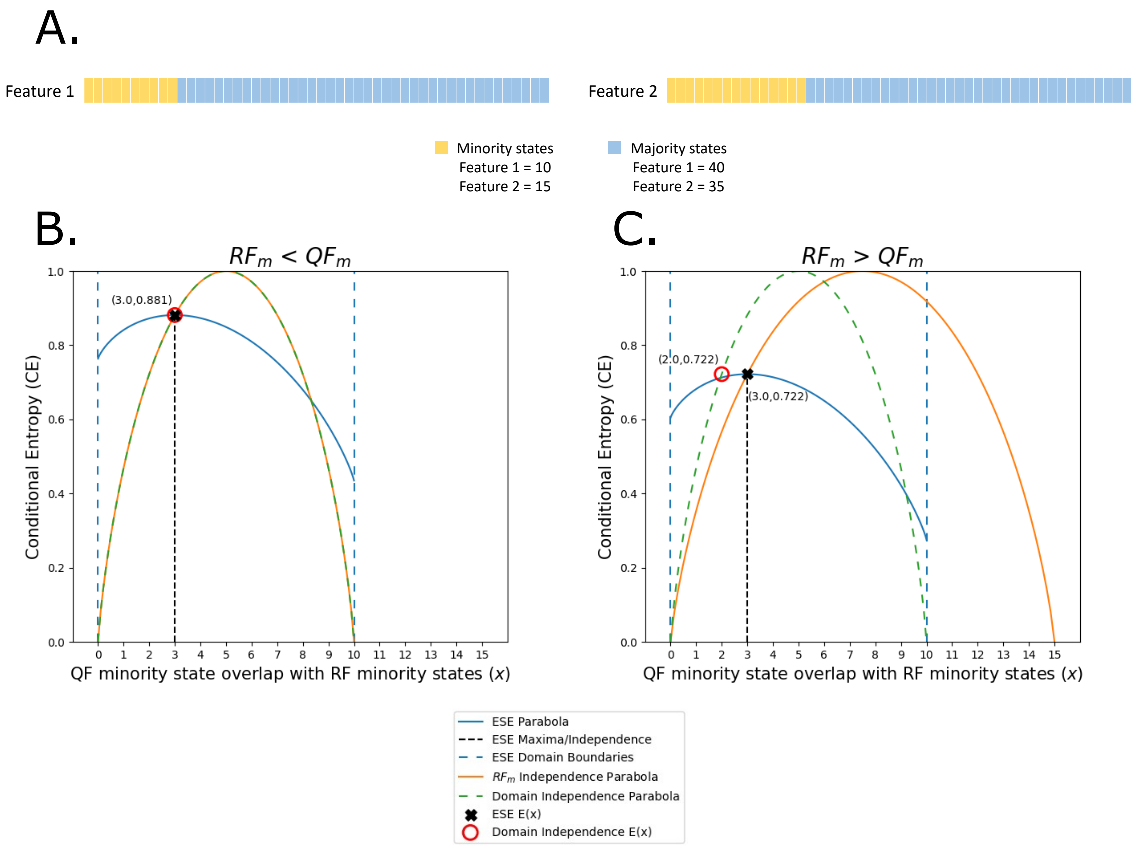


**Figure SI 5. Incompatible frequency distributions when $RF_m > QF_m$ allow us only to accept RF/QF scenarios where $RF_m < QF_m$. A.** A simple pair of features for exploring the independent and dependent frequency distributions of $x$ (the number of overlapping RF and QF minority states) when $RF_m < QF_m$ or $RF_m > QF_m$. **B, C.** The ESE and independence parabolas that arise when considering the observed RF/QF systems as having been sampled from either a dependent system between the two features through the ESE, or as a completely independent system where the RF/QF samples are drawn from separate distributions.

the value of $E(x)$ under the assumption of sampling from two independent distributions, or using the ESE constrained to the assumption of independence.

To identify $E(x)$ under the assumption that $F1$ and $F2$ are taken from two independent distributions, we first note that the probabilities for observing a minority state when sampling from independent RF or QF distributions can be found by calculating the fraction of RF or QF states observed as minority states in all 50 samples. Hence, $P(F1_m) = \frac{10}{50} = 0.2$ and $P(F2_m) = \frac{15}{50} = 0.3$. Now, since $F1$ is the RF when $RF_m < QF_m$, when we partition all samples based on the RF states, the size of group 1 is 10. We use this knowledge to subset the QF sampling of minority states down to the same domain of the ESE so that they are comparable. This is justifiable since if the RF and QF are truly independent, then reducing the number of samples to $N = 10$ has no effect on the properties of the distribution the QF states are sampled from. We are simply saying that we found 10 samples displaying the minority state of the RF, and if we were to retrieve 10 samples from the QF distribution, how many minority states would we expect to find (how many $QF_m$ would overlap with the $RF_m$)? Since $P(F2_m) = P(QF_m)$, then $E(x) = 0.3 \times 10 = 3$. In Fig. SI 5B we show this expected value under independent sampling as the red circle. In summary, we have shown that if the RF and QF are sampled from two truly independent distributions we would expect to find on average 3 overlapping minority states.

Now we check whether using the ESE to identify $E(x)$ under the assumption of independence produces the same value of 3. To do so we identify $x$ at the maximum of the ESE parabola – the point at which the overlap of minority states between the two features is completely independent. From this we find that once again $E(x) = 3$ (black 'X', Fig. SI 5B). This demonstrates that when $RF_m < QF_m$, considering the two features as sampled from separate independent distributions or quantifying their relationship via the ESE appear to be sampling from equivalent distributions when constraining the system to an assumption of independence.

We now demonstrate that performing the same analysis when $RF_m > QF_m$ leads to incompatible $x$ frequency distributions.

In this scenario, $E(x) = 3$, since this remains the value of $x$ at which the maximum of the ESE parabola occurs (black 'X' in Fig. SI 5C). If we calculate $E(x)$ for the completely independent RF/QF sampling distributions as for Fig. SI 5B, we find that $E(x) = P(QF_m) \times 15 = 0.2 \times 15 = 3$. Hence, up to this point $x$ appears to be sampled from the same distribution, regardless of whether you view the system as two independent sampling distributions or two dependent features through the ESE. In Fig. SI 5C this is apparent as the point of intersection between the ESE parabola (blue) and the independent distributions parabola (orange) is at the maximum of the ESE parabola, where $E(x) = 3$. However, when we extrapolate the independence curve (orange curve), we find that the range of observable $x$ values for the scenario where the RF/QF are sampled from independent distributions is incompatible with the frequency distribution when observing the system through the ESE. In other words, even though the frequency distribution of $x$ have the same $E(x)$ under each alternative scenario, they can take on a different range of values, and hence cannot be being drawn from the same underlying distribution. For example, $0 \leq x \leq 10$ through the ESE, whereas $0 \leq x \leq 15$ through the independent sampling regime. This occurs because, albeit unlikely, sampling $N = 15$ samples from the independent $F1$ distribution could theoretically lead to drawing a value of $x > 10$ minority states. Conversely, when viewing the same pair of independent features through the ESE, it is impossible to observe $x > 10$, because the domain of the ESE is constrained to that of the feature with the smallest minority state cardinality.

To further demonstrate that independent sampling of the RF/QF compared to through the ESE are incompatible when $RF_m > QF_m$, we force the independent sampling regime to have the same range of observable $x$ values as the ESE. We can do so by only sampling $N = 10$ samples for the QF ($F1$). Having altered the sampling range, we can re-calculate $E(x)$ such that $E(x) = P(QF_m) \times 10 = 0.2 \times 10 = 2$. Because $E(x) = 2$ (red circle, Fig. SI 5C) for the independent sampling when constrained to the domain of the ESE, compared to $E(x) = 3$ when observing the system through the ESE, we again find that the underlying sampling distributions are different. As such, when $RF_m > QF_m$, the hypothesis that the observed data could arise from sampling either two independent features or two dependent features described by the ESE are incompatible, and we must reject all such possibilities. As such, we can always accept the RF/QF arrangement where $RF_m < QF_m$.

In summary, we have provided two distinct rationales for designating the feature with the larger minority state cardinality as the QF for any RF/QF pair. Our first justification demonstrates how the Principle of Maximum Entropy may be applied to the ES framework to preferentially choose the ESE parabola with the maximum entropy over the alternative ESE parabola. The second justification demonstrates that if we were to choose the ESE parabola with the lower entropy, the implied sampling regimes under the assumption of independence and possible dependence through the ESE would not be equivalent when constrained to independent sampling. As such, we must reject the scenario with lower maximum entropy. Being able to demonstrate that not using the MaxEnt distribution in our hypothesis testing would lead to testing results from different distributions removes ambiguity around which feature should be the RF and which should be the QF. The practical need to reject the lower entropy parabola becomes apparent in the context of ES hypothesis testing and algorithms, FFAVES and ESFW.

#### SI 5 FFAVES

In Fig. SI 6 we present pseudocode for the FFAVES algorithm. FFAVES formulates theory from ES into a workflow that takes discrete high dimensional data and iteratively switches the expression states of those data points that appear to be in the wrong state until the system converges. There are multiple ways in which this task could be formulated and hence FFAVES is just one potential workflow for implementing ES. Additionally, recall that we derived the ESE such that $x$ designates how many RF minority states overlap with QF minority states (SI 2). Hence, in this arrangement of the ESE a FP always designates when a data point displaying a minority state of a given feature should instead display the majority state. Conversely, a FN indicates that a sample displaying the majority state of a feature should instead display the minority state.

A summary of the nomenclature of the FFAVES pseudocode is provided alongside Fig. SI 6, however it is useful to clarify a few symbols.

- $m_{min}$: The minimum cardinality for an accepted feature, i.e. the minimum number of minority states a feature can have before it is excluded from analysis.

  The default value of $m_{min} = 10$, such that features with fewer than 10 data points in the minority state will be ignored when running FFAVES. The default value is a relatively arbitrary cutoff, but we provide an empirical motivation for selecting a default of $m_{min} = 10$ in Section. SI 7. There we demonstrate that for $m_{min} < 20$, the sensitivity of a feature to FN data points reaches a point where ES hypothesis testing automatically detects that there is not enough information to suggest FNs through positive $EP$ values (Eqn. 8). Hence, ES hypothesis testing incorporates quality control checks that will tend to be automatically enforced at a value of $m_{min} > 10$, further demonstrating the arbitrary nature of the $m_{min}$ parameter.

- $CI$: The confidence interval. A threshold used to determine whether the $EP$ values of individual data points are statistically unlikely to be part of a half normal distribution of all calculated $EP$ scores.

  The default value of $CI = 0.99$, such that $EP$ values found outside this confidence interval have only a 1% chance of being a member of the distribution. As such, we can confidently say that these data points are anomalous and suggest them to be FP or FN expression states.

- $T$: The tolerance. A threshold for when the expression states of the FFAVES adjusted matrix ($M'$) have converged, and FFAVES may terminate before reaching the maximum number of iterations defined by Max_Cycles.

  The default value of $T = 0.1\%$, such that if the change in the number of suggested FN data points between cycle $i$ and cycle $i-1$ is less than 0.1% of all data points in $M'$ for three cycles in a row, FFAVES will terminate early.

The pseudocode in Fig. SI 6 shows that FFAVES has three main steps.

1. Identify statistically significant FP data points and switch their states in the discretised matrix ($M$).

2. Identify FN data points in the adjusted state matrix ($M'$), followed by a second state switch of data points that appear in the wrong state due to being FNs.

3. Repeat step 1 on $M'$ to identify further possible FP data points. If any of the suggested FPs identified in step 3 were suggested as FNs in step 2, they are considered spurious suggestions and removed from the set of suggested FN points.

The final set of FN data points are saved and applied to the data at the start of the next iteration. This continues until the algorithm reaches the maximum number of cycles or converges within the tolerance (T) limit.

The workflow outlined in Fig. SI 6 is an example of a specific application of ES. It is intentionally conservative in suggesting that the minority state of any feature be enriched. The motivation for this is that we are often primarily interested in how the presence of uncommon minority state observations distinguish samples from one another. In other words, real minority state expression states are information rich. Correctly identifying FN minority state expression states through ES amplifies the prevailing structure in the data. However, the unintentional introduction of FP minority states diminishes the value of the real minority states, potentially leading to a lower resolution of differing sample identities than if we had not tried to correct the expression states at all (Andrews and Hemberg 2019). For example, if shared gene expression states between distinct cell types through FPs, those cell types may end up looking more similar in the data than they really are.

To minimise the introduction of FP minority states, Step 1 aims to prune minority state expression states that appear significantly contradictory to the prevailing overlapping state structure in the data. This is important for Step 2, when FFAVES seeks to identify FN minority state data points. If the spurious minority state points from Step 1 are not pruned, they could provide evidence for suggesting FN data points with equal weighting as data points that have low potential of being FPs. Hence

**Algorithm 1:** FFAVES

**Input** : M, $m_{min}$, CI, T, Max_Cycles
**Output** : Suggested Divergent Points
**begin**
  i = 0
  **while** $i \leq Max\_Cycles$ & $\Delta FN < T$ **do**
    *M' = M # Always start from initial discretised matrix*
    **if** *i > 0* **then**
      *# Switch FN states learned from previous cycle*
      $M'[FN_{i-1}] = (M'[FN_{i-1}] \times -1) + 1$
    **end**
    ***Step 1: Identify False Positive (FP) data points***
      *1) Calculate FP error potential matrix*
      *2) Find statistically significant FP divergent indices*
      *3) Temporarily switch states of FP data points*
        $M'[FP] = (M'[FP] \times -1) + 1$
    ***Step 2: Identify False Negative (FN) data points***
      *1) Calculate FN error potential matrix*
      *2) Find statistically significant FN divergent indices*
      *3) Switch states of FN data points*
        $M'[FN] = (M'[FN] \times -1) + 1$
    ***Step 3: Identify spurious suggested False Negative (FN) data points***
      *1) Calculate FP error potential matrix*
      *2) Find statistically significant FP divergent indices*
      *3) Null FN data points that also appear as FPs*
    ***Save suggested FP and FN indices***
  **end**
**end**

**Figure SI 6. Pseudocode of the FFAVES algorithm.** $M$ = input discrete state matrix, $M'$ = discrete matrix augmented by suggested divergent points, $m_{min}$ = minimum minority state cardinality for an accepted feature (default = 10), $CI$ = confidence interval for identifying divergent points (default = 0.99), $T$ = tolerance for convergence (default = 0.1%), Max_Cycles = maximum number of cycles before terminating FFAVES (default = 15), $i$ = cycle number, FN = suggested false negative data points, FP = suggested false positive data points, $\Delta FN$ = change in number of suggest FN data points between cycle $i$ and cycle $i-1$ as a percentage of the number of data points in $M$.

by temporarily removing likely FP data points in Step 1, we mitigate the possibility of poor quality data points generating more poor quality data points.

An example of this could be cells transitioning from one state to another. As cell identity evolves from cell type 1 ($CT_1$) to cell type 2 ($CT_2$), sets of genes are upregulated and downregulated. However, there may be a delay in the breakdown of some RNA species and we may detect expression of a gene specific to $CT_1$ in a sample belonging to $CT_2$. Failure to implement Step 1 in FFAVES could lead to ES suggesting that the sample from $CT_2$ should inherit a portion of the gene signature from $CT_1$. This is because FFAVES would quantify that it is rare to find the spurious $CT_1$ gene active in the absence of other $CT_1$ genes. By implementing Step 1 we minimise the likelihood of these pre-existing FPs events generating new FPs.

Step 3 is another quality control step designed to minimise the introduction of FPs. Once again consider that there was enough evidence in Step 2 to suggest that a gene specific to $CT_1$ should be expressed in $CT_2$, but it is being displayed as inactive (i.e. a FN). This may be due to stochastic overlapping gene signatures as the cell transitions between the two cell states. Now imagine that in Step 3 that same gene expression point is also found to be a FP due to the conflicting gene signatures. In other words, the data point is found to be both a FN and a FP. In this scenario, Step 3 identifies that there is conflicting information. Hence, we cannot make a judgement as to whether the data point should change state or not, so we remove the data point from the list of suggested FNs, thereby defaulting back to the expression state observed in the data.

Note that state switching of potential FPs in Step 1 is only a temporary removal of suggested FP states from the initial matrix ($M$). We choose only to switch the states of proposed FN points at the start of each cycle for two reasons: i) We believe that the presence of FP minority states in $M$ is very low. This is primarily inspired by scRNA-seq data, where it is unlikely that

FP data is introduced during the experimental generation of the gene expression matrix. We believe this is not unreasonable for many data sets, but it is up to the user to decide. Thus, if they are not FPs, but real biological data points due to factors such a gradual decay of mRNA fragments, we do not want to entirely remove them from the data as they may elucidate gene expression gradients/dynamics in the whole data set. Rather, we temporarily remove them to protect from the introduction of FPs by FFAVES. ii) By returning to $M$ at the start of each cycle and only changing the states of suggested FN data points from the previous cycle, FFAVES checks whether FPs identified in the previous cycle are no longer significantly out of place in the adjusted data. If so, they can confidently be used to amplify the prevailing structure in the data to further elucidate FN data points. One way to interpret this is that we are anchoring the convergence of the expression states around the information rich, high confidence minority states.

##### *Calculating the error potential matrix*

The first step for identifying FP or FN states is to calculate the Error Potential Matrix (EPM). The EPM can exist in two forms: the false positive EPM ($EPM_{FP}$) and the false negative EPM ($EPM_{FN}$). These quantify the likelihood of data points in $M'$ being FP or FN respectively. Hence, $EPM_{FP}$ is used in Steps 1 and 3 of the FFAVES algorithm, and $EPM_{FN}$ is used in Step 2 (Fig SI 6).

We visualise the process of generating an EPM matrix in Fig. SI 7. The majority of ES theory is utilised in steps (1) and (2). Step (1) draws upon the logical deduction of 8 specific error scenarios where divergence could be observed under the assumption of dependence between two features (Fig. S2), and the application of the maximum entropy principle (Section SI 4), to determine which set of calculations should be undertaken to form the required EPM. Step (2) performs these calculations using ES hypothesis testing to quantify the evidence suggesting that data points in $M/M'$ are displaying the wrong state. The process of identifying the evidence that states are FPs or FNs is carried out for each feature. Hence, each column, $j$, of an EPM corresponds to the evidence suggesting that each sample, $i$, of feature $j$ in $M'$ is presenting as the wrong state.

We now describe each step of generating the EPM in more detail. Steps (1)-(4) in Fig. SI 7 highlight the process for a single feature, $j$, which we designate to be the Fixed Featured (FF). The FF refers to when we inspect a specific feature in the data and seek to quantify the evidence that the samples in the FF are FPs or FNs. During the process of quantifying the likelihood of FPs or FNs, the FF may act as either the RF or QF during ES hypothesis testing (determined in step (1)).

Step (1) determines how ES hypothesis testing should be carried out between the $FF$ and any other feature (Fig. SI 8). Having designated the $FF$ and $SF$, we first identify whether the $FF$ should be the RF or QF by comparing the cardinality of the $FF$ and $SF$ minority states ($FF_m$ and $SF_m$, respectively - see Section SI 4). Subsequently, we identify which of the 8 possible error scenarios (Fig. S2) any divergence observed in the $FF/SF$ pair relates to. We only consider scenarios in which $QF_m > RF_m$ (Section SI 4), and thus only ever observe error scenarios 2, 4, 6 and 8. Having applied the maximum entropy principle, we use the split direction (SD, Eqn. (1)) to determine the appropriate error scenario. If the $FF/SF$ pair corresponds to scenarios 2, 4 or 8, any observed positive $EP$ can be attributed to FPs in the $FF$. Conversely, if the $FF/SF$ pair corresponds to scenario 6, any observed positive $EP$ can be attributed to FNs in the $FF$ (Fig. S2). Having applied the logic in Fig. SI 8 to the $FF$ and all other features in the data set, we retrieve $j_{FP}$ and $j_{FN}$, which identify all features where the calculated $EP$ indicates the likelihood of FPs or FNs in the $FF$, respectively. We then feed $j_{FP}$ or $j_{FN}$ into step (2) of the EPM calculation (Fig. SI 7).

Step (2) performs ES hypothesis testing between the $FF$ and every relevant $SF$, as designated by $j_{FP}$ or $j_{FN}$. This produces a matrix with $i$ rows for each sample and $j_{FP}$ or $j_{FN}$ columns for each $SF$ (Fig. SI 7). Accordingly, each element of this matrix represents the $EP$ for $i$th sample in the $FF$ when it is considered against the $j_{FP}$th or $j_{FN}$th $SF$.

Step 3) ignores all negative $EP$ values by setting them to 0. ES tells us that negative $EP$ values indicate that divergent cells in the $FF/SF$ pair are more likely to have occurred due to feature independence than due to the introduction of error. Conversely, positive $EP$ values indicate that divergent samples are more likely due to the introduction of error. We are interested in quantifying all the evidence available that suggests that observed states in the $FF$ are erroneous.

Finally, in step (4) we sum the rows of the ES hypothesis test matrix, such that we are left with a vector of $i$ samples. Each element in this vector represents all evidence in the data set that suggests sample $i$ in the $FF$ is currently displaying the incorrect state in $M'$. This vector then fills column $j$ of the EPM.

Depending on whether the $EPM_{FP}$ or $EPM_{FN}$ matrix has been calculated, the next step of FFAVES is to identify if any of the states in $M'$ are statistically likely to be observed in the wrong state by identifying if their corresponding values in the EPM are unusually high. Such data points are considered to be consistently out of place based on the evidence provided by structural relationships between any $FF$ of interest and all the other features in the data.

##### *Identify statistically significant divergent data points*

Having generated an $EPM_{FP}/EPM_{FN}$, we now wish to identify states that are likely FPs/FNs. To explain our approach, we provide an example in Fig. SI 9, which relies on the synthetic data presented in this paper, which contains 5 synthetic cell types. These cell types are distinguishable from one another by modules of highly structured synthetic genes (Fig. SI 9A).

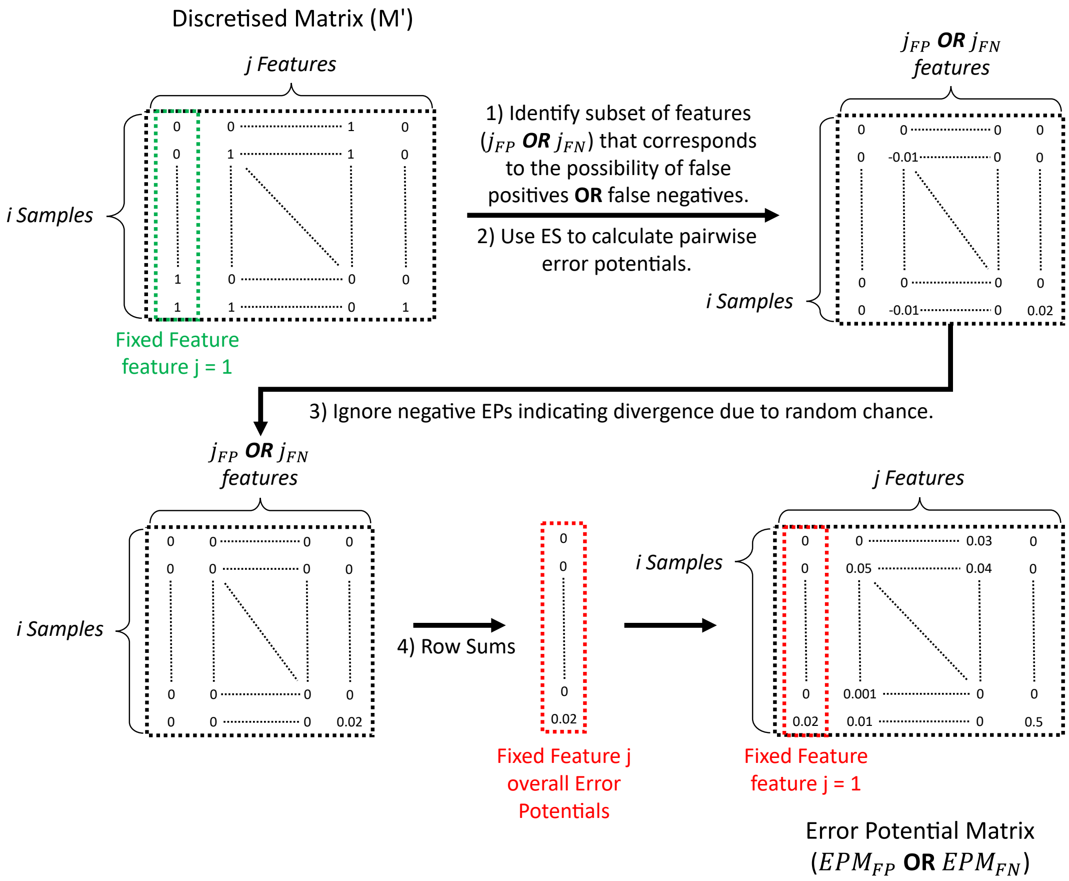


**Figure SI 7. Calculating an EPM.** The EPM is calculated by performing ES hypothesis testing in a pairwise fashion for every feature in the data. The EPM quantifies all the evidence available indicating that individual data points are in the wrong state in the discrete data matrix ($M'$). We summarise the calculation of an EPM in four main steps. **(1)** For each feature, identify all other features that would provide evidence that data points for the given feature are FPs/FNs. The identification of those features contained in $j_{FP}$ or $j_{FN}$ is summarised in Fig. SI 8. **(2)** Perform ES hypothesis testing to calculate all EPs (Eqn. 8) between feature $j$ and features $j_{FP}$ or $j_{FN}$. **(3)** Set all negative values to 0 (we are only interested in scenarios where there is evidence that data points are in the wrong state due to the introduction of error). **(4)** Calculate the row sums of the positive ES hypothesis test matrix to get the total error potential evidence that the samples of feature $j$ are displaying the wrong states in $M'$.

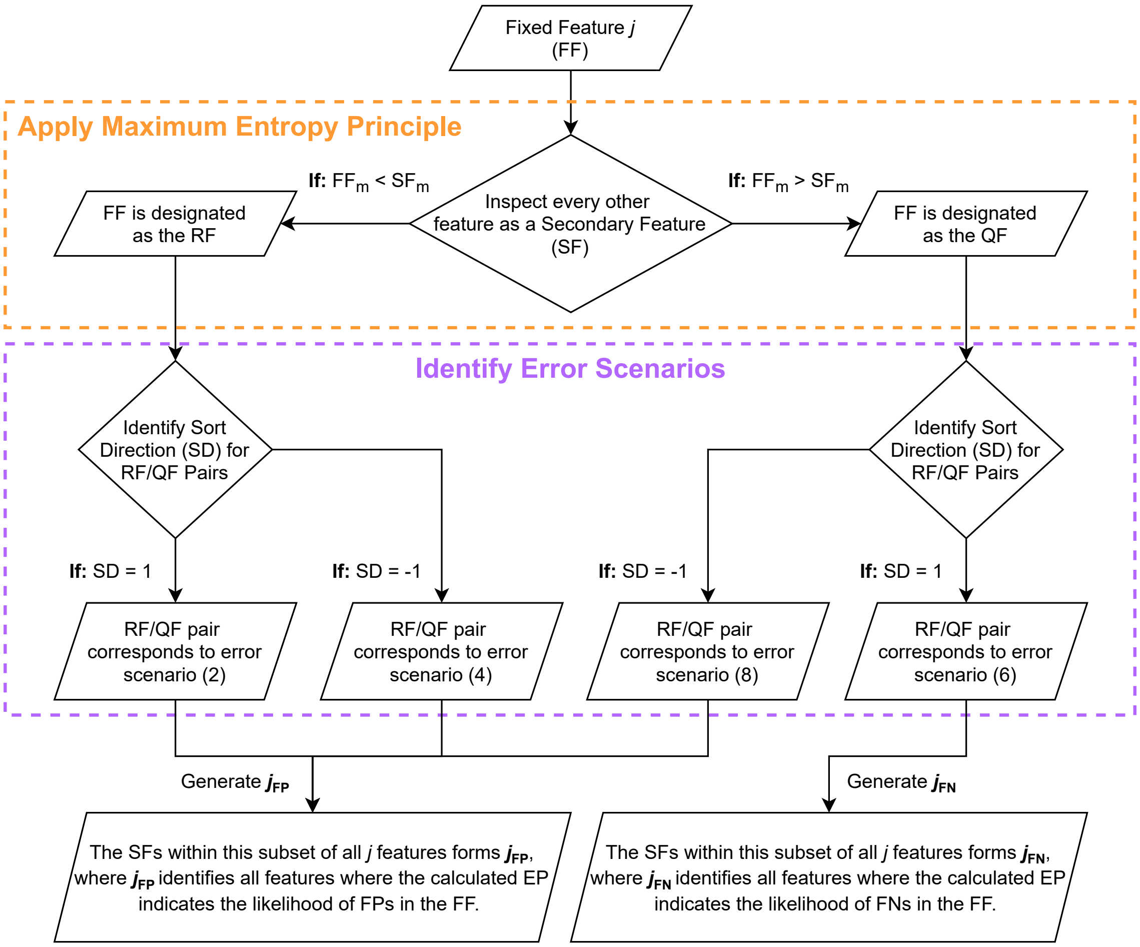


**Figure SI 8. Identifying $j_{FP}$ or $j_{FN}$ for any given feature $j$.** For any feature in a data set, there exists a subset of features that have the potential to inform whether expression states in $j$ are displaying as FPs ($j_{FP}$), and a subset of features that would suggest FNs ($j_{FN}$). Whether a secondary feature ($SF$) provides evidence for FPs or FNs is determined via whether the fixed feature ($FF$) has a larger or smaller minority state cardinality, followed by whether the $FF$ and $SF$ have split directions (SDs) equal to 1 or -1. Following this, for any $FF$ we can identify all the features that are members of $j_{FP}$ and/or $j_{FN}$.

In this example, we are specifically evaluating the $EPM_{FN}$ from in Step (2) of FFAVES (Fig. SI 6). We provide pseudocode for identifying statistically significant divergent data points in Fig. SI 10. Before attempting to find relationships or structure within the $EPM$, it is important to understand the two **if** statements at the start of the pseudocode. When calculating $EPM_{FP}$, we are only quantifying the EPs of all data points in $M'$ that display feature minority states. Conversely, when calculating $EPM_{FN}$ we are only quantifying the EPs of data points that are displaying majority states. Thus, when we seek to compare the values observed in an $EPM_{FP}$ or $EPM_{FN}$, we should only inspect the indices that are relevant to the possibility of FP or FN error, respectively. More precisely, when inspecting an $EPM_{FP}$ ($EPM_{FN}$), we should only take the indices in $M'$ that relate to feature minority (majority) states, since these are the only points where we can quantify the potential for FPs.

Having identified which data points within a given EPM are to be analysed, we can now examinine the distribution of EPM values to identify those that are statistically higher (Fig. SI 9C). From the histogram we observe that the vast majority of potential FNs have EPM values around 0. There is also an exponentially smaller frequency of values within a given bin of the histogram as the EPM values increase. This provides a firm basis to model the distribution of the data as a half-normal distribution. This should be generally true for the majority of high dimensional data sets. If we observe a highly structured data set with little to no FN/FP data points, then all of the highly structured data points should have little to no error and hence have EPM values close to or equal to 0. Conversely, if we have a data set with little to no correlative structure between feature states in $M$, then the only scenarios we should see positive $EP$ values during the calculation of the EPM would be by random chance. This in turn has a low chance of generating a high $EP$ score, and hence the vast majority of scores in the EPM will be close to or equal to 0. By fitting the data to a half-normal distribution, we are then able to use the resulting cumulative distribution function (CDF) to identify when values are significantly larger than the distribution mean. Before we can fit the data to a distribution it is useful to convert the data into z-scores. The z-score of any data point within a distribution is a measurement of how many standard deviations above or below the population mean the original data point is. Transforming into z-scores is useful as it allows us to easily identify which values lie outside of a given confidence interval when the data is fitted to a half-normal distribution. We calculate the z-scores as follows:

$$z - score = \frac{x - \mu}{\sigma}, \tag{22}$$

where $\mu$ and $\sigma$ are the mean and standard deviations of the data, respectively, and $x$ is the value taken from the EPM. Since we are treating the data as a half-normal distribution, $\mu = 0$.

We fit the data to a half-normal distribution using the *halfnorm* function from the *scipy.stats* python package. This takes the z-scores as input, and returns where each z-score lies on the CDF. This converts each z-score into a probability of whether sampling from the distribution would produce a value less than or equal to the given z-score. From this we can form confidence intervals to identify when individual data points are statistically unlikely to be members of the observed distribution. For the distribution in Fig. SI 9C, we identify that a z-score of 2.58 corresponds to a confidence interval (CI) of 0.99 (red dashed line). A CI of 0.99 indicates that there is a $1\%$ chance that if you were to sample from the fitted distribution you would observe a value greater than the respective z-score. Since it is statistically unlikely that observed z-scores outside this stringent CI (i.e. z-scores > 2.58) are part of the observed distribution, we denote any points with higher z-scores as statistically divergent data points. In this example we consider an $EPM_{FN}$ matrix, thus indicating that the statistically divergent points are FNs. The approach follows for $EPM_{FP}$ matrices and FPs. Returning to the context of ES, identifying FPs/FNs data points as having statistically high EPM values is equivalent to stating that there is a significant body of evidence within the pairwise relationships of the features to suggest that the currently observed state in $M'$ is incorrect.

In Fig. SI 9D we visualise the data points that are identified as having statistically high EPM values as determined above. We observe that the identified FNs are highly specific to the cell type specific regions of the data. This is an initial demonstration that ES and FFAVES can identify when feature states are likely incorrect based on the evidence provided by co-regulated features. Where FNs are identified outside of the highly structured cell types (cyan triangles) relates to samples created by combining 2 or more samples together (i.e doublets), where the true cell identity is ambiguous. Crucially, the presence of doublets in the synthetic data demonstrates that FFAVES does not borrow information from the doublet cells in a manner that deteriorates the identity of the singlet cells.

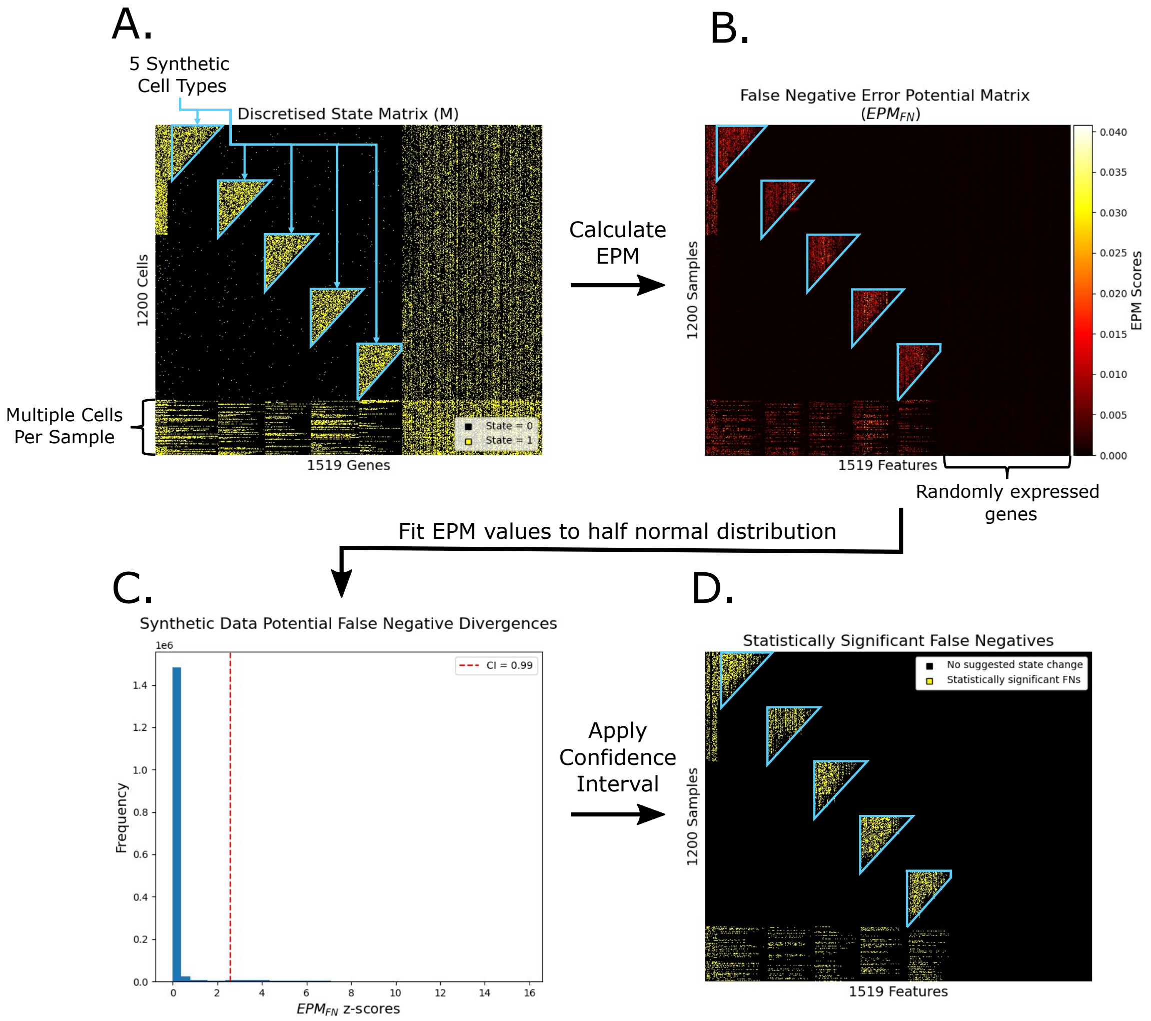


**Figure SI 9. Identifying statistically significant FP or FN data points from $EPM_{FP}$ or $EPM_{FN}$.** We show the steps taken to identify statistically significant FN data points in the first cycle of FFAVES, as applied to our synthetic data set.The process for identifying FPs is the same, but uses $EPM_{FP}$. **A.** The initial discrete state matrix, $M$. Each data point in state 0 is shown in black, and each in state 1 is shown in yellow. The triangular regions highlighted in cyan correspond to regions of highly structured gene expression that correspond to the 5 synthetic cell types. **B.** $EPM_{FN}$ - the heatmap is coloured according to the total $EPM_{FN}$ score calculated for each individual data point. Higher scores indicate more evidence that the data point is displaying the wrong state in $M$. **C.** To discriminate statistically high EPM values, we fit a half normal distribution, and identify all data points outside a 0.99 confidence interval. **D.** All points identified as having statistically significant evidence for being FNs in $M$ are highlighted in yellow.

**Algorithm 2:** Identifying statistically significant divergent data points.

**Input** : $M'$, $EPM$, $CI$
**Output** : Significantly Divergent Points
**begin**
  **if** *Identifying False Positives* **then**
    Potential Divergent Inds = **where**(M = Feature Minority States)
  **end**
  **if** *Identifying False Negatives* **then**
    Potential Divergent Inds = **where**(M = Feature Majority States)
  **end**
  Observed Divergences = EPM[Potential Divergent Inds]
  Transform Observed Divergences into z-scores
  Fit z-scores to a half normal distribution
  Identify Statistically Significant Divergent points as determined by CI
**end**

**Figure SI 10. Pseudocode for identifying statistically significant divergent data points.** Given $M'$ and its corresponding $EPM$, the pseudocode describes how a set of statistically significant divergent data points are identified, based on a given confidence interval ($CI$).

#### SI 6 ESFW

Here we present our algorithm Entropy Sorting Feature Weighting (ESFW). Using the divergent data points suggested by FFAVES and the rationale from ES to calculate $EPM_{FN}$ matrices, ESFW assigns a weight to each feature in the dataset. Higher weights correspond to those pairs of features that have stronger correlation. Conversely, low weights indicate features with weak co-regulatory relationships, indicating that the distribution of their minority states is essentially random compared to other features in the data.

We provide pseudocode for ESFW Fig. SI 11. The first step is to take the discrete state matrix ($M$) used as the input for FFAVES and create an augmented state matrix ($M'$), by switching the states of the indices that FFAVES suggested as significantly divergent. The next step is to intentionally add FN data points in the minority states of each feature in $M'$. Having started with the converged state matrix $M'$, we can be confident that the majority of divergence observed by intentionally adding FNs is caused by the controlled introduction of the FNs rather than unknown factors. Hence, when we calculate $EPM_{FN}$ and test whether relationships between dependent features gain significant divergence due to FNs, we can directly relate the observed positive EPs to features with strong dependent relationships. To ensure that the observed positive EPs are balanced, the same fraction of minority states, $FN\_Fraction$, are switched to the majority state for each feature. We set the default $FN\_Fraction = 0.1$, such that 10% of the minority state samples for each feature are randomly switched to the majority state. This default value was found empirically to produce good results over multiple data sets, and relatively stable when varied from 0.05-0.3 (data not shown).

After intentionally introducing FN data points to $M'$, the resulting $EPM_{FN}$ will be comparable to the example in Fig. SI 9B. To generate feature weights, we take the column means (columns correspond to features) for all points in $EPM_{FN}$ that have values greater than 0. We inspect only $EPM_{FN}$ values greater than 0 because these represent cases where introducing FNs has weakened feature dependencies through divergence. We take the column means of these $EPM_{FN}$ values rather than the columns sums, to allow for the fact that features may be members of different sized correlated networks. For example, in scRNA-seq data, mRNA expression relating to GRNs that control one aspect of a cell's function/identity, e.g. its lineage, may be significantly smaller/larger than the GRN that identifies its lineage subtype. The introduction of FNs ($FN\_Fraction$) to random features will lead to zero or low average $EPM_{FN}$ values. This is exemplified in Fig. SI 9B, where the randomly expressed features in the synthetic data display very low divergence values throughout the region of the data that they encompass.

Each of the column means obtained from $EPM_{FN}$ is the weight for each feature in $M$. We repeat this process multiple times and the $Iteration\_Weights$ for each feature are saved. Multiple iterations are carried out to ensure the search space for introducing random FNs is well covered. The number of iterations undertaken is defined by the $Iterations$ variable, with a default of 5. The final variable, $Feature\_Weights$, is a vector of length $j$ and is the mean weight of each feature from $Iteration\_Weights$. Having obtained a vector of weights representing the importance of each feature in a given data set, we are then able to subset the data down to highly informative features, as demonstrated in the main text.

**Algorithm 3:** ESFW

**Input** : $M$, $FFAVES\_Divergent\_Indices$, $FN\_Fraction$, $Iterations$
**Output** : $Feature\_Weights$
**begin**
  **for** *i = 1* **to** *Iterations* **do**
    $M' = M$ # Always start from initial discretised matrix
    $M' = (M'[FFAVES\_Divergent\_Indices] \times 1) + 1$ # Switch states
    **for** *Each feature in M'* **do**
      Randomly switch $FN\_Fraction$ of minority states to majority states
    **end**
    Calculate $EPM_{FN}$
    **for** *Each feature in M'* **do**
      $Iteration\_Weights$ = Mean of $EPM_{FN}$ values greater than 0
    **end**
    Store vector of $Iteration\_Weights$
  **end**
  **for** *Each feature in M'* **do**
    $Feature\_Weights$ = Mean of $Iteration\_Weights$
  **end**
**end**

**Figure SI 11. Pseudocode for estimating feature importance weights.** Given the initial discrete data matrix ($M$) and the FP/FN data points suggested by FFAVES ($FFAVES\_Divergent\_Indices$), the pseudocode describes the process for estimating a feature importance weight for each feature in $M$. $FN\_Fraction$ describes the fraction of minority states in each feature intentionally converted to majority states (default = 0.1). $Iterations$ defines how many times the feature weights will be estimated before the average is taken (default = 5).

#### SI 7 ES Minority State Cardinality Sensitivity

In Fig. 4C and D and Fig. S6 we quantify and visualise the performance of FFAVES in identifying FNs. A closer look at the ground truth FNs we fail to be identify suggests that the majority occur in genes that are active in very few cells (Fig. S6d). This demonstrates one of the limitations of FFAVES. We hypothesise that as the number of samples displaying the minority state of a feature decreases, the sensitivity of that feature to FN drop outs increases. We can demonstrate this empirically with an idealised simulation.

In Fig. SI 12 we set up one round of our proposed simulation and visualise the output after repeating the simulation for different parameters. For each simulation, we start with two perfectly overlapping features. We then randomly introduce a fixed fraction of FNs to each feature. We then use ES hypothesis testing to calculate the error potentials ($EP$) of the divergent points between the features. After repeating this multiple times for different minority state feature cardinalities and for different fractions of FNs, we formulate a heatmap of the average observed $EP$ for each ideal pair of features (Fig. SI 12). From this heatmap we can see that as the minority state cardinality of a feature decreases, the average observed $EP$ switches from positive to negative. Recall that a negative $EP$ value indicates that there is no longer any evidence that the observed divergence was due to the introduction of erroneous observations. Hence, this agrees with our hypothesis that for a fixed rate of introduced FN error, the smaller the minority state cardinality of the feature, the more susceptible it is to having its co-regulatory dependencies disrupted beyond a point that ES can quantitatively identify the error introduced.

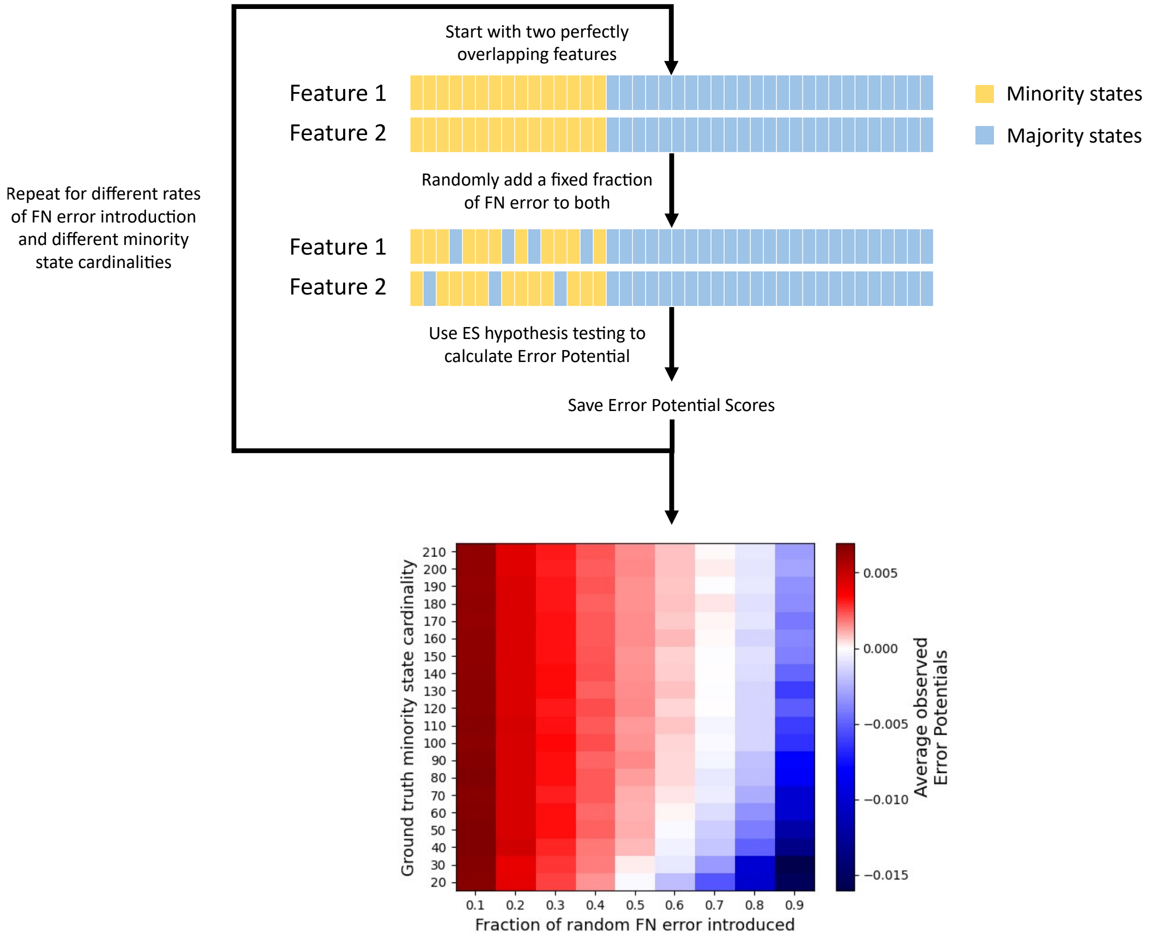


**Figure SI 12.** **Feature sensitivity to FN drop outs increases as minority state cardinality decreases.** Here we use a simple experiment to demonstrate that when the ground truth of a feature has too few samples displaying its minority state, the feature can become overly sensitive to FN drop outs. That is, there is a point where FN drop outs can no longer be detected by ES hypothesis testing. **(Top)** Two perfectly dependent features are created and a fixed fraction of samples are randomly forced into the majority state (if not already in the majority state). The $EP$ of the divergent points is then calculated. **(Bottom)** This process was carried out for a system with 1000 samples per feature, while varying the fraction of FN error to be introduced from 0.1-0.9 ($x$-axis), and the size of the ground truth minority state cardinalities from 20-210 ($y$-axis). Each simulation was carried out 100 times for each condition and the average $EP$ for each condition is displayed in the heatmap.

#### SI 8 Synthetic Dataset Details

The synthetic data set that we derive is designed to reflect the major properties of scRNA-seq data while allowing us to have a well defined ground truth to interrogate. It was created by combining nine different properties that are typically observed in scRNA-seq data sets to create a synthetic data set that reasonably represents the challenges of real data. As such, it has no underlying model or simulation of gene regulation. The benefit of using a relatively simple methodology is that we can be precise about the ground truth of the data before we intentionally add error to test ES. This subsequently allows us to confidently quantify how well any particular method performs when trying to recapitulate the ground truth properties of the data. The nine properties of the synthetic data are described below ((ii)-(viii) are highlighted in Fig. SI 13A).

(i) **Continuous gene expression**: Whenever it is deemed that a cell should express a given gene, its expression value is sampled from a $N(5,1)$ distribution.

(ii) **Five synthetic cell types**: We initiate SD1 by creating five distinct 'cell types', which are each sets of 200 cells. Within each cell type there is a hierarchical structure of gene expression, such that the number of cells that a set of genes is expressed in decreases from 200 to 20 in increments of 10. That is, one set of genes is expressed in all 200 cells of the given cell type, followed by a set of genes expressed in 190 of the cells, and so on, until a final set of genes is only expressed in 20 of the cells. The size of each subset of genes is randomly sampled from a discrete uniform distribution with values between 5 and 15 genes. In this way, each cell type is identified by a set of tightly regulated genes. Simultaneously, within each cell type there contains easily interpretable heterogeneity through the sequential gradient of cell type specific gene expression.

(iii) **Multi modal gene activity**: A simple example of a set of 50 genes that are defined to be active in 2 of the cell types. However, in one of cell types the average expression is twice as high than in the other.

(iv) **Structured genes**: Of the 1519 genes in the synthetic data, 969 are considered to be the 'structured' genes, which are tightly regulated within the 5 cell types. The process of generating these tightly regulated modules of genes is outlined in item (ii).

(v) **Randomly expressed genes**: One of the inherent challenges of scRNA-seq is filtering out those genes that are poorly structured with regards to cell type or cell function (Ramskö Ld et al. 2009). In our synthetic data, we introduce a subset of 500 genes with random expression throughout all 5 cell types. For each gene we randomly select the proportion of cells that will express the gene from a $N(0.3,0.3)$ distribution, but only accept values between 0 and 1. That proportion of cells are randomly selected and their expression value is sampled from a $N(5,1)$.

(vi) **Leaky gene expression**: Each of the 5 cell types represents an idealised situation where genes are tightly regulated to define a cell identity. We introduce leaky gene expression to represent when genes are detected as having non-zero expression, but such gene activity is out of context according to the structure in the rest of the data. In reality this could occur due to poor gene expression control in the cell, or the cell transitioning from one state to another. To represent this in our synthetic data, we identified the number of cells where active gene expression was detected for each gene. We then randomly pick a value between 0.02 and 0.1 to choose what percentage of leaky gene expression to add to the data for that feature (2-10%). The number of leaky expression data points to be added to a particular gene is equal to the number of cells found to have the gene active, multiplied by the sampled value between 0.02 and 0.1. Leaky gene expression points are added by randomly switching that proportion of samples in the data from zero to non-zero expression values for each gene.

(vii) **Multiple cells per sample**: To incorporate the reality of a small number of samples with multiple cells (JD 2018) we added 200 cells (20% of the initial 1000 cells) made by randomly combining cells from the 5 synthetic cell types. For each multiplexed sample, the number of cells to be combined was determined by sampling from a Poisson distribution with mean = 1, but continuing to sample until a value greater than 1 was obtained. A Poisson distribution with mean = 1 is used in accordance with the methodology used when diluting a pool of cells to be sequenced (JD 2018). Having determine the number of cells in the multiplexed sample, that number of cells are randomly taken from the 5 synthetic cells types and summed together.

(viii) **Random up/down-regulation between cell types**: ES was developed with the aim of identifying functional rather than simply correlative relationships between features. To demonstrate this, we added a set of 50 genes that are randomly up-regulated in just two of the synthetic cell types. Although there is a higher frequency of observing these 50 genes in these two cell types, the likelihood of observing the active gene states overlapping with one another for the 50 features is still relatively low. To create the 50 genes, for each we randomly picked a fraction of the samples (2-10%) in the two

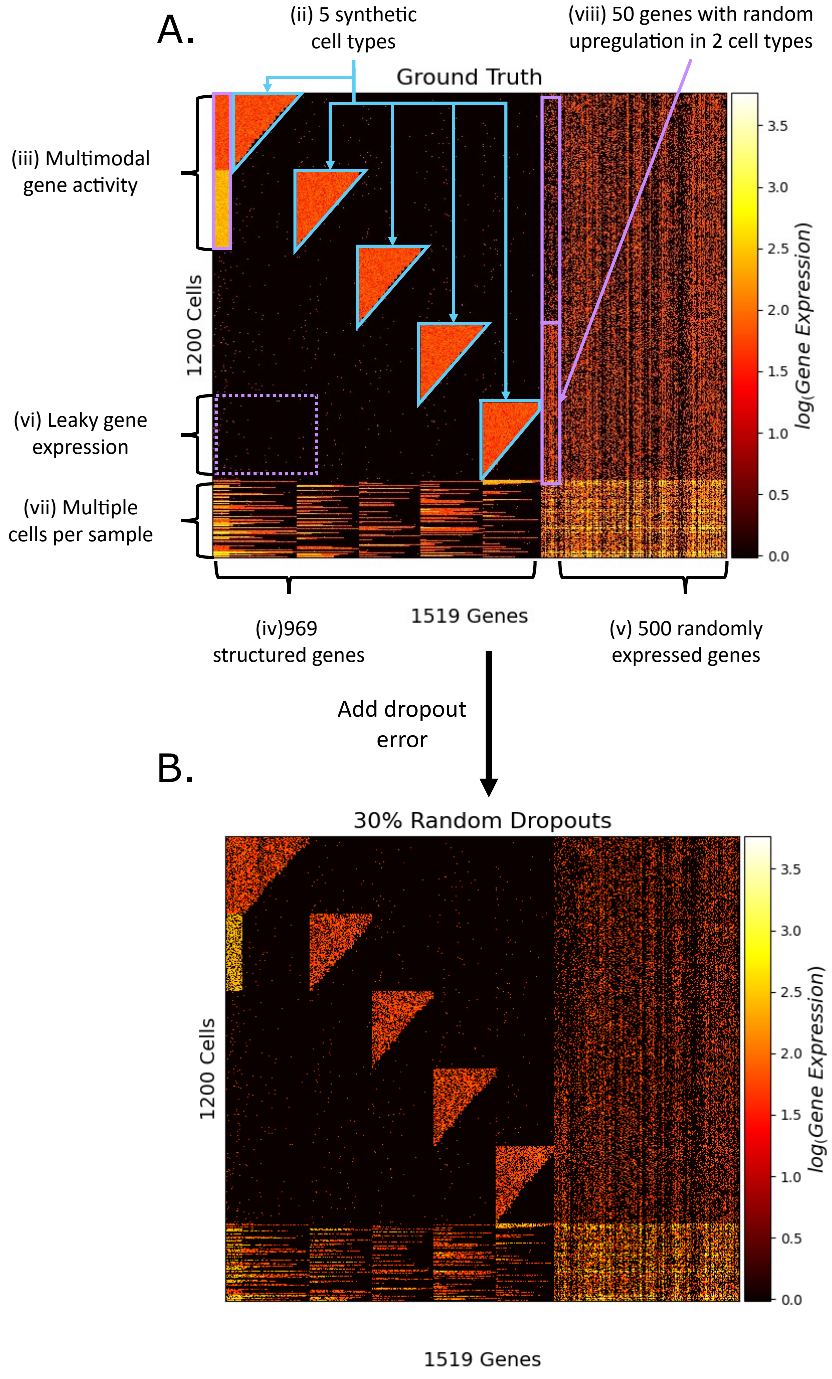


**Figure SI 13.** **Synthetic data properties A.** Visualisation of the ground truth synthetic gene expression matrix. Each of the defined properties of the data marked as (ii)-(viii) are described in Section SI 8. **B.** Visualisation of SD1 after 30% of the ground truth data points are converted to FNs (to mimic technical drop outs).

chosen cell types, and switched that fraction of cells to show the gene as active. For the remaining three cell types, we halved the fraction of cells showing the gene as expressed in the “up-regulated" cells and switched this smaller faction of cells to show the gene as active in the “down-regulated" cells. For example, if a gene would be active in 8% of the cells in the two cell types where it was up-regulated, the same gene would only display as active in 4% of the cells in the remaining three synthetic cell types.

(ix) **Batch effects**: Batch effects can be a significant confounding factor in the analysis and interpretation of scRNA-seq data. Having generated our synthetic dataset by combining each of the data properties in (i)-(viii), we add batch effects as follows. We create two batches of cells by grouping together odd and even numbered cells. For each gene we randomly select one of the batches and add a random drop out bias to all of the cells in that batch. This may be thought of as mimicking technical batch effects, where there is random variability in the capture efficiency of genes between the two batches. To add the drop out bias, we identify all cells in the batch in which the given gene is active. We take the absolute value after sampling from a $N(0, 0.4)$ distribution to determine the fraction of active cells contained in the batch that will have their expression value set to 0 for the given gene. Finally, we randomly introduce the dropouts into the gene by switching the sampled fraction of cells from non-zero to zero values.

The culmination of each of these 9 synthetic data properties is shown in Fig SI 13A. This represents our known ground truth. We then add a final layer of complexity to the data by introducing a significant portion of FN random dropouts (Fig SI 13B). This is achieved by randomly selecting 30% of all data points and switching them to a value of 0 if they are not already 0. This noisy representation of the synthetic data serves as the starting point for us to try to recover the different substructures within the data, such as FNs/FPs and modules of highly co-regulated gene.

#### SI 9 Human embryo immunostaining experimental procedures

Supernumerary frozen blastocysts (E5 and E6) were thawed and cultured in N2B27 medium under mineral oil until reaching the desired stage of development from E5 to E7. Embryonic stage was assessed based on thinning of the zona pellucida and blastocoele expansion.

The zona pellucida of D5 and D6 blastocysts was removed using acid Tyrode’s solution before fixation with 4% PFA in PBS for 15 minutes at room temperature. Embryos were rinsed in PBS containing 3mg/ml polyvinylpyrrolidone (PBS/PVP), permeabilised using 0.25% Triton X-100 in PBS/PVP for 30 minutes and blocked in blocking buffer comprising PBS supplemented with 0.1% BSA, 0.01% Tween20 and 2% donkey serum for 2 hours at room temperature. Primary and secondary antibodies were diluted in blocking buffer (Table S1). Embryos were incubated in primary antibody solution overnight at 4 degrees and rinsed three times for >15 minutes each in blocking buffer before incubation in secondary antibody solution for 1-2 hours at room temperature in the dark. Embryos were rinsed in blocking buffer and imaged through a Poly-D-Lysin coated Mattek dish (P356-0-14) whilst submerged in blocking buffer. Embryos were imaged in a Leica Stellaris Confocal microscope and image analysis was performed using FIJI.
